# A new theoretical framework jointly explains behavioral and neural variability across subjects performing flexible decision-making

**DOI:** 10.1101/2022.11.28.518207

**Authors:** Marino Pagan, Vincent D Tang, Mikio C. Aoi, Jonathan W. Pillow, Valerio Mante, David Sussillo, Carlos D. Brody

## Abstract

The ability to flexibly switch our response to external stimuli according to contextual information is critical for successful interactions with a complex world. Context-dependent computations are necessary across many domains^1–3^, yet their neural implementations remain poorly understood. Here we developed a novel behavioral task in rats to study context-dependent selection and accumulation of evidence for decision-making^4–6^. Under assumptions supported by both monkey and rat data, we first show mathematically that a network can solve this problem through a combination of three defined components. These components can be identified and tested directly with experimental data. We further show that existing electrophysiological and modeling data are compatible with the full variety of possible combinations of these components, suggesting that different individuals could use different component combinations. To study variability across individual subjects, we developed automated, high-throughput methods to train rats on our task, and we trained many subjects on it. Consistent with theoretical predictions, neural and behavioral analyses revealed substantial heterogeneity across rats, despite uniformly good task performance. Our theory further predicts a specific link between behavioral and neural signatures, which was robustly supported in the data. In summary, our results provide a new experimentally-supported theoretical framework to analyze individual variability in biological and artificial systems performing flexible decision-making tasks, they open the door to cellular-resolution studies of individual variability in higher cognition, and they provide insights into neural mechanisms of context-dependent computation more generally.

---

In our daily lives, we are often required to use context or top-down goals to select relevant information from within a sensory stream, ignore irrelevant information, and guide further action. For example, if we hear our name called in a crowded room and our goal is to turn towards the caller, regardless of their identity, information about *location* of the sound will drive our actions; but if we wish to respond based on the identity of the caller, the *frequencies*, in the very same sound, will be most important for driving our actions. As with other types of decisions, when the evidence for or against different choices is noisy or uncertain, accumulation of many observations over time is an important strategy for reducing noise^1,4,8,9^. What neural mechanisms underlie our ability to flexibly accumulate evidence about external stimuli, and to switch our response according to contextual information?

Here we developed a series of new experimental and computational techniques to address this question. First, we developed a novel behavioral pulse-based task in rats to study context-dependent selection and accumulation of evidence for decision-making. Delivering evidence in highly-random, yet precisely-known pulses provided us with high statistical power to precisely characterize the rats’ behavior and neural dynamics. Then, using an automated, high-throughput procedure, we trained many rats to solve the task, allowing us to uncover a surprising degree of variability in the behavior and neural dynamics across individuals, even when they were all well-trained, high performing, animals. Next, we developed a novel mathematical framework that defined the space of solutions for networks that can implement the required computation. The theoretical framework predicted that variability in position in that solution space, within and across individuals, should be the underlying variable that would jointly drive variability in behavior and neural responses-- implying that behavioral and neural variability should be tightly correlated. Our experimental data robustly confirmed this theoretical prediction. Finally, we developed techniques to engineer artificial recurrent neural networks across the full range of our theoretical solution space, and showed that gradient-descent methods, as typically used to train network models, lead to only one corner of the possible data-compatible solutions.

## Context-dependent accumulation of evidence in rats

To study the neural basis of context-dependent selection and accumulation of sensory evidence, we trained rats on a novel auditory task where, in alternating blocks of trials, subjects were cued to determine either the prevalent location (“LOC”) or the prevalent frequency (“FRQ”) of a sequence of randomly-timed auditory pulses (Fig. 1a). The relative rates of left vs. right and high vs. low pulses corresponded to the strength of the evidence about LOC and FRQ, respectively (Fig. 1b). These relative rates were chosen randomly and independently on each trial, and were used to generate a train of pulses that were maximally randomly-timed, i.e., Poisson-distributed. Correct performance requires selecting the relevant feature for a given context, accumulating the pulses of evidence for that feature over time, and ignoring the irrelevant feature. Many rats were trained to good performance on this task using an automated training procedure (Fig. 1c; training code available at https://github.com/Brody-Lab/flexible_decision_making_training), with most rats learning the task in a timespan between 2 and 5 months (Extended Data Fig. 2g). After training, rats associated the audio-visual cue presented at the beginning of each trial with the correct task context, and were able to switch between selected stimulus features within ∼4 trials of a new context block (Extended Data Fig. 1e). Our task structure was inspired by a previous visual task used with macaque monkeys^4^; major distinctions between the previous and current tasks included the species difference, the sensory modality difference, and the pulse-based nature of our task; this last will be key for the analyses performed below. Despite the important differences across tasks, attained performances were similar across the two species (Extended Data Fig. 1c,f). We reasoned that the highly random yet precisely known stimulus pulses, together with large numbers of trials and subjects, would provide us with statistical power to characterize both behavioral^10^ and neural responses.

**Figure 1.**
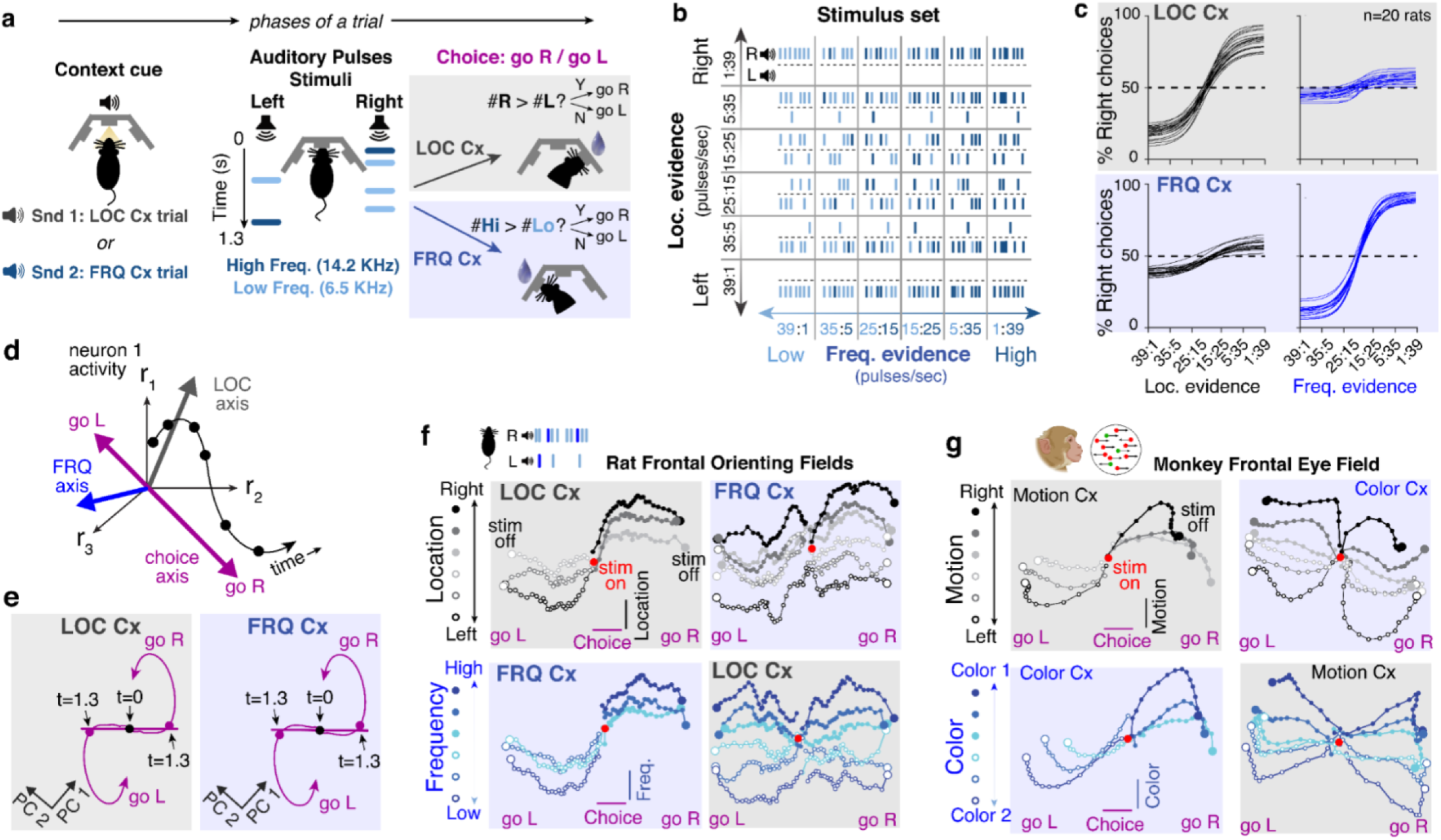
Rats can be trained to perform context-dependent evidence accumulation with behavioral performance and average neural dynamics similar to macaque monkeys. **a)** Task design. Each trial starts with a sound indicating current context, either location **(LOC)** or frequency **(FRQ)**. Rats are then presented with a 1.3 sec-long train of randomly-timed auditory pulses. Each pulse is played either from a Left or Right speaker, and is either Low-(6.5 KHz; light blue) or High-frequency (14 KHz; dark blue). In **LOC** trials, subjects are rewarded if they turn, at the end of the stimulus, towards the side that played the total greater number of pulses, ignoring pulse frequency; in **FRQ** trials, subjects should turn Right(Left) if there was a greater number of High(Low) pulses, ignoring pulse side location. Because the two features are independent, an identical stimulus can be associated with opposite responses in the two contexts. **b)** Schematic of the stimulus set used. The relative rates of left:right and high:low pulses correspond to the strength of the evidence about location and frequency, respectively. The relative rates were chosen randomly on each trial from the set shown on the horizontal and vertical axes (keeping the sum of the two rates fixed at 40 pulses/sec), and were then used to generate a Poisson-distributed pulse train. **c)** Psychometric curves showing good performance of 20 rats after training (n>120,000 trials for each rat): In the **LOC** context, location evidence strongly affects choices, while frequency evidence has only small impact; and in the **FRQ** context, frequency evidence strongly affects choices, while location evidence has limited impact. Curves are fits to a logistic function; see Methods. **d)** Illustration of analysis of population dynamics. The activity of many neurons at a given point in time corresponds to a point in high-dimensional “neural space”. As activity evolves, this traces out a trajectory over time. This population trajectory can be projected onto axes chosen to best encode momentary **LOC** or **FRQ** evidence, or to best predict the subject’s choice. In this figure, we consider analyses in which the traces and their projections are computed after averaging over trials within each block of panel 1b. **e)** The trajectory in neural space that best accounts for the choice-dependence of the neural activity, projected onto its first two principal components (PCs) (accounting for 81.3% of its variance; Between t=0 and t=1.3s the choice axis explains 73.3% of the variance; Methods). The choice trajectory was computed separately for each context, but the PC directions were computed in common for the two contexts. The “choice axis” was defined as the straight-line fit to the trace t=0 to t=1.3s; the angle between choice axes in the two contexts was negligible (average 1.6^°^, significantly different from 0 in only 1 out of 7 rats; see Extended Data Figure 5). **f)** Neural population trajectories from recordings in the Frontal Orienting Fields (FOF) of rats performing the task shown in panel a. Trajectories are projected onto the choice and **LOC** axes (top row), or onto the choice and **FRQ** axes (bottom row). Only correct trials are shown. Trajectories are further sorted, and color-coded, by the strength of **LOC** feature (top row}, or the strength of the **FRQ** feature (bottom row). **g)** Same analysis and conventions as in (f) for neural recordings from the Frontal Eye Fields (FEF) of macaque monkeys performing a visual task requiring context-dependent accumulation of either motion or color evidence features (Mante et al., 2013).

To compare neural dynamics in a decision-making region across monkeys and rats, we examined neural activity in the Frontal Orienting Fields (FOF) while rats performed our task. The FOF are a rat cortical region thought to be involved in decision-making for orienting choice responses^11,12^, and have been suggested as homologous or analogous to macaque Frontal Eye Fields (FEF)^11,13,14^, which are the cortical region recorded in the previous monkey task^4^. Consistent with a key role for the FOF in our task, bilateral optogenetic silencing of rat FOF demonstrated that it is required for accurate performance of the task (Extended Data Fig. 4; n=3 rats). We implanted tetrodes into the FOF and into another frontal region, the medial prefrontal cortex (mPFC), and we recorded from n=3495 putative single neurons during n=199 sessions from n=7 rats while they performed the task of Fig. 1. As with previous reports in frontal cortices of both macaques and rodents, we found that task-related firing rates were highly heterogeneous across neurons. We then carried out the same analysis that had been applied to the (also heterogeneous) neurons recorded from monkey FEF^4,15^, and found strong qualitative similarities across the two species (compare Fig. 1f to Fig. 1g). The analysis, known as “targeted dimensionality reduction” (TDR) begins by describing neural population activity at a given moment in time as a point in “neural space”, where each axis represents the firing rate of one of the *N* recorded neurons. As activity evolves over the duration of a trial, a trajectory in *N*-dimensional neural space is traced out (Fig. 1d). Following ref. ^4^, neurons recorded separately in different sessions were combined into a single time-evolving *N-*dimensional neural vector. This “pseudo-population” activity was averaged across trials with a given generative pulse rate (i.e. within each of the 36 blocks in Fig. 1b), for each of the two contexts, and for each of the animal’s choices. These trajectories were projected onto the orthogonalized linear subspaces that best predicted the subject’s choice, or momentary location evidence, or momentary frequency evidence (illustrated as different axes in Fig. 1d). We found that trajectories for different evidence strengths were clearly separated along each sensory feature’s axis (see separation of traces along the vertical axes of the panels of Fig. 1f; only correct trials are shown). This was true regardless of whether the feature was relevant or irrelevant (compare vertical separation for left versus right columns in Fig. 1f). A similar observation in the monkey data (Fig. 1g) previously led to the conclusion that irrelevant feature information was not gated out from reaching frontal cortex^4^; the same conclusion applies to our rat data. In the next section of the manuscript, we present a theoretical analysis that applies equally to this scenario (i.e. no gating of irrelevant information before reaching frontal cortex), as well as to alternative mechanisms relying on early gating, an aspect we return to in the Discussion. Overall, the striking qualitative similarity between the rat (Fig. 1f) and monkey (Fig. 1g) traces suggests that the underlying neural mechanisms in the two species may be similar enough that an active exchange of ideas between studies in the two species will be very fruitful.

Using model-based TDR analysis^15^, we found the 2-dimensional subspace that best accounts for the contribution of the animal’s choice to the neural activity. We then projected the kernel-based estimates of “go Right” and ”go Left” trajectories (which are noise-reduced versions of the raw trajectories (see Extended Data Fig. 5h) onto it (Fig. 1e). During the stimulus presentation (t=0 to t=1.3s, a period in which subjects must accumulate sensory evidence), this choice-related information in firing rates evolved along an essentially one-dimensional straight line in neural space, only later curving into a second dimension (see Ext. Data Fig. 5 for per-animal analysis). This is consistent with previous findings, with the initial linear phase having been suggested as corresponding to gradual evidence accumulation, while the subsequent rotation may correspond to formation of a motor plan^16,17^, perhaps after commitment to a decision^15,18^. We will focus on evidence accumulation during this linear phase, while the decision is being formed, and will refer to the corresponding line in neural space as the “choice axis”: the animal’s upcoming choice can be predicted from position on this axis. Crucially, both correct and incorrect trials are used for this analysis, allowing to separate this choice-predictive signal from responses to sensory stimuli. In a final similarity with the monkey data, we found that the choice axes, estimated separately for each of the two contexts, were essentially parallel (average angle between contexts = 1.6º ; not significantly different from 0 (p>0.1) for 6 out of 7 animals; Methods).

Consequently, in the theoretical development below we will assume that the direction of the choice axis is the same in the two contexts. However, this simplifying assumption can be relaxed, as will be addressed in the discussion and as we detail in Extended Data Fig. 10.

## The space of possible solutions is spanned by three distinct components with different behavioral and anatomical implications

It has long been hypothesized that neural dynamics around the choice axis are well approximated by a “line attractor”^7,19^, i.e. that the choice axis is formed by a closely-packed sequence of stable points. This follows from the idea that the position of the system on the choice axis corresponds to net accumulated evidence towards Right vs Left choice; in temporal gaps between pulses of evidence, an accumulator must be able to stably maintain accumulated values, and thus position anywhere along this axis should be a stable point. We now develop theoretical implications of this computation-through-dynamics^7^ line attractor hypothesis, which lead to a new description of the space of possible network solutions consistent with the hypothesis, and to new experimental predictions that we find to be robustly supported by the data.

A key implication of the line attractor hypothesis, which follows from linearized approximations of the system’s dynamics, is that a sensory stimulus pulse that perturbs the system along direction **i** has a net effect on position along the choice axis given by^4,19^ the dot product of that input vector **i** and another vector known as the “selection vector” **s**. That is, the change in choice axis position = **s•i** ; see “Box 1”. Thus, in the linear dynamics approximation, and under the line attractor hypothesis, the simple dot product **s•i** summarizes the result of the interaction of local recurrent dynamics (represented by **s**) with a pulse of external input (**i**).

### BOX 1

**Dynamics around line attractors**

Linearized dynamics around a fixed point in neural space can be represented by equation (1), where **M** is a matrix and **r** is a vector representing the system’s position in neural space relative to the fixed point. The eigencoordinates **e**, defined in equation (2), where the columns of **V** are the eigenvectors of **M**, can also be used to describe these dynamics. The advantage of eigencoordinates is that each element *j* of the vector **e** evolves over time independently of the others, following *e*_*j*_*(t) = e*_*j*_*(t=0)* exp(*λ*_*j*_ *t)*, where *λ*_*j*_ is the eigenvalue corresponding to the *j*^*th*^ eigenvector.

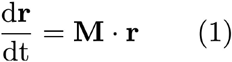

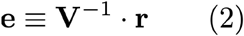

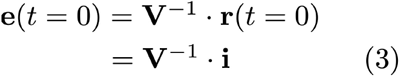

For a line attractor, one eigenvalue (by convention the one with index *j* = 0) has value 0 (i.e., *λ*_*0*_ = 0) and consequently *e*_*0*_*(t)* = const. = *e*_*0*_*(t = 0)*. All other eigenvalues have real part < 0, implying all eigencoordinates *j > 0* decay to zero over time, as the system state relaxes back onto the line attractor. Thus, if an external input pulse **i** perturbs the system off the line attractor onto position **r***(t = 0)* = **i**, it follows that, after the transients in which eigencoordinates *j > 0* decay to zero, the new position on the line attractor, relative to the starting fixed point, will be given by *e*_*0*_*(t)* = *e*_*0*_*(t = 0)*, since this will be the only nonzero eigencoordinate. Equation (3), for the eigencoordinates of the initial position, tells us that the zero^th^ eigencoordinate of the initial position, *e*_*0*_*(t = 0)*, will be the dot product of the the top row of **V**, which we label as the “selection vector” **s**, and the input vector **i**. (refs. ^4,19^)

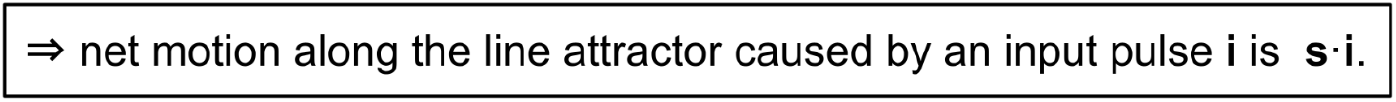

It follows that for a pulse of evidence to have a greater effect on choice in the context in which it is relevant than when it is irrelevant, **s•i** must be greater in the relevant than in the irrelevant context. The recurrent dynamics in the decision-making region could be different in the two contexts; similarly, context-dependent modulation of early sensory responses^20–24^ could lead the direction **i** along which a pulse of a given feature perturbs the system to be different in the two contexts. Thus, indicating relevant vs irrelevant context with a subscript (**s**_**REL**_ vs **s**_**IRR**_, and **i**_**REL**_ vs **i**_**IRR**_), the general condition for a given feature’s input pulse to have greater effect on choice when relevant vs irrelevant is:

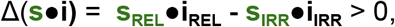

with Δ indicating difference across contexts. For each of the features being considered (in our experiments, LOC and FRQ), this difference Δ(**s•i)** can be rewritten as the sum of three components (Fig. 2c).

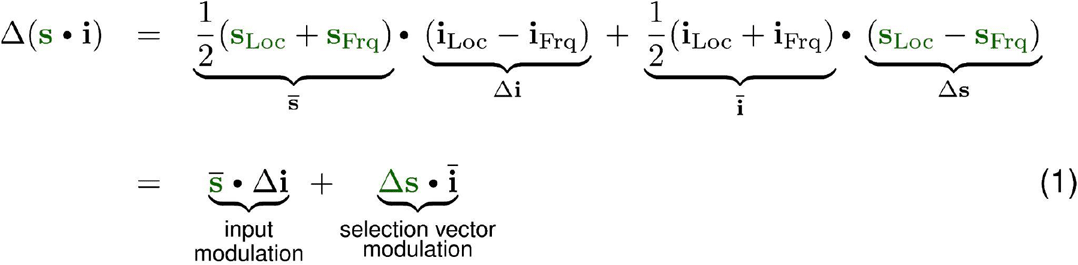

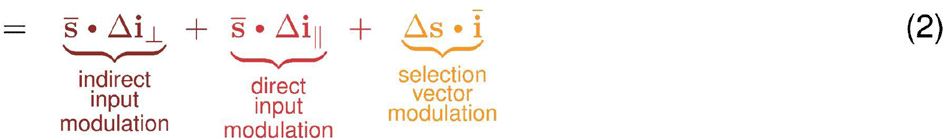

where the overbar symbol represents the average over the two contexts, Δ represents difference between the two contexts, and Δ**i**_⊥_ and Δ**i**_**‖**_ represent the component of Δ**i** that is orthogonal and parallel to the choice axis, respectively. For any given feature (here, either LOC or FRQ), and for any given network that solves the task (and thus has Δ(**s•i)** > 0), the percentage that each of the components contributes to the total Δ(**s•i)** can be visualized in terms of distances from the vertices of a triangle, i.e., a point in barycentric coordinates (Fig. 2g). We emphasize that all positions on the triangle have Δ(**s•i)** > 0 and thus all describe solutions; what the different positions describe is variations across networks that embody different solutions for the task. This will be a key aspect to understanding variability across different individuals that all solve the task. The first component in equation (2), “indirect input modulation” is what follows if the difference across contexts is a change in the input vector **i**, with the change *orthogonal* to the line attractor; the second, “direct input modulation,” follows from change in the input **i** that is *parallel* to the line attractor; and the third, “selection vector modulation,” follows from a change in the selection vector **s** that represents the recurrent dynamics in the decision-making region itself. The manner in which each of the components of equation (2) lead to a greater change in line attractor position for the relevant than the irrelevant context is illustrated in Figs. 2d,e,f.

**Figure 2.**
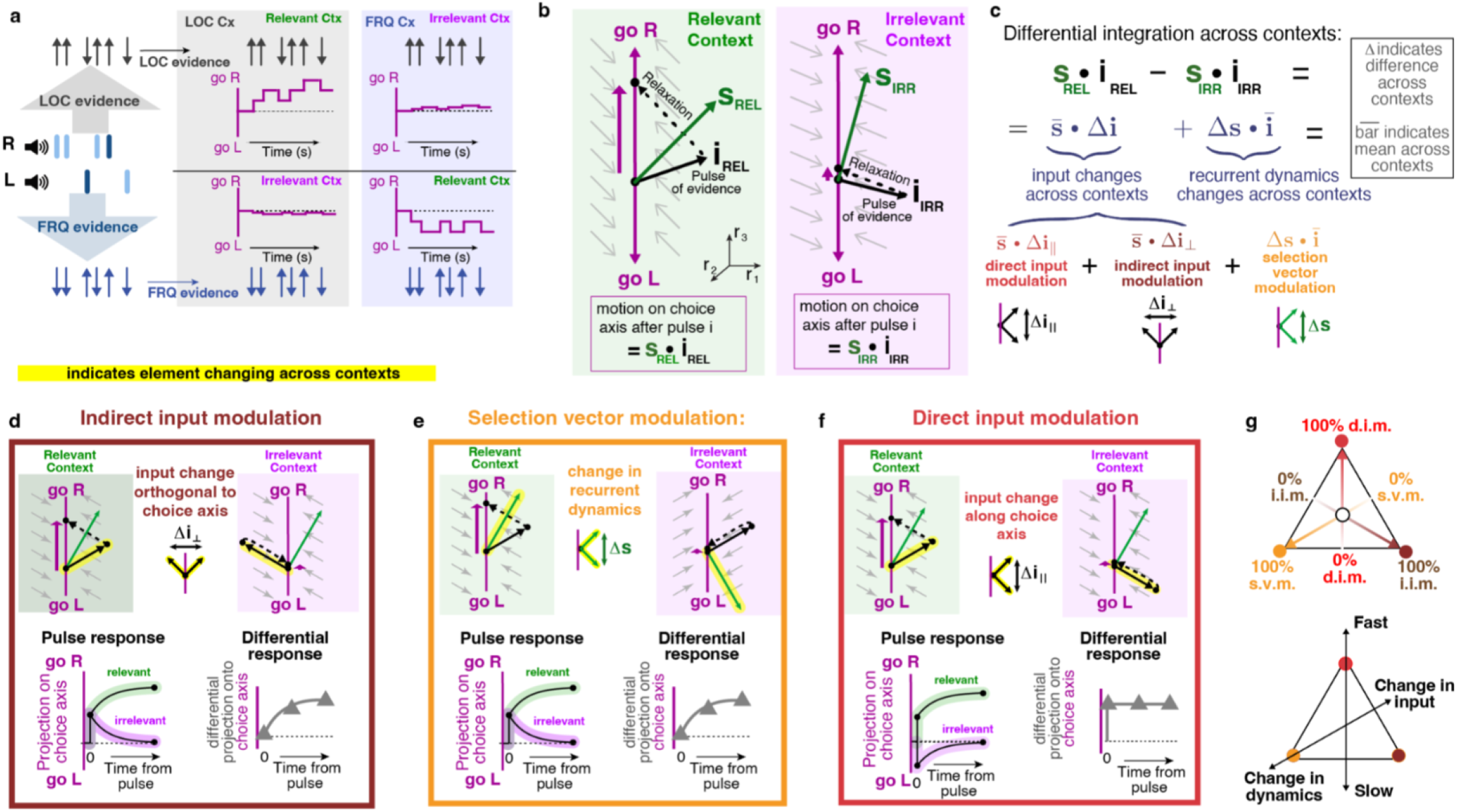
Context-dependent evidence selection can be dissected into three distinct components, each with different biological implications, **a)** Conceptual illustration of task solutions. A given pulse train can be thought of as providing a train of Go-Left (|) versus Go-Right (f) pulses of LOC evidence (top), and independently, of FRQ evidence (bottom). All solutions require pulses of relevant evidence to have an effect on the system’s position along the choice axis, thus driving choices, while irrelevant evidence should have little effect on the choice axis, **b)** Assuming that the choice axis is a line attractor, the net effect on position along the choice axis, from a single pulse of evidence of a given feature presented in its relevant context, is given by the dot product of the “selection vector” s_REL_ that represents the recurrent dynamics, with the input vector **i**_**REL**_.that represents the initial deflection caused by the evidence pulse (Main text Box 1). Similarly, in the irrelevant context, the net effect on choice axis position is s_|R•_**i**_|**RR**_. Following Fig. 1 e, the choice axis is taken to be parallel across contexts, **c)** Solving the task means that relevant evidence should have a bigger effect on choices than irrelevant evidence, i.e., s_REL •_i_**REL**_ • SI_RR•_**i**|_**RR**_> 0. This can be rewritten as the sum of three components that together span the space of possible solutions. **d,e,f)** Illustration of how each of the three components can lead to greater motion along the choice axis in the relevant than in the irrelevant contexts. Same conventions in all three panels, **d)** “Indirect input modulation” is a change in input, across the two contexts, that is orthogonal to the choice axis. Bottom left: The projection onto the choice axis is initially the same in the two contexts, differing only after the relaxation dynamics. Bottom right: The “differential pulse response” (difference across contexts in the projection onto the choice axis of the response to an evidence pulse) grows gradually from zero, **e)** “Selection vector modulation” describes changes across contexts in the recurrent dynamics. As with indirect input modulation, the differential pulse response is initially zero and grows only after the relaxation dynamics, **f)** “Direct input modulation” is a change in the input vector that is parallel to the choice axis. In contrast to d) and e), the differential pulse response is non-zero immediately upon presentation of the input pulse, **g)** Top: All recurrent networks that solve the task, after linearizing, can be expressed as a weighted sum of the three components, and therefore mapped to a point on a triangle with barycentric coordinates. Bottom: A network’s position on this barycentric coordinate space has important biological implications: the vertical axis on the triangle quantifies how quickly the differential pulse response diverges from zero, and thus quantifies the speed of context-dependent effects, with neural and behavioral implications explored in subsequent figures. A second axis (shown as oblique line) captures the extent to which contextual computation relies on context-dependent inputs to the decision-making network, versus context-dependent recurrent dynamics of the decision-making network.

Different positions on the triangle of Fig. 2g are not merely distinct mathematically; they have different, and important, biological implications: First, where a network solution lies along the “Change in inputs” versus “Change in dynamics” tilted axis in Fig. 2g has significant anatomical implications. For networks at the “Change in dynamics” corner, the anatomical locus of context-dependence must be in decision-making regions, for it is the recurrent dynamics of these regions that differ across contexts. In contrast, for networks at the “Change in inputs” end of the axis, the anatomical locus of context-dependence could be outside decision-making regions— for example, it could lie in modulation of responses in sensory regions^20–24^, or in modulation of the pathways from sensory to decision-making regions^25^. Second, where a network solution lies along the vertical “Fast” versus “Slow” axis in Fig. 2g has both neural and behavioral implications. We describe the neural implications first. Networks at the “Slow” end of the axis have 0% direct input modulation (d.i.m.), i.e., they are all mixtures of indirect input modulation (i.i.m.) and selection vector modulation (s.v.m.). For both i.i.m. and s.v.m, the projection of the system’s position onto the choice axis immediately after a pulse of evidence is the same for the two contexts, and the difference across contexts develops only gradually (Fig. 2d,e, “differential pulse response” in bottom panels). In contrast, networks at the “Fast” end of the axis are 100% direct input modulation (d.i.m.), and for these a difference across contexts in the projection onto the choice axis is immediate (Fig. 2f, bottom panels). It is in this sense that neural context-dependence effects on the choice axis are fast at the d.i.m. end of the axis, and slow at the base of the axis (s.v.m / i.i.m). If behavioral choices are driven by the system’s position on the choice axis, it follows that solution diversity on this axis will produce consequent behavioral diversity; we examine this idea further in Fig 5.

Two parenthetical remarks follow from the algebraic rewriting in equation (2). First, early gating out of irrelevant information (**i**_**IRR**_=0) is a special case within this framework, and can be either direct input modulation (example 1 in Extended Discussion) or indirect input modulation (example 2 in Extended Discussion). Second, the direction of the line attractor enters the rewriting only in the step from equation (1) to equation (2), when distinguishing indirect versus direct input modulation. This is because this step describes Δ**i** as the sum of a component orthogonal and a component parallel to a particular reference direction that is fixed across the two contexts; here, this reference is the direction of the line attractor. We will focus on the case where the line attractor direction is the same in the two contexts both for simplicity and because it is what we found in our rat data (Fig. 1) as well as what was found in the monkey data of Mante et al. 2013. But equation (2) can be extended to the case of line attractors that are not parallel across the two contexts^16^ (see also Discussion and Extended Data Figure 10).

## Pulse-based analyses can distinguish different network solutions that all perform the task, and reveal variability within and across individuals

Artificial model networks can be used to illustrate approaches to solving the task. To find networks with many individual heterogeneous units, as observed in the experimental data (see e.g. Extended Data Fig. 4), Mante et al. (2013) trained recurrent neural networks (RNNs) to perform the task. Using the analyses of Fig 1d-g, they observed important similarities between the neural trajectories in the experimental data and in the trained RNNs. Upon analyzing the RNN’s linearized dynamics, they found that the trained RNNs solved the task using selection vector modulation. This prompted their influential suggestion of selection vector modulation as the leading candidate for how the brain implements context-dependent decision-making. What was unappreciated at the time was that the linearization they used (“activation space” linearization -- see “Linearizing RNN dynamics in firing rate space versus activation space” in Extended Discussion) precluded observing input vector modulation (whether direct or indirect) for the type of inputs used in their networks^26^. We therefore repeated their analysis, but now using a linearization (“firing rate space” linearization) that does permit observing input vector modulation in these RNNs^27^. Starting from randomly chosen initial network weights, we trained many RNNs to solve the task, analyzed their linearized dynamics, and using eqn. (2), plotted each RNN’s position in barycentric coordinates. The results with the new linearization at first sight confirmed the essence of Mante et al.’s 2013 conclusion, namely, that the trained RNN solutions are densest near the selection vector modulation corner at bottom left (Fig. 3a). However, the insight in eqn. (2), together with our choice of linearization in firing rate space, also allowed us to engineer RNNs that solve the task and lie at *any* chosen point within the barycentric coordinates (see Methods)-- that is, we are no longer constrained to exclusively examine the set of RNN solutions produced through training. Surprisingly, we found that selection vector modulation is not required to produce trajectories like those in Fig. 1f,g. Instead, network solutions at *any* point within the barycentric coordinates, not only those close to the selection vector modulation corner, produce traces that are qualitatively similar to the experimental data (see Fig. 3d and Extended Data Fig. 6). This suggests that analyses such as the one in Fig. 1f,g, which averages trials within each stimulus block in Fig. 1b, cannot readily distinguish between different solutions across the barycentric coordinates of Fig. 2g -- a space which, as described above, spans all possible solutions consistent with the choice axis being parallel across the two contexts.

**Figure 3.**
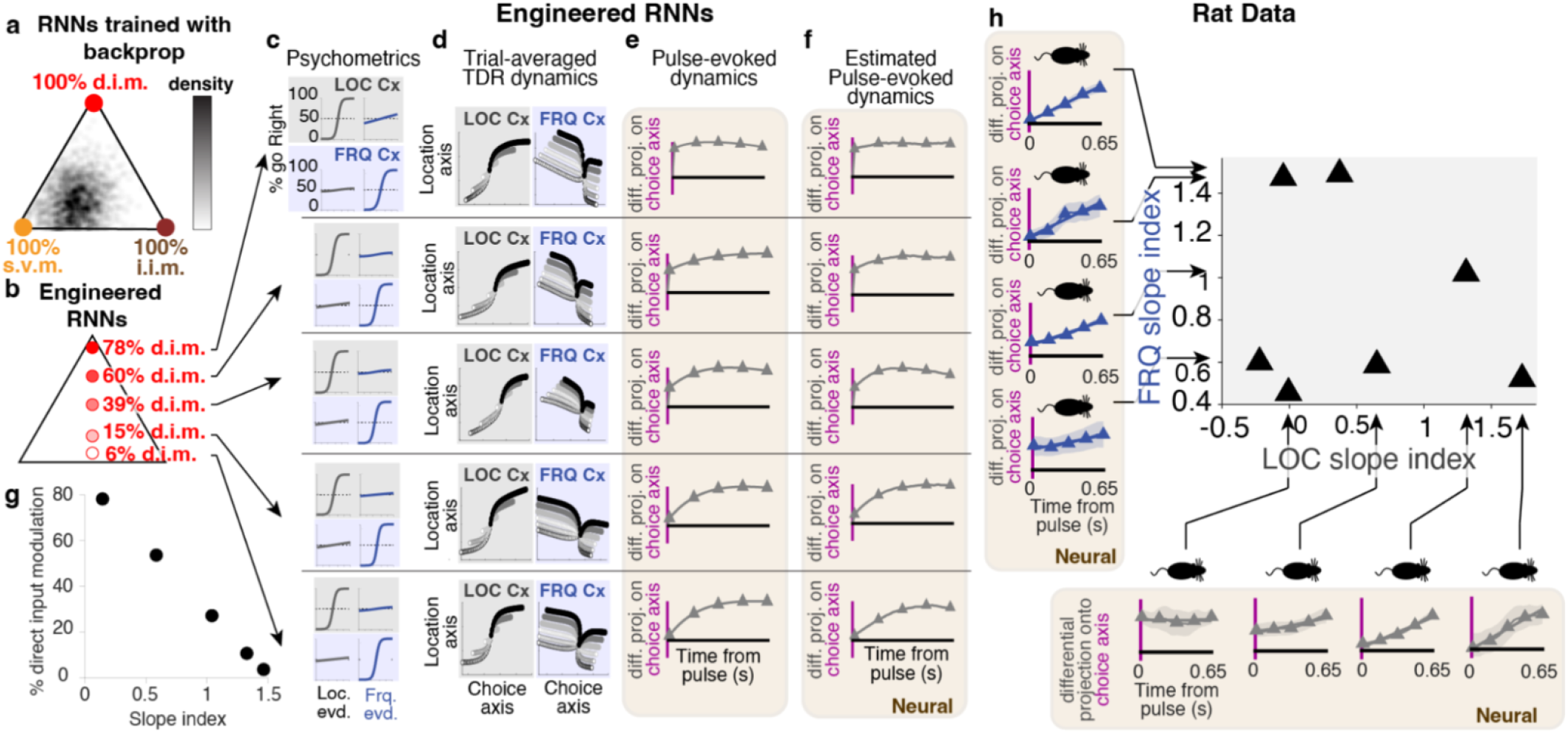
While trained recurrent neural networks (RNNs) explore only a subset of possible solutions, engineered RNNs can span the full space of solutions and produce heterogeneity in fast vs slow pulse-evoked dynamics that matches corresponding heterogeneity found in experimental data. **a)** Distribution of one thousand 100-unit RNNs trained to solve the task using backpropagation through time: After training starting from random initial points, the networks favored selection vector modulation (s.v.m.), as found in Mante et al. (2013). **b)** Instead of using backprop, RNNs can be engineered to lie anywhere in the space of solutions (Extended Data Fig. 6), including, as shown here, the vertical axis, from 0% direct input modulation (d.i.m.) to 100% d.i.m. **c,d,e,f)** Each row analyzes a single trained RNN, with different rows having different % d.i.m. percentage, as shown in b). c) Networks across the O to 100% d.i.m. axis perform the task with psychometric curves qualitatively similar to the experimental data (compare to **Fig, 1c**). **d)** All of the networks have neural activity that produces TDR traces qualitatively similar to the experimental data (compare to **Fig. 1f,g**). **e)** In contrast to c) and d), calculated differential pulse responses (difference across contexts in projection onto the choice axis, as in Fig. **2d,e,f**) distinguish the different RNNs, e) Estimation of the differential pulse responses using kernel regression methods applicable to experimental data (see Methods), match the calculated differential pulse responses from **d). g)** The “slope index” (see Methods) quantifies the slope of the traces. Applied to the estimated differential pulse responses in **e)**, it has a monotonic relationship with d.i.m. percentage, and therefore can be used as a proxy measure for d.i.m. %. **h)** Differential pulse responses estimated from experimental data, for each of the FRO (examples arranged horizontally along the bottom) and LOG features (vertically on left), with the corresponding parallel indices plotted against each other. Arrows point to the parallel index value of each of the examples shown. Data from n=7 recorded rats. There is substantial heterogeneity in the differential pulse responses, suggesting heterogeneity in % d.i.m., both across animals, and within animals across the two features.

In contrast, the descriptions of the three components illustrated in Fig. 2d,e,f suggest that analyzing the system’s response to pulses of evidence would better distinguish different solutions-- an analysis that our pulse-based task is well suited to. A full characterization would require an estimate of each of the dynamics selection vectors **s**_**REL**_ and **s**_**IRR**_, which unfortunately are not directly observable. Nevertheless, the direction of the choice axis is straightforwardly estimated (Fig. 1), making the projection of the system’s state onto the choice axis a readily assayed measure. The bottom right panels of Fig. 2d,e,f, show that the difference across contexts of the time-evolution of this projection (the “differential pulse response”) can serve as an assay of the percentage of direct input modulation in the solution because it can distinguish solutions along the “Fast” vs “Slow” axis of Fig. 2g. This is illustrated in Fig. 3 using engineered RNNs, for which we can analytically compute their position on the barycentric coordinates (Fig. 3b), and can also directly measure the differential pulse response (Fig. 3e; see Methods). As a summary of the temporal shape of the differential pulse response, we use the slope of a straight line fit to it (“slope index”; see Methods); the smallest slope index corresponds to the top panel in Fig. 3f, and the largest slope index to the bottom panel. Fig. 3g confirms that in the RNNs, this slope index can be used as a measure of a network solution’s position on the “Fast” vs “Slow” axis. Based on previous approaches^28^, we developed kernel-based regression methods to measure the differential pulse response from neural activity recorded experimentally, and validated these methods in the RNNs (compare Fig. 3e and 3f). We then applied them to experimental data from each of our 7 rats and for each of the LOC and FRQ features (Fig. 3h). Strikingly, we did not find that a particular slope index consistently characterized solutions across animals. Instead, there was high variability across animals in this measure, and even across features within a single animal: no apparent correlation between the LOC and FRQ slope indices was visible (Fig. 3h main panel).

## A theoretical link between separate neural and behavioral measures is supported by the data

A widespread hypothesis in the field is that behavioral choices are driven by the system’s position on the choice axis^29–31^. If this is correct, then fast versus slow context-dependent effects on the choice axis, as produced by large versus small direct input modulation percentages (Fig. 3e,f), should have corresponding behavioral correlates. To assess the impact on behavioral choices of pulses at different times of a trial, we used logistic regression to compute behavioral kernels for LOC and FRQ evidence in each of the two LOC and FRQ contexts; each these kernels is a measure, from behavioral data, of the relative weight that evidence presented across different timepoints of a trial has on the subject’s choices (Methods). For a given feature, either LOC or FRQ, we refer to the difference across contexts as the “differential behavioral kernel” (panels along vertical and horizontal axes of Fig. 4; Extended Data Fig. 3). The shape of an individual’s differential behavioral kernel for one feature did not appear to predict the shape of the kernel for the other feature (Fig. 4 main panel), similar to our finding with the neural differential pulse responses (Fig. 3h). Nevertheless, the theory predicts that neural differential pulse responses and differential behavioral kernels should be tightly linked. Figs. 5a and 5b illustrate the concept. We use the simplifying assumption that the neural differential pulse response (Fig. 3) does not depend on time within a trial nor on previously presented evidence (See Extended Data Fig. 7g for data supporting this assumption). If *T* is the time at which position on the choice axis is read out to commit to a Right vs Left choice, then the context-dependent difference in the impact on behavioral choices of a pulse at time t will follow the neural differential pulse response at an interval *T*-*t* after the pulse. For direct input modulation, with a differential pulse response that is immediate and sustained (Fig. 3e,f top panels), the differential behavioral impact of a pulse should then be the same whether it is presented close to, or long before, the choice commitment time *T*, producing a flat differential behavioral kernel (i.e. slope index = 0 ; Fig. 5a). But for selection vector modulation or indirect input modulation, with differential pulse responses that grow only gradually from zero (Fig. 3e,f bottom panels), the differential behavioral impact of a pulse will be small if presented shortly before choice commitment, and larger if presented longer before. This should result in a converging differential behavioral kernel (i.e. slope index > 0; Fig. 5b). In other words, the shape of the differential behavioral kernel should be the reflection on the time axis of the differential pulse response. These two very different types of measures --behavioral versus neural-- are thus predicted to have the same slope index (but with opposite sign). We tested this prediction on RNNs engineered to solve the task using different amounts of direct input modulation. As predicted, the slope indices of the two different measures were tightly anti-correlated (Fig. 5c). We then tested whether a similar relationship existed for the rats’ behavioral and neural experimental data. To avoid any spurious correlations, we used different sets of pulses to assay each measure: we used pulses from the first half of the stimulus to measure the neural differential pulse response, and pulses from the second half of the stimulus to measure the differential behavioral kernel. We found robust support in the data for the theoretical prediction that the two measures should be correlated (Fig. 5d, r=-0.73, p<0.01), with the correlation also holding for LOC evidence alone (r=-0.71, p<0.1) or for FRQ evidence alone (r=-0.71, p<0.1). Thus, although there is no correlation within the neural measure (Fig. 3h) or within the behavioral measure (Fig. 4), and although the two measures were assayed on entirely different sets of pulses, the theoretical prediction that they should be strongly correlated was confirmed (Fig. 5d). These results support both the overall theoretical framework, which was built around the line attractor hypothesis for the choice axis from which behavior is read out, and the idea that variability in a solution’s position in the barycentric coordinates of Fig. 2g is the common source underlying, and explaining, the neural and behavioral variability in Fig. 3h and Fig. 4.

**Figure 4.**
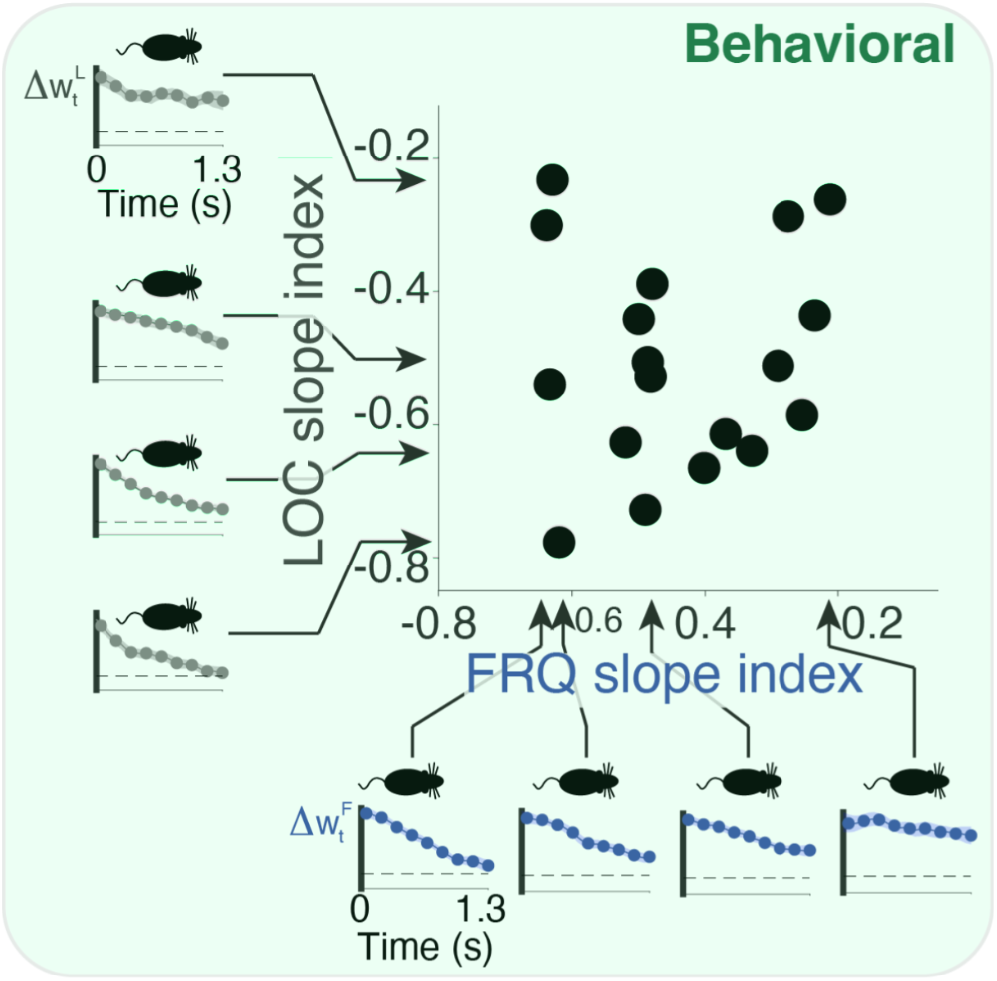
Differential behavioral kernels showed substantial heterogeneity across and within subjects, even while all these subjects performed the task well. Behavioral kernels are a behavior-based measure of how much weight the pulses from different timepoints within a trial have on a subject’s decision (Methods and Supplementary Figure 1). For a given feature, the differential behavioral kernel, shown here, is the difference in the behavioral kernel when that feature is in its relevant versus irrelevant context. Time axes run from start of the auditory pulse trains (t=O) to their end (t=1.3 sec). Figure conventions as in **Fig. 3h**, but data here is behavioral, not neural. n=20 rats.

**Figure 5.**
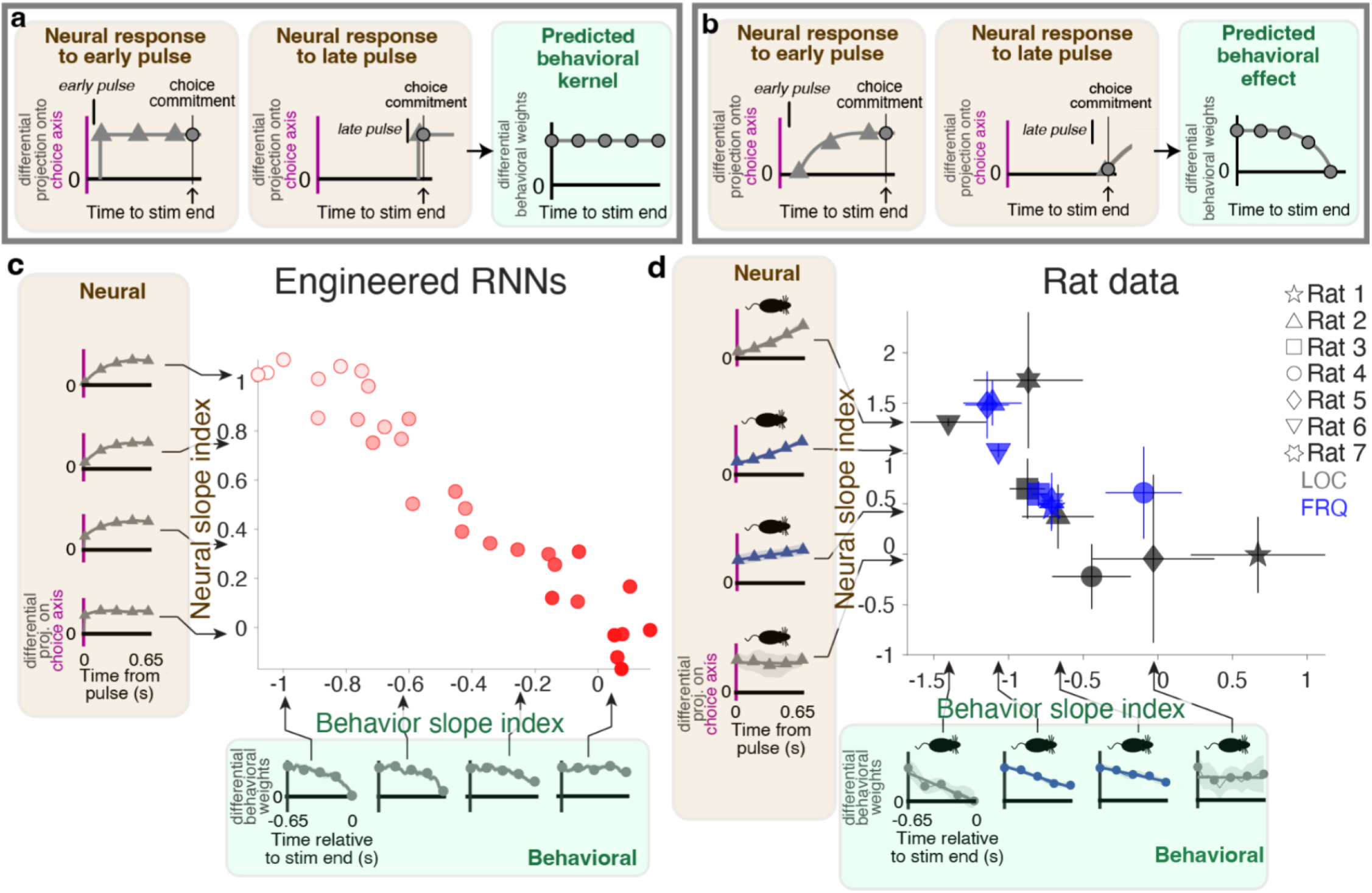
Theory predicts, and experimental data confirm, that variability in the neural slope index should explain variability in the behavioral slope index. **a,b)** Schematics of theoretical reasoning, **a)** In the case of a network using mostly direct input modulation, there will be immediate and sustained separation along the neural choice axis between relevant and irrelevant pulses (following **Fig. 3**). Thus the differential impact (across the two contexts) of a pulse on choice will not depend on whether the pulse is presented early (left panel) or late (middle panel) relative to choice commitment. The temporally flat differential pulse response of the neurons thus results in a temporally flat differential behavioral kernel (right panel), **b)** In contrast, if the network is using mostly selection vector modulation and/or indirect inputmodulation, pulses have a differential impact on choice only after the relaxation dynamics (**Fig. 3**). Pulses presented well before choice commitment will have a substantially different impact on choice in the relevant versus irrelevant contexts (left panel), while pulses presented immediately before choice commitment will have no time to give rise to a differential impact (middle panel). The gradually diverging differential pulse response of the neurons thus results in a gradually converging differential behavioral kernel (right panel), c) Data from n=30 engineered RNNs spanning the vertical axis of the barycentric coordinates (same colors as **Fig. 3**). Insets along the vertical axis are examples of neural differential pulse kernels (as in **Fig. 3**), each from a single RNN. Insets along horizontal axis are examples of differential behavioral kernels (as in **Fig. 4**). The RNN models follow the theoretical prediction, with highly correlated slope indices for the neural differential pulse kernels and differential behavioral kernels, **d)** Same conventions as in panel c, but now showing experimental data. To keep behavioral and neural estimates completely separate, only pulses from the first half (650ms) of the stimulus were used to compute neural differential pulse kernels, and only pulses from the last half (650ms) of the stimulus were used to compute differential behavioral kernels. The data follow the theoretical prediction, with highly correlated slope indices for the behavioral and neural measures. Individual data point shapes indicate each of LOC and FRQ features for each of the n=7 rats. Error bars indicate bootstrapped standard errors.

## Discussion

An influential conceptual approach known as “computation through dynamics”^7,19,32^ has posited that an understanding of neural activity from a mathematical dynamical systems perspective will allow elucidation of high-level phenomena such as cognition. Our work supports this view: starting from the longstanding hypothesis that decision evidence accumulation occurs along a line attractor (a concept drawn from dynamical systems; see Extended Discussion), with the system’s position on this line attractor driving choice behavior, and adding an algebraic rewriting of how the linearized dynamics around such an attractor would differ across two contexts, we developed a theory that describes and accounts for the variability in the properties of different solutions used by equally performant individuals. The theory predicted a tight link between otherwise disparate neural and behavioral measurements. This prediction was then found to be well supported in the data across multiple animals.

The approach led to multiple insights: theoretical insights, defining the space of possible solutions (Fig. 2g); biological insights, describing the behavioral, neural, and anatomical implications of the different solutions; conceptual insights, identifying the underlying source that links neural and behavioral variability (Fig. 5); and technical insights, allowing us to engineer recurrent neural networks that could not be constructed before, spanning the full space of solutions (Fig. 3a,b).

We describe our theoretical work as a “framework” because it does not specify particular network implementations. Instead, it defines axes onto which all possible dynamical solutions can be projected and described, with a solution’s position on this space quantifying how pulse-evoked dynamics change across contexts. The different components of the barycentric coordinates of Fig. 2g can also be viewed in terms of an associated latent circuit that clearly separates each component (Extended Data Fig. 9). Each point in the space constrains features of the circuits that map to it, but each point could nevertheless be implemented in multiple ways. Recent computational work has described several different implementations of context-dependent decision-making in recurrent neural networks (RNNs)^33–35^ (but see ref.^36^ regarding ref.^35^). Since the barycentric coordinates of Fig. 2g can be used to describe any network that solves the task with line attractors that are parallel across contexts (and see Extended Data Fig. 10 otherwise), all of the networks in refs ^33–35^, as well as in ref. ^4^, can be located on those coordinates. The rank-1 networks described in ref. ^33^ map onto points lying exclusively along the right edge of the triangle of barycentric coordinates in Fig 2g (i.e., the input modulation edge). This is because networks with a non-zero selection vector modulation component require rank 2 or higher (see Extended Discussion). The idealized latent network solution of ref. ^34^ (their Fig. 3b) maps onto the bottom right corner of Fig. 2g, 100% indirect input modulation. The recurrent network version of ref. ^35^ (their Fig. S5H), which modulates the linearized inputs and the recurrent dynamics equally, maps onto a point at the center of the left edge of the triangle. Finally, as described in Fig. 3, ref. ^4^ maps onto the bottom left corner, 100% selection vector modulation. All three of ref. ^4^, ref. ^34^, and ref. ^35^ each describe solutions that cover only a restricted region of the barycentric coordinates, and therefore do not address the variability we observed across individuals. (See Extended Discussion for more on the relationship between refs^33–35^ and our work.)

Our work also provides a cautionary note, highlighting the fact that trained RNNs, which are commonly used to model brain function^4,37–42^, need not comprise the full set of solutions consistent with the biological data. In our work we found that training led towards only one corner of the full space of solutions (Fig. 3a). It was a deeper understanding of the mathematics behind solutions (equations 1 and 2), not the use of trained networks, that allowed us to engineer data-compatible RNNs across the full space of solutions (Fig. 3b-f, Extended Data Fig. 6).

The interactions between afferent input signals and recurrent dynamics are a key part of understanding context-dependent computations. This view is closely related to the alignment of inputs and dynamics recently reported for sensory learning^43^. For example, large context-dependent changes in the sensory input (i.e., a large in equation 1) are not sufficient to conclude that those context-dependent changes in inputs drive context-dependent decision-making: only those input changes that are aligned to 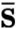, the average direction in neural space representing the recurrent dynamics, will produce a context-dependent effect on decisions (through 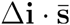). For the same reason, we note that although our data (Fig. 1) and that of ref. ^4^ are not compatible with “early gating” (i.e., blocking irrelevant evidence from reaching decision-making regions), the data are nevertheless compatible with input modulation (Fig. 3 and Extended Data Fig. 6). Several further studies have also provided evidence against early gating^5,6,44^, but there are nevertheless multiple studies providing evidence in favor of early gating^24,35,45,46^, making the issue a matter of ongoing debate. It has been argued that early gating is indicated by a representation of evidence in decision-making regions that is weaker in the irrelevant context (i.e., smaller magnitude |**i**| in our terminology)^45^, but example 3 in the Extended Discussion illustrates a counterexample in which the context with smaller |**i**| is actually the one where **i** has the greater impact on decisions, because it has the larger s·**i**; in other words, the interaction with recurrent dynamics needs to be taken into account before firm conclusions can be drawn. Similar to individual variability across the vertical axis of the solution space of Fig. 2g, which we believe is a result of all of the encompassed solutions being capable of solving the task, solutions either using or not using early gating are equally capable of solving the task (and both lie within the framework we have described; see examples 1 and 2 In Extended Discussion). It is thus possible that, regarding the use of early gating, there could be variability across tasks and individuals, perhaps even within them. Further work will be needed to resolve the relative prevalence, or absence, of early gating.

We have focused on the case where the choice axes of the two contexts are parallel to each other. A recent study^16^ reported that, in contrast to the findings of ref. ^4^ in monkey FEF and our findings in rat FOF, choice axes in monkey parietal cortex rotated across two task contexts. This motivated a broadening of our barycentric coordinates framework, and the Extended Discussion and Extended Data Fig. 10 describe how it can be extended to choice axes that rotate across contexts. In that more complex case, there are four components that add up to the net context-dependent effect, not three, and the barycentric coordinates therefore exist within a tetrahedron, not a triangle. But the core concepts of the framework remain the same. The same study^16^ further contrasted with the approximately linear choice axis that we (Fig. 1e, t=0 to t=1.3s) and others^4,15,29,47,48^ have found in that they reported a curved choice axis, due to a direction in neural space that encoded the magnitude of a trial’s difficulty, regardless of the sign of the subject’s upcoming choice. We speculate that differences across the studies could perhaps be explained by individual differences in the strength of difficulty encoding. In tasks or individuals where the difficulty encoding is stronger, the curvature would become a more significant feature.

Even though our experiments were carried out in rats, the similarity in the results of behavioral (Extended Data Fig. 1c,f) and neural (Fig. 1e,f) analyses that could be carried out in common with monkeys suggests that conclusions reached from rat data may generalize to other species as well. Using humans, a recent context-dependent decision-making study^49^ found that different stimulus features were processed independently. This finding is in line with our result that rat subjects can use separate mixtures of context-dependent components to select and accumulate each of the two features (Fig. 3h and Fig. 4).

Electrophysiological studies often center on findings that are similar across subjects, and it is common practice to report the result for an “average subject”. However, our results reveal a surprising degree of heterogeneity across, and even within, individual subjects, underscoring the importance of characterizing the computations used by each individual^50^. This issue may be of particular significance for cognitive computations, which are largely internal and therefore potentially subject to substantial covert variability across subjects. Here, studying how computations vary across subjects was made possible by two key methodologies. First, an efficient, automated procedure to train a sufficient number of rats to be able to observe and quantify cross-subject variability^10^. Second, characterization of each individual’s computations by leveraging the statistical power afforded by a randomly-timed, pulse-based stimulus^10^.

A limitation of our analyses of the experimental data is that we are currently unable to discriminate between mechanisms relying on context-dependent changes of recurrent dynamics (selection vector modulation) versus changes in the linearized sensory inputs (input vector modulation -- i.e. the oblique axis in Fig. 2g, bottom). A full characterization of the relevant neural dynamics will require estimation of the selection vector **s** for each context. Simultaneous recordings from large neural populations, combined with the application of recently-developed latent dynamics estimation methods, such as LFADS^51^ or FINDR^52^, may prove instrumental in future work in this direction. Another potential limitation stems from the possibility that recurrent dynamics might evolve more rapidly^53^ than the current time resolution in our measurements, leaving us unable to discriminate between contextual input modulation versus fast recurrent modulation. However, our results indicate that our analyses quantified the speed of evidence selection as smoothly varying across subjects (Fig. 3h; Fig. 4; Ext. Data Fig. 8), suggesting that in most subjects dynamics are slow enough to be captured with our method.

In sum, our work provides a new, general framework to describe and investigate artificial and biological networks for flexible decision-making, and opens the door to the cellular-resolution study of individual variability in neural computations underlying higher cognition.

## Supporting information

Extended discussion

## Data availability statement

All neural and behavioral data necessary to replicate the paper’s figures will be made available before the time of publication.

## Code availability statement

The code to train rats is available at https://github.com/Brody-Lab/flexible_decision_making_training. All the code for training, analysis, and engineering of RNNs will be made available before the time of publication at: https://github.com/Brody-Lab/flexible_decision_making_rnn. All the code for the analysis of neural data and behavior will also be made available before the time of publication.

## Acknowledgments

We thank S. Ostojic, K. Miller, S. Fusi and S. Druckmann for discussion and feedback on the manuscript. We thank J. Teran and C. Kopec for animal and laboratory support. This work was funded by the Howard Hughes Medical Institute and by NIH grant R21MH124383. M.P. was supported by a Simons Collaboration on the Global Brain Postdoctoral Fellowship, and by a Simons Foundation Autism Research Initiative Bridge to Independence Award.

## Author Contributions

M.P. and C.D.B. designed the experiment. M.P., V.M. and C.D.B. designed the automated training procedure.

M.P. and V.D.T. performed the experiments. M.C.A. and J.W.P. developed the mTDR analysis. M.P., M.C.A. and J.W.P. designed the pulse-based analysis of neural data. All authors contributed to the conceptual development of the theory. M.P. and C.D.B. developed the mathematical framework. M.P. and D.S. trained and analyzed artificial neural networks. M.P. and C.D.B. wrote the manuscript after discussions among all authors. C.D.B. supervised the project.

## Extended Data Figures

**Extended Data Figure 1.**
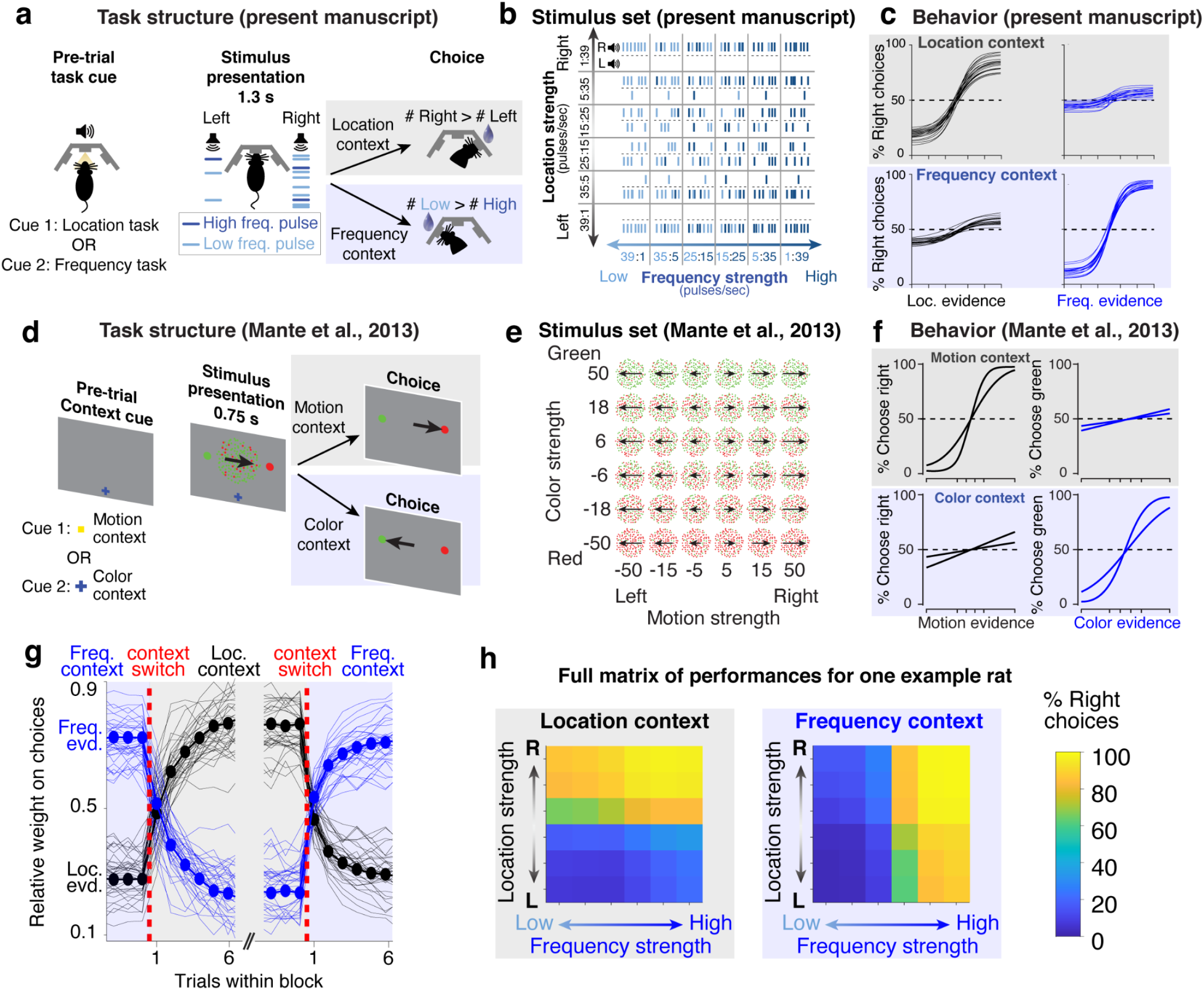
**a-d)** Comparison of rat task and monkey task^4^. **a)** In the rat task, the subject is cued using an audiovisual stimulus, and is presented with a train of randomly-timed auditory pulses varying in location and frequency. In different contexts, the subject determines the prevalent location or the prevalent frequency of the pulses. **b)** Stimulus set for the rat task: strength of location and prevalent frequency are varied independently on each trial. **c)** Psychometric curves for the rat task (n=20 rats). **d)** In the monkey task, the subject is cued using the shape and color of a fixation dot, and is presented with a field of randomly-moving red and green dots. In different contexts, the subject determines the prevalent color or the prevalent motion of the dots. **e)** Stimulus set for the monkey task: strength of motion and prevalent color are varied independently on each trial. **f)** Psychometric curves for the monkey task (n=2 macaque monkeys). **g)** Rats rapidly switch between contexts. Performances saturate within the first 4-5 trials in the block. The weight of location and frequency evidence is computed using a logistic regression (see methods). Thin lines indicate individual rats, thick lines indicate the average across rats. **h)** Full matrix of behavioral performances for one example rat across the two contexts.

**Extended Data Figure 2.**
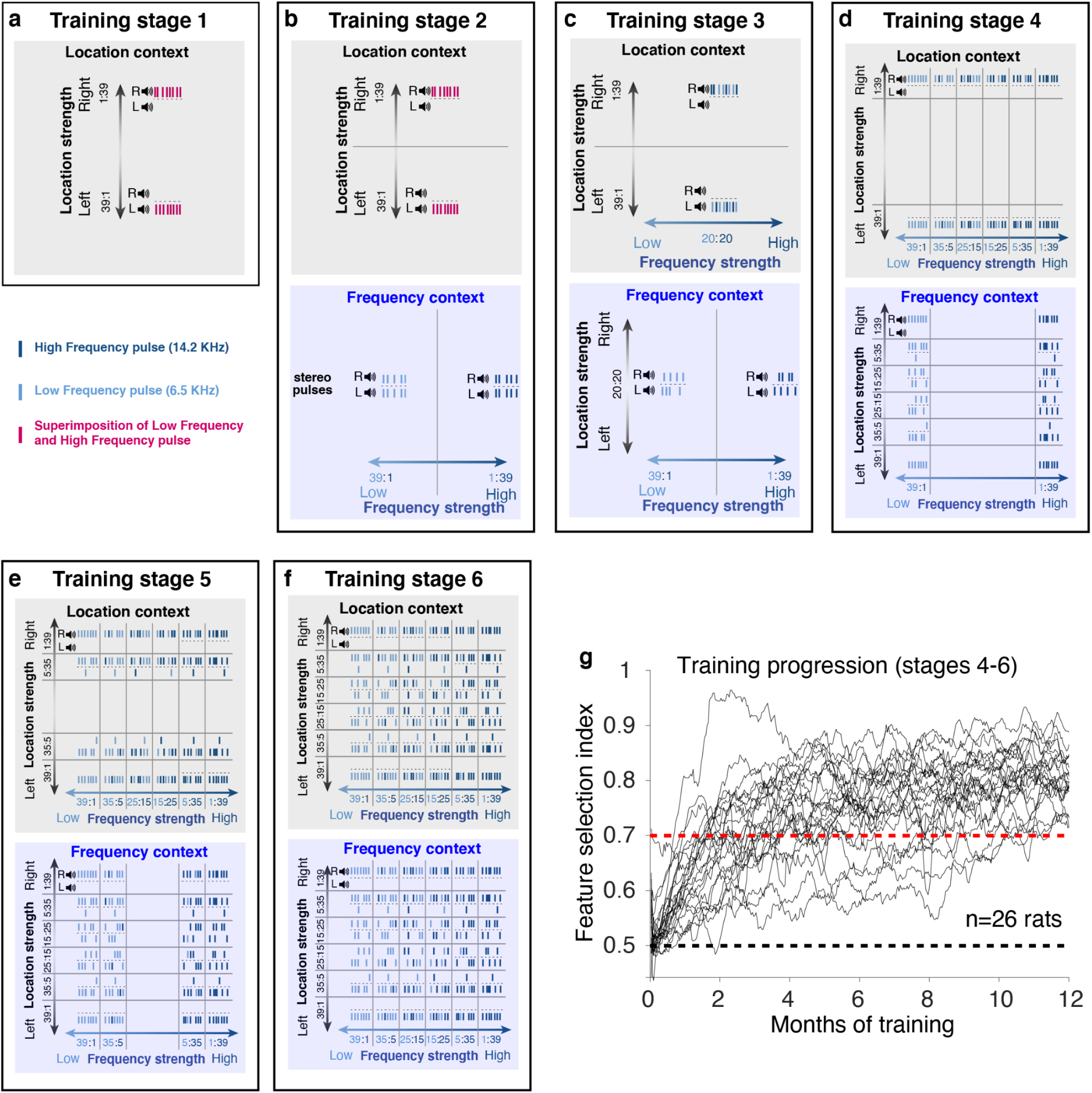
**a-f)** Training procedure. **a)** Stage 1: rats are trained only on the location task, with strong location evidence and no frequency evidence (pulses consist of superimposed low and high frequency). The context cue is played before each trial. **b)** Stage 2: rats learn to alternate between the location and frequency context. In the frequency context rats are presented with strong frequency evidence and no location evidence (stereo pulses). **c)** Stage 3: introduction of pulse modulation. In the frequency context, pulses are now presented on either side (but with no prevalent side). In the location context, pulses are either high-frequency or low-frequency (but with no prevalent frequency). **d)** Stage 4: irrelevant information is introduced, but the relevant information is always at maximum strength. **e)** Stage 5: relevant information can have intermediate strength. **f)** Stage 6: relevant information can have low strength. **g)** Training progression. Most rats learn stages 1-3 in approximately 2 weeks, but it takes a much longer time to learn stages 4-6 because of the introduction of irrelevant evidence. The feature selection index quantifies whether rats attend to the correct feature and ignore the irrelevant feature (see methods). The black dashed line indicates chance, the red dashed line indicates the threshold performance to consider a rat trained. Most rats learn the task within 2-5 months.

**Extended Data Figure 3.**
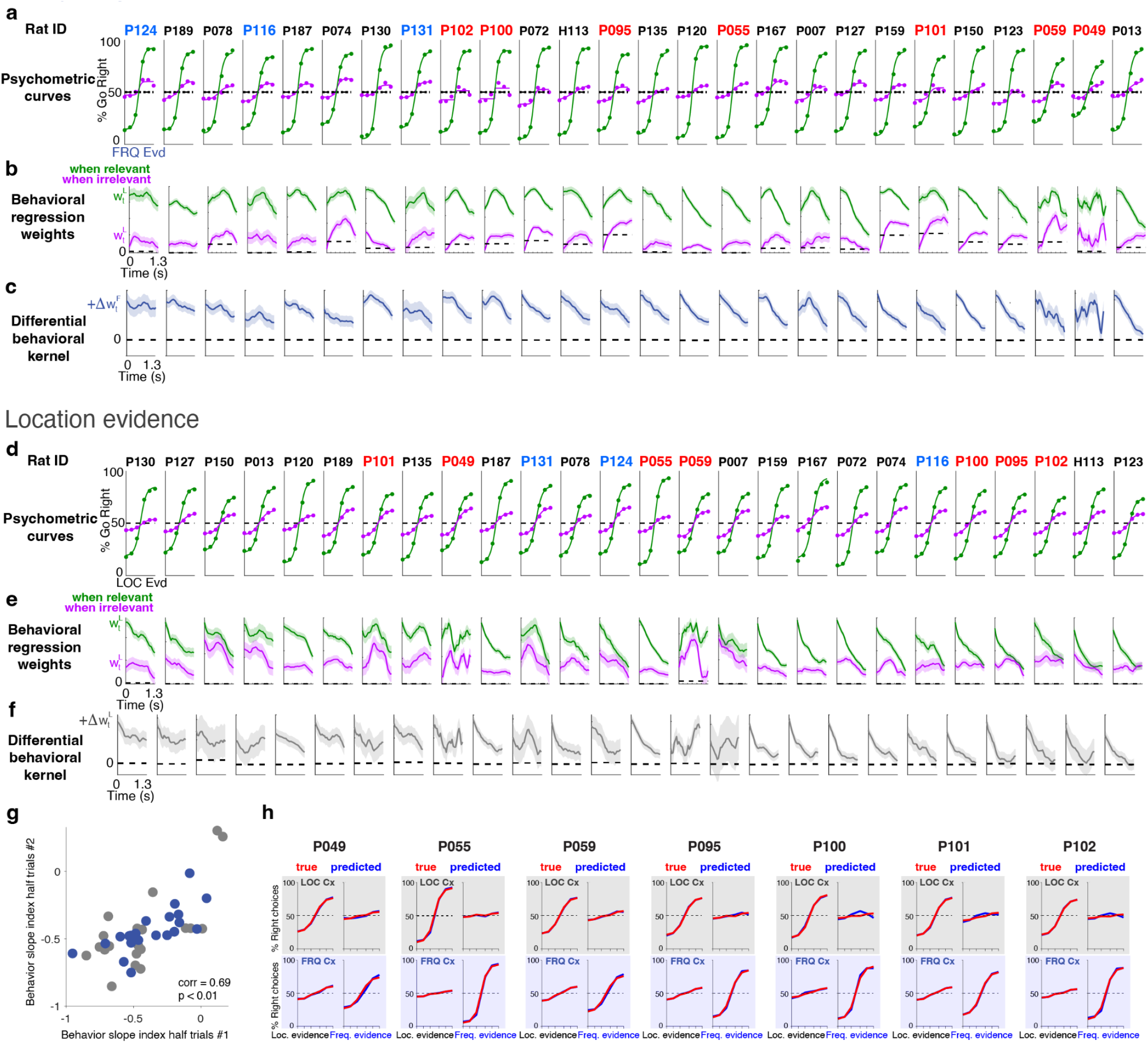
Behavioral data for all rats. Rat ID color indicates whether rat was used for electrophysiology (red), optogenetics (cyan) or only for behavior (black). **a)** Psychometric curves for frequency evidence, measuring the fraction of right choices as a function of strength of frequency evidence (6 levels of strength, see Fig. 1b). Green indicates frequency context (relevant), purple indicates location context (irrelevant). **b)** Weights for frequency evidence computed using the behavioral logistic regression for each rat (see Fig. 1d); colors as in panel a. **c)** Differential behavioral kernel for frequency evidence across all rats. **d)** Psychometric curves for location evidence, measuring the fraction of right choices as a function of strength of location evidence (6 levels of strength, see Fig. 1b). Green indicates location context (relevant), purple indicates frequency context (irrelevant). **e)** Weights for location evidence computed using the behavioral logistic regression for each rat (see Fig. 1d); colors as in panel d. **f)** Differential behavioral kernel for location evidence across all rats. Shaded areas indicate bootstrapped standard errors. **g)** The slope index computed from behavioral trials in the first half split is highly correlated with the slope index computed using the second half split. **h)** Psychometric curves can be predicted with high precision from the weights of the logistic regression. Data are shown from the seven rats used for electrophysiology recordings.

**Extended Data Figure 4.**
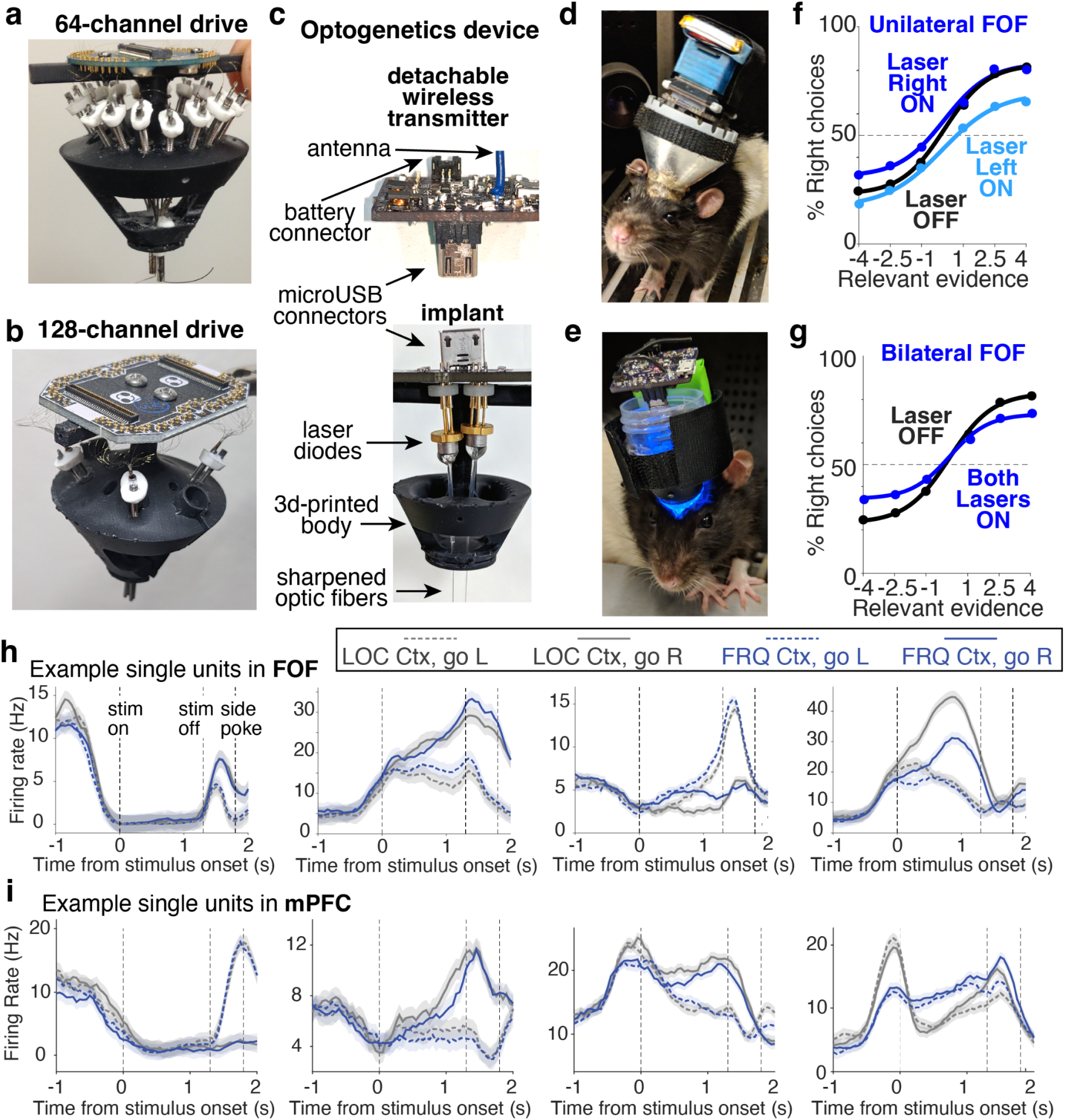
**(a)** 64-channel custom-made multi-tetrode drive, allowing independent movement of 16 tetrodes. This drive was used in one rat for wired recordings. **(b)** 128-channel custom-made multi-tetrode drive, allowing independent movement of 4 bundles with 8 tetrodes each. This drive was used in six rats for wireless recordings. **(c)** Device for wireless optogenetic perturbation. In the implant, two chemically sharpened optic fibers targeting both hemispheres are attached using optical glue to two laser diodes. The laser diodes are controlled independently by a control board, which communicates wirelessly with the computer controlling the behavior. The control board can be attached/detached using a microUSB connector. **(d)** Example rat with wireless electrophysiology implant and headstage. **(e)** Example rat with wireless optogenetic implant and control board. (**f-g)** Result of inactivation of FOF. 3 rats expressed AAV2/5-mDlx-ChR2-mCherry and were stimulated with blue light (450 nm, 25mW) for the full duration of the stimulus. **(f)** Result of unilateral inactivation on rats’ choices as a function of strength of relevant evidence (averaged across the two contexts). Activation of each laser was randomized across trials. **(g)** Result of bilateral FOF inactivation on rats’ choices as a function of strength of relevant evidence (averaged across the two contexts). **(h,i)** Example responses of single units recorded in FOF **(h)** and in mPFC **(i)**. Shown are the peri-stimulus time histograms of responses for correct trials, averaged according to context and choice. Units in both areas exhibit significant heterogeneity and large modulation according to combinations of the rat’s upcoming choice and the current context. The dashed vertical lines indicate the beginning of the pulse-train stimulus presentation, the end of the pulse-train stimulus presentation, and the average time when the rat performed a poke in one of the two side ports to indicate his choice. Shaded areas indicate standard errors.

**Extended Data Figure 5.**
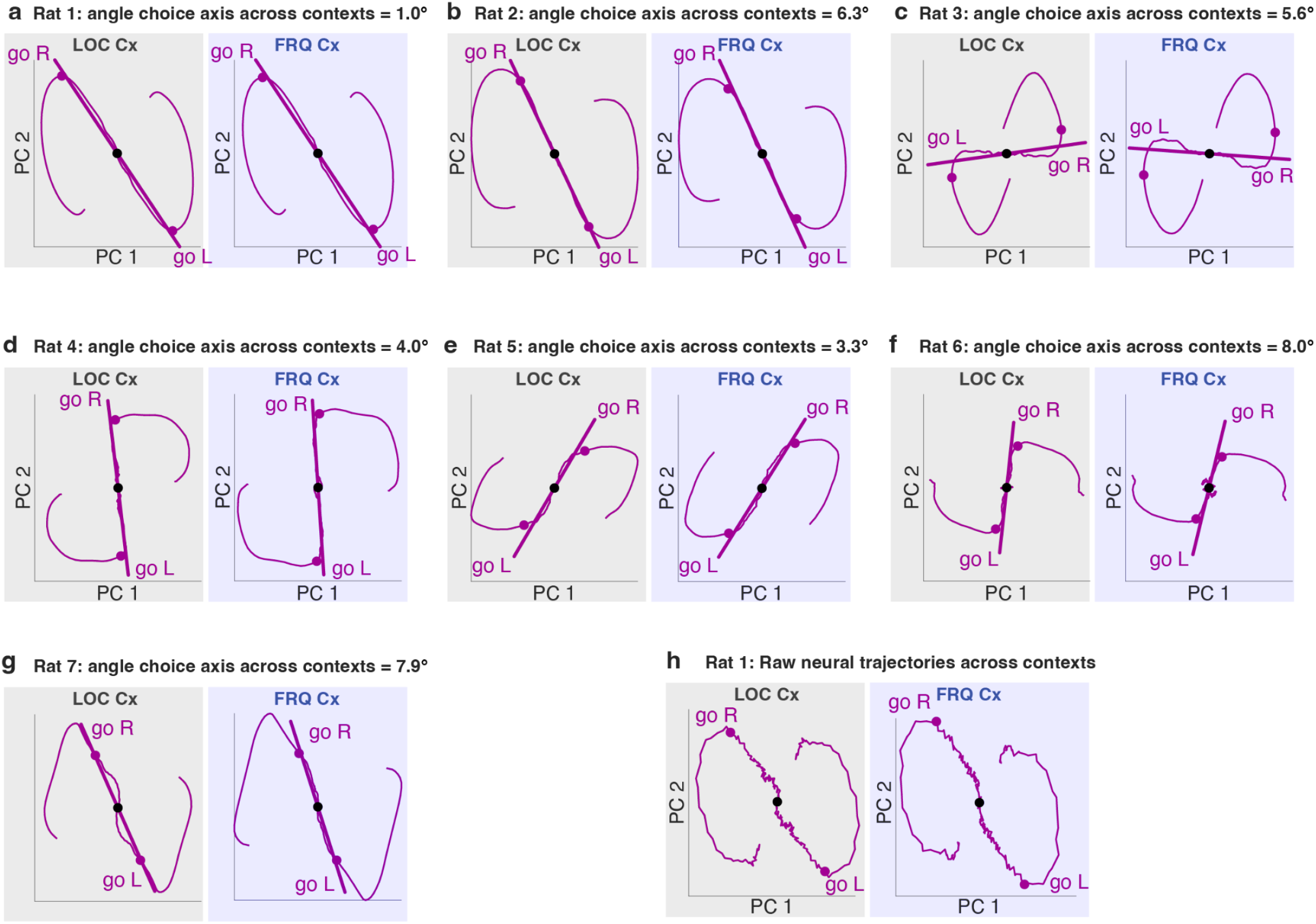
Choice-related dynamics, computed independently for each rat, and across the two contexts. For each rat, the horizontal and vertical axes in the two subpanels are the same across the two panels, and are computed using data from both contexts. In panels a-g, the dynamics in each context are computed using the choice kernels of the pulse-based regression (see Fig. 1.1 in Extended Discussion). The kernels provide a regularized, noise-reduced version of the raw trajectories (which are shown for Rat 1 in panel h). The black dot indicates the time of the start of stimulus presentation (t=0), the purple dots indicate the end of stimulus presentation (t=1.3s). The line indicates the choice axis computed in the given context, and above the panels is indicated the angle between the choice axes computed across the two contexts.

**Extended Data Figure 6.**
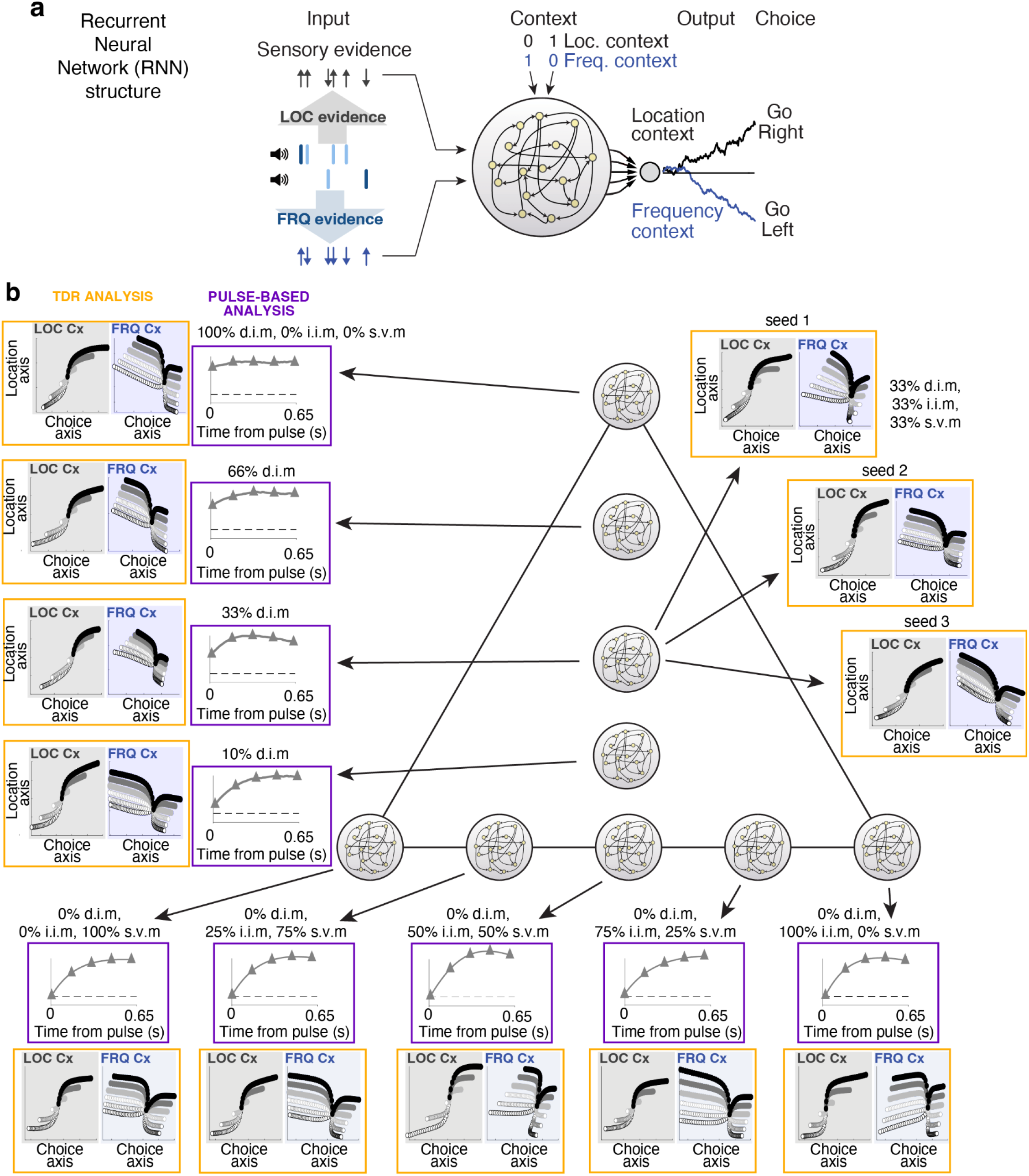
Engineered recurrent neural networks (RNNs) across the entire solution space (Fig. 2g) all qualitatively reproduce rat TDR trial-based dynamics, but are distinguished by pulse-based analysis. **a)** Architecture of the RNNs. **b)** TDR analysis (orange frame) and pulse-based analysis (purple frame) applied to RNNs generated to span different points within the solution space, as indicated by the RNN symbol on the barycentric coordinates. The TDR analysis and the pulse-based analysis of one RNN at each position are shown, connected to their RNN position by the arrow. For the position at the very center of the triangle, three different RNNs at that position, trained by starting from different random initial weights, are shown. All RNNs qualitatively reproduce rat TDR trial-based dynamics. The variability of trial-based TDR seen across RNNs is not predictive of the position within the solution space, and even RNNs generated from the same point can produce variable TDR trajectories. In contrast, the estimated pulse-triggered response reliably indicates the position of RNNs along the vertical axis of the solution space.

**Extended Data Figure 7.**
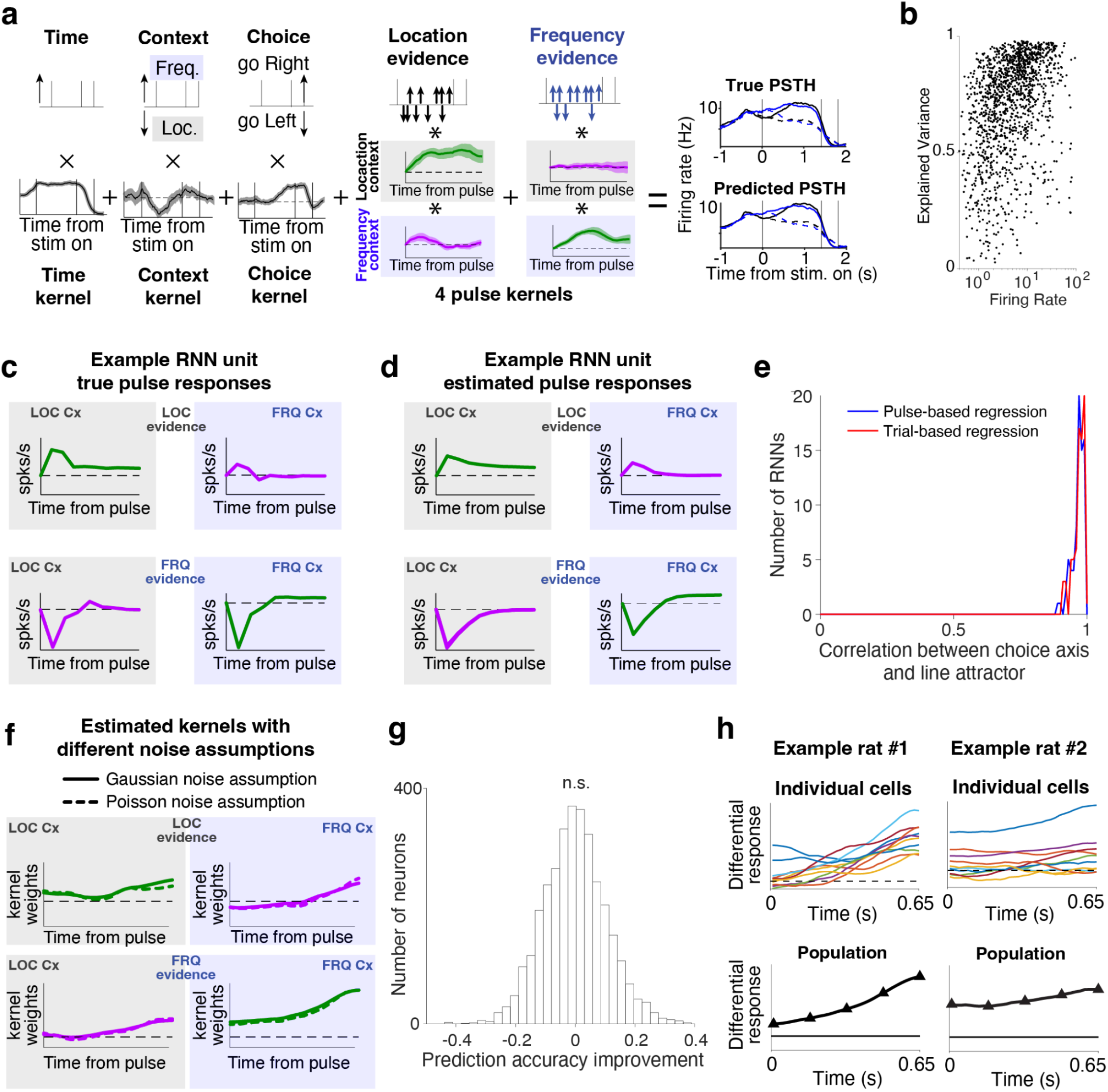
Validation of pulse regression method. **a)** Example application of the pulse regression to one example recorded unit. **b)** Fraction of explained variance as a function of firing rate across all recorded units. **c-d)** The pulse-regression kernels provide an accurate estimate of the response to a single isolated pulse. In **c)** are shown the responses to a single isolated pulse of either location or frequency evidence in both contexts for an example RNN unit. In **d)** are shown the estimates of these pulses from the dynamics of the RNN solving the task with regular trials featuring many consecutive pulses presented at 40Hz. **e)** Comparison of the direction of the true line attractor (computed by finding the RNN’s fixed points, see methods) with the choice axis estimated by the trial-based regression (Fig. 1f,g) and the pulse-based regression (Fig. 3). The choice axis closely approximates the direction of the true line attractor. **f)** Kernels estimated using the assumption of gaussian noise closely approximate those estimated using the assumption of Poisson noise. Kernels are shown here for one example neuron. **g)** Prediction accuracy does not improve when two separate kernels are computed for the early portion of the stimulus and the late portion of the stimulus. Here is shown the improvement in cross-validated prediction accuracy across all recorded neurons when using two separate kernels as compared to using a single kernel throughout the stimulus.

**Extended Data Figure 8.**
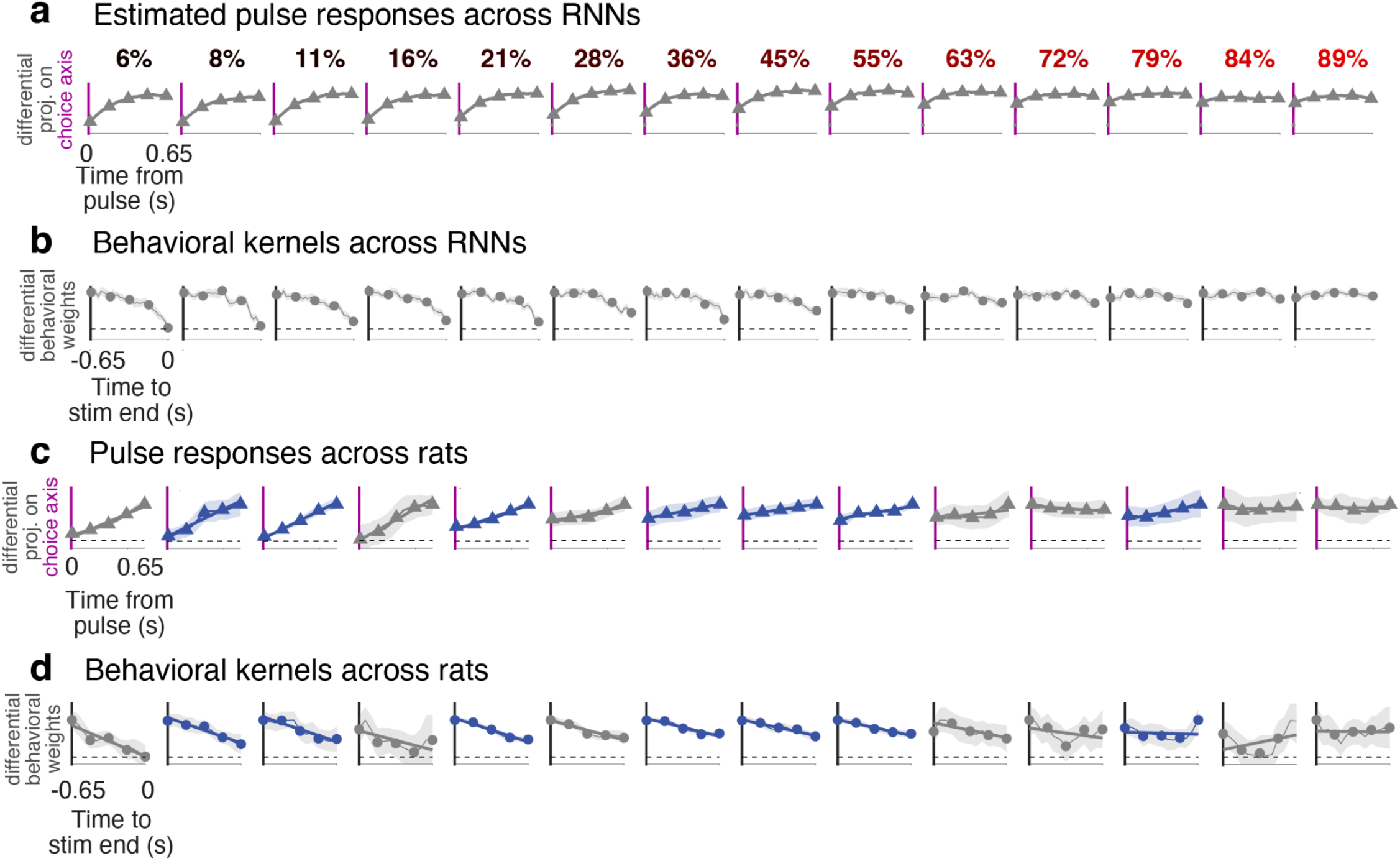
**(a)** Differential pulse responses across the RNNs shown in Fig. 5c. The number above each behavioral kernel indicates the fraction of direct input modulation for the associated RNN (same notation as in Extended Data Figure 6). **(b)** Corresponding behavioral kernel for each RNN. **(c)** Differential pulse responses across all rats shown in Fig. 5d (n=7 rats, two features per rat). Gray indicates location feature, blue indicates frequency feature. **(d)** Corresponding behavioral kernels for each rat and feature. Shaded areas indicate bootstrapped standard errors.

**Extended Data Figure 9.**
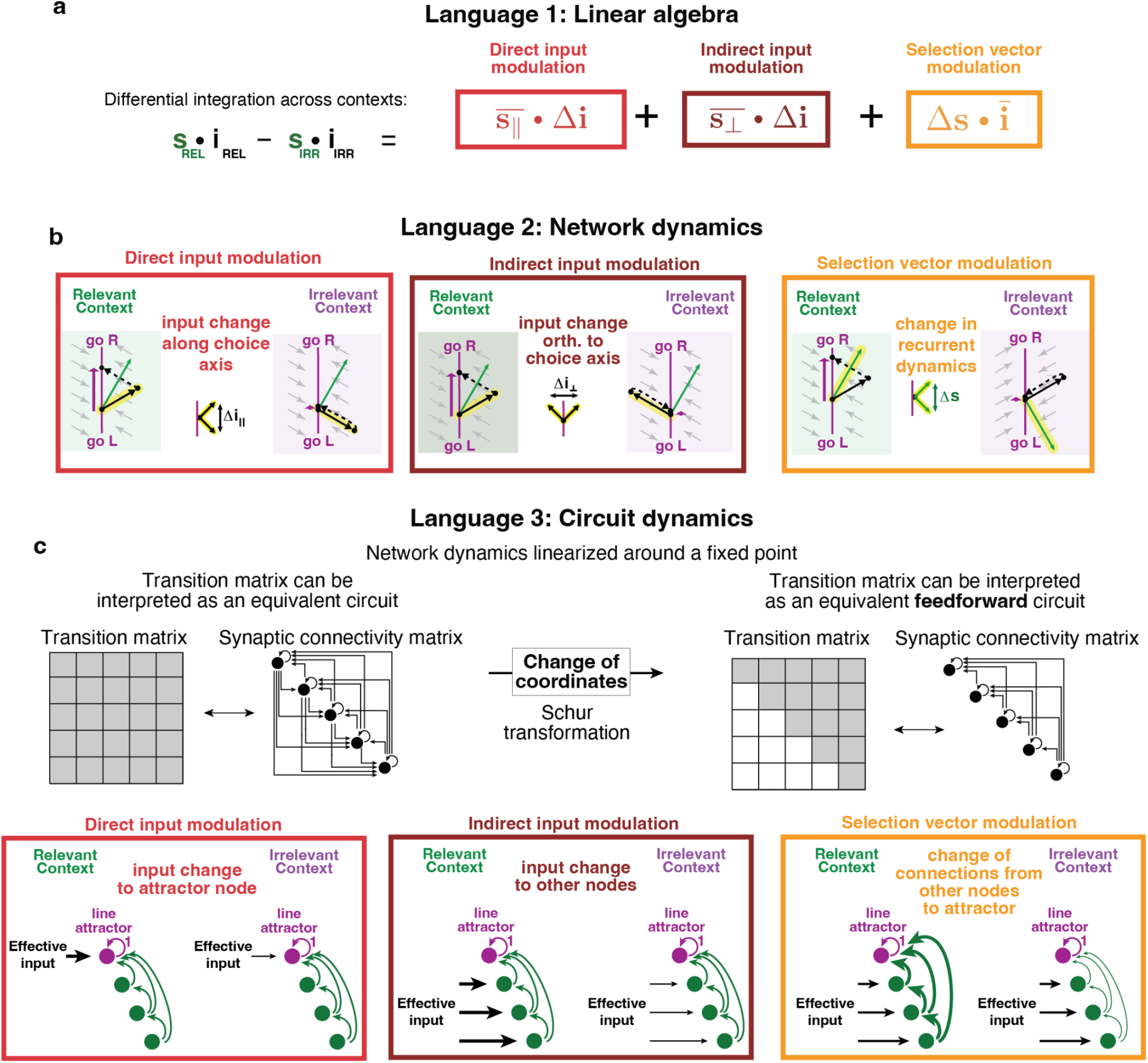
Three distinct “languages” can capture the three fundamental solutions to the task. **(a)** “Linear algebra language”. As derived in the main text in Equations 1 and 2, the overall differential integration can be expressed as a sum of three terms. **(b)** “Network dynamics language”. The three solutions are associated with distinct pulse-evoked dynamics within the space spanned by the line attractor and the selection vector. **(c)** ”Circuit dynamics language”. The three solutions are associated with three different latent circuit structures. To show this, we first note that our derivation of task solutions stems from focusing on linearized dynamics around fixed points of a line attractor (Fig 2c). These linearized dynamics can be interpreted as an equivalent linear circuit whose synaptic connectivity matrix is defined by the state transition matrix (i.e. matrix M in Equation (1)). This circuit can be further simplified into a feedforward circuit using the Schur transformation (Goldman, 2009), which operates a change of coordinates to transform the state transition into an upper triangular form. In the resulting circuit, the first node represents the accumulator (i.e. the line attractor), and it receives feed-forward inputs from the other nodes of the circuit. Our three solutions can be interpreted as three different ways to modulate the connectivity of this circuit across the two contexts. In the case of “direct input modulation”, it is the input to the accumulator node that varies across contexts. In the case of “indirect input modulation”, it is the input to the other nodes that changes across contexts, and this differential input eventually reaches the accumulator through the feed-forward connections. Finally, in the case of “selection vector modulation”, the input to all nodes stays the same across contexts, but the feed-forward connections between the other nodes and the accumulator node change across contexts.

**Extended Data Figure 10.**
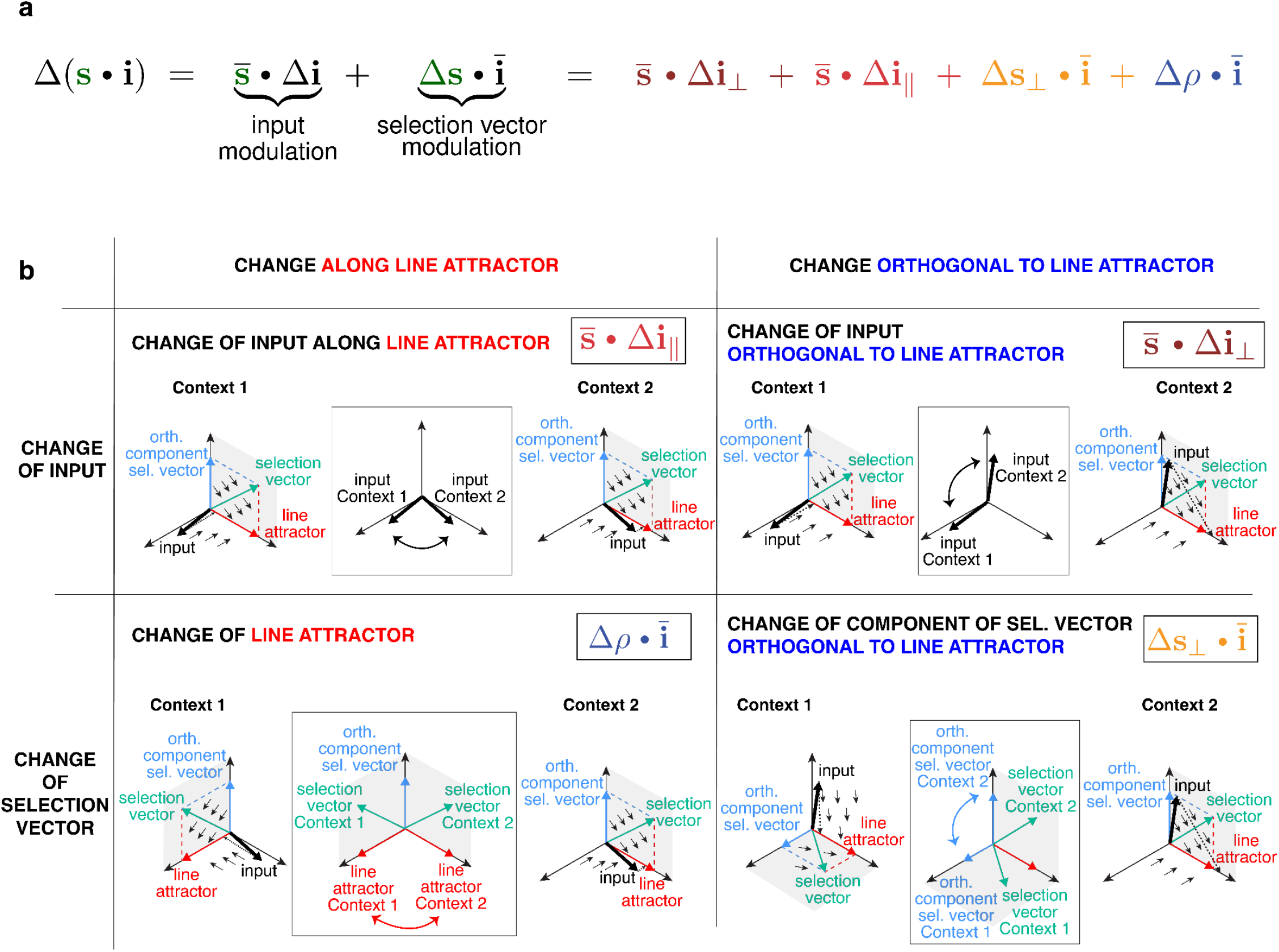
Extension of the theory to the general case with context-dependent line attractors. **a)** Rewriting of the equation describing the differential integration of a pulse across contexts (Equation 1) after the assumption that the line attractor is parallel across the two contexts is dropped. In this equation, the first three terms correspond to the same terms as in Equation 2, with addition of a fourth term, which captures changes in the direction of the line attractor along the average input direction. **b)** Graphical intuition of the four solutions in the general case where the line attractor is not parallel across the two contexts. Top left: changes of the input along the direction of the line attractor (“direct input modulation”). Top right: changes of the input along a direction orthogonal to the line attractor (“indirect input modulation). Bottom left: changes of the direction of the line attractor across the two contexts. Bottom right: changes of the component of the selection vector orthogonal to the line attractor across contexts (“selection vector modulation”).

## Methods

### 1 Subjects

All animal use procedures were approved by the Princeton University Institutional Animal Care and Use Committee (IACUC) and were carried out in accordance with NIH standards. All subjects were adult male Long-Evans rats that were kept on a reversed light-dark cycle. All training and testing procedures were performed during the dark cycle. Rats were placed on a restricted water schedule to motivate them to work for a water reward. A total of 26 rats were used for the experiments presented in this study. Of these, 7 rats were used for electrophysiology recordings, and 3 rats were implanted with optical fibers for optogenetic inactivation.

### 2 Behavior

All rats included in this study were trained to perform a task requiring context-dependent selection and accumulation of sensory evidence (Figure 1a). The task was performed in a behavioral box consisting of three straight walls and one curved wall with three “nose ports”. Each nose port was equipped with an LED to deliver visual stimuli, and with an infrared beam to detect the rat’s nose entrance. In addition, above the two side ports were speakers to deliver sound stimuli, and water cannulas to deliver a water reward. At the beginning of each trial, rats were presented with an audiovisual cue indicating the context of the current trial, either “location context” or “frequency context”. The context cues consisted of 1s-long, clearly distinguishable FM modulated sounds, and in addition the “location context” was signaled by turning on the LEDs of all three ports, while in the “frequency context” only the center LED was turned on. After the end of the context cue, the rats were required to place their nose into the center port. While maintaining fixation in the center port, rats were presented with a 1.3s-long train of randomly-timed auditory pulses. Each pulse was played either from the speaker to the animal’s left or from the speaker to their right, and each pulse a 5 ms pure tone with either low-frequency (6.5 KHz) or high frequency (14 KHz). The pulse trains were generated by Poisson processes with different underlying rates. The strength of the location evidence was manipulated by varying the relative rate of right vs left pulses, while the strength of the frequency evidence was manipulated by varying the relative rate of high vs low pulses (Fig. 1b). The overall pulse rate was kept constant at 40 Hz. In the ”location context”, rats were rewarded if they turned, at the end of the stimulus, towards the side that had played the greater total number of pulses, ignoring the frequency of the pulses. In blocks of ”frequency” trials, rats were rewarded for orienting left if the total number of low frequency pulses was higher than the total number of high frequency pulses, and orienting right otherwise, ignoring the location of the pulses. The context was kept constant in blocks of trials, and block switches occurred after a minimum of 30 trials per block, and when a local estimate of performance reached a threshold of 80% correct. Behavioral sessions lasted 2-4 hours, and rats performed on average 542 trials per session. On average, rats switched across 14.6 context blocks per session.

### 3 Electrophysiology

Tetrodes were constructed using nickel/chrome alloy wire, 12.7 *µ*m (Sandvik Kanthal), and were gold-plated to 200 kΩ at 1 kHz. Tetrodes were mounted onto custom-made drives (Ext. Data Fig. 4a,b)[2], and the microdrives were implanted using previously described surgical stereotaxic implantation techniques[6]. Five rats were implanted with bilateral electrodes targeting FOF, centered at +2 anteroposterior (AP), ±1.3 mediolateral (ML) from bregma, while two rats were implanted with bilateral electrodes targeting the prelimbic (PL) area of mPFC, with coordinates +3.2 anteroposterior (AP), ±0.75 mediolateral (ML) from bregma. In one rat with an implant in FOF, 16 tetrodes were connected to a 64-channel electronic interface board (EIB), and recordings were performed using a wired setup (Open-Ephys). In the other six rats, 32 tetrodes per rat were connected to a 128-channel EIB and recordings were performed using wireless headstages (Spikegadgets; Ext. Data Fig. 4d).

### 4 Optogenetics

Preparation of chemically-sharpened optical fibers (0.37 NA, 400 *µ*m core; Newport) and basic virus injection techniques were the same as previously described[6]. At the targeted coordinates (FOF, +2 AP mm, ±1.3 ML mm from bregma), injections of 9.2nl of adeno-associated virus (AAV) (AAV2/5-mDlx-ChR2-mCherry, three rats) were made every 100 *µ*m in depth for 1.5mm. Four additional injection tracts were completed at coordinates 500 *µ*m anterior, 500 *µ*m posterior, 500 *µ*m medial and 500 *µ*m lateral from the central tract. In total, 1.5*µ*l of virus was injected over approximately 30min. Chemically sharpened fibers were lowered down the central injection track. Virus expression was allowed to develop for 8 weeks before optogenetic stimulation began. Optogenetic stimulation was delivered at 25mW using a customized wireless system derived from the “Cerebro” system (https://karpova-lab.github.io/cerebro ; Ext. Data Fig. 4c,e)[10, 3].

### 5 Analysis of behavior

Data was extracted from all behavioral sessions in which rats’ fraction of correct responses was equal or above 70%, feature selection index (see below) was equal or above 0.7, and in which rats performed at least 100 trials. Analysis of behavior was performed for all rats with electrophysiology or optogenetics implants, as well as for all other rats that performed at least 120,000 valid trials, i.e. where the rat maintained fixation for the full duration of the pulse train before making a decision. Psychometric curves (Fig. 1c; Extended Data Figure 3) were used to display the fraction of rightward choices as a function of the difference between the total number of right pulses and left pulses (location evidence strength), and as a function of the difference between the total number of high pulses and low pulses (frequency evidence strength). These curves were fit to a 4-parameter logistic function[4]:

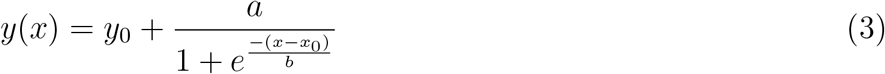

To quantify whether a rat selected the contextually relevant evidence to form his decisions on a given session, we computed a “feature selection index”. For this purpose, we performed a logistic regression for each of the two contexts, where the rat’s choices were fit as a function of the strength of location and frequency evidence. For each context, we considered all valid trials, and we compiled the rat’s choices, as well as the strength of location and frequency evidence. The vector of choices was parameterized as a binary vector (Right = 1; Left = 0), the strength of location evidence was computed as the difference between the rate of right and the rate of left pulses, while the strength of frequency evidence was computed as the difference between the rate of high-frequency and the rate of low-frequency pulses. In the location context, we fit the probability of choosing right on trial k using the logistic regression:

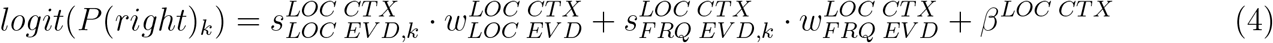

where 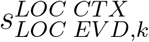 indicates the strength of location evidence on trial k, 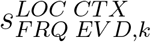 indicates the strength of frequency evidence on trial k, 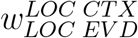 is the weight of location evidence on the rat’s choices, 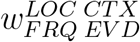 is the weight of frequency evidence on the rat’s choices, and 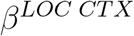 is a bias term. The relative weight of location evidence in the location context was computed as:

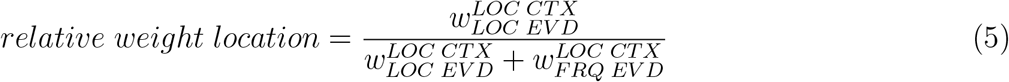

Similarly, in the frequency context we fit the rat’s choices as:

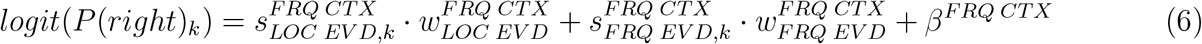

where 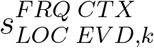 indicates the strength of location evidence on trial k, 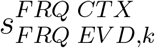 indicates the strength of frequency evidence on trial k, 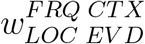 is the weight of location evidence on the rat’s choices, 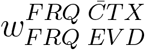 is the weight of frequency evidence on the rat’s choices, and 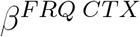 is a bias term. The relative weight of location evidence in the frequency context was computed as:

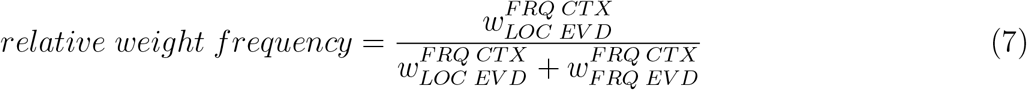

Finally, the feature selection index was then computed as the average between the relative weight of location in the location context (Eq. 5), and the relative weight of frequency in the frequency context (Eq. 7):

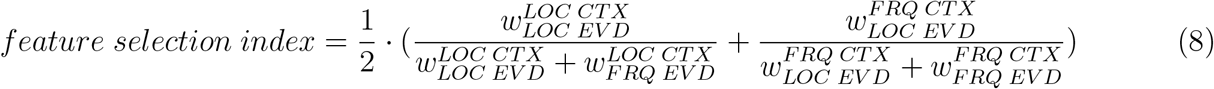

The feature selection index was used to precisely quantify the rats’ learning during training, as this metric allows to compare data across stages with different evidence strength (Extended Data Fig. 2g). In addition, the relative weight of location and frequency were computed for each rat as a function of the position of a trial within the block (e.g. immediately after a block switch, one trial after a block switch etc.), providing a measure of the rats’ ability to rapidly switch attended feature upon context switching (Extended Data Fig. 1g).

#### 5.1 Behavioral logistic regression

To quantify the dynamics of evidence accumulation, behavioral data was analyzed using another logistic regression. Importantly, in Eq. 5 and 7 we quantified the rat’s weighting of evidence using a single number, because we considered the generative rates, i.e. the expected strength of location and frequency evidence on a given trial. Now, we seek instead to quantify how these weights vary throughout stimulus presentation, by taking advantage of the knowledge of the exact pulse timing. For each rat, data across all sessions was compiled into a single vector of choices (Right vs Left), and two matrices detailing the pulse information presented on every trial. More specifically, the choice vector was parameterized as a binary vector (Right = 1; Left = 0), with dimensionality N, where N is the total number of valid trials. Pulse information was split into location evidence and frequency evidence, and was binned into 26 bins with 50 ms width. For a given bin, the amount of location evidence was computed as the natural logarithm of the ratio between the number of right and the number of left pulses, and was compiled in a location pulse matrix *X*^*L*^ with dimensionality N x 26. Similarly, frequency evidence was computed as the logarithm of the ratio between high-frequency and low-frequency pulses, and was compiled into a frequency pulse matrix *X*^*F*^ with dimensionality N x 26. We chose to use the logarithm of the ratio instead of the difference because it provided a better fit to cross-validated data. To quantify the impact on choices of evidence presented at different time points we fit a logistic regression, where the probability of choosing right at trial k was given by:

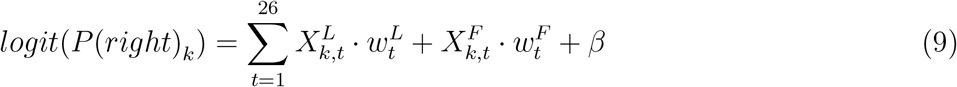

where 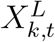 indicates the location evidence at time t on trial k, 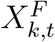 indicates the frequency evidence at time t on trial k, 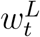 indicates the location weight at time t, 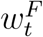 indicates the frequency weight at time t, and *β* indicates the bias to one particular side. Weights were fit using ridge regression, and the ridge regularizer was chosen to optimally predict cross-validated choices. The regression was applied separately for trials in the location context, and trials in the frequency context, resulting in four sets of weights computed for each rat (Figure 1.2 in Extended Discussion). To study how evidence was differentially integrated across the two contexts, we then computed a differential behavioral kernel. The location differential kernel was equal to the difference between the location weights computed in the location context, and the location weights computed in the frequency context. Similarly, the frequency differential kernel was equal to the difference between the frequency weights computed across the two contexts.

To quantify the shape of the differential behavioral kernels, we computed a behavioral ”slope index”. To obtain this, we computed the straight line that provided the least-square fit of the difference between the weights across the two contexts. The slope index was defined as the slope of this fitting line.

As a result, a slope index = 0 indicates that the fitting line is perfectly horizontal (i.e. the difference between the two sets of weights is constant at all time points), while a slope index *<* 0 indicates a decreasing difference between the weights across contexts, and a slope index *>* 0 indicates a rising difference. Empirically we found that differential behavioral kernels predominantly displayed convergence towards the end of the pulse stimulus presentation (Fig. 4, Extended Data Fig. 8).

### 6 Analysis of neural data

Spike sorting was performed using MountainSort[5], followed by manual curation of the results. 3285 putative single units were recorded from 5 rats in FOF (number of units in each rat: 2047, 832, 258, 94, 54), while 210 units were recorded from 2 rats in mPFC (number of units in each rat: 112, 98). To measure the responses of individual neurons, peri-stimulus time histograms (PSTH) were computed by binning spikes in 20 ms intervals, and averaging responses for trials according to choice and context. Responses of single neurons in both areas were highly heterogeneous and multiplexed multiple types of information (Extended Data Fig. 4), and no systematic difference was found in the encoding of task variables across the two regions (see e.g. Extended Data Fig. 5), so all studies of neural activity were carried out at the level of neural populations, and pooling data from FOF and from mPFC.

#### 6.1 Trial-based targeted dimensionality reduction (TDR) analysis of neural population dynamics

To study trial-averaged population dynamics, we applied model-based targeted dimensionality reduction (mTDR)[1], a dimensionality-reduction method which seeks to identify the dimensions of population activity that carry information about different task variables. This method was applied to our rat dataset, and to reanalyze a dataset collected while macaque monkeys performed a similar visual task [7] (Extended Data Fig. 1). In brief, the goal of mTDR is to identify the parameters of a model where the activity of each neuron is described as a linear combination of different task variables (choice, time, context, stimulus strength). For each of these task variables, the model retrieves a time-varying weight vector *w*_*i*_(*t*) (with number of elements, indexed by *i*, equal to the number of recorded neurons) specifying the linear relationship between the value of that variable and the activity of each neuron at each timepoint (each variable *v* contributes an additive component *v · w*_*i*_(*t*) to the firing rate of neuron i), and the collection of these weight vectors across all neurons are constrained to form a low-rank matrix. Singular Value Decomposition of this low-rank weight matrix is then used to identify basis vectors that maximally encode each of the task variables. Using this method, we identified one axis maximally encoding information about the upcoming choice of the animal (“choice axis”), one axis maximally encoding information about the momentary strength of the first stimulus feature (location for rat data, motion for monkey data), and one axis maximally encoding information about the momentary strength of the second stimulus feature (frequency for rat data, color for monkey data). To study how neural dynamics evolved in this reduced space, we first averaged the activity of each neuron across all correct trials according to the strength of location evidence, strength of frequency evidence (i.e., within each of the 36 blocks Fig. 1b), and context, and choice. For this analysis, spike counts were computed in 50 ms non-overlapping bins with centers starting at the beginning of the pulse train presentation and ending 50 ms after the end of the pulse train presentation. For any given trial condition, a “pseudo-population” (i.e. including non-simultaneously recorded neurons) was computed for each time point by compiling the responses of all neurons into a single vector. The trajectory of this vector over time was then projected on the retrieved task-relevant axes to evaluate population dynamics (Fig. 1d-g).

#### 6.2 Pulse-based TDR analysis of neural population dynamics

To estimate the impact of evidence pulses and other task variables on neural responses, we fit the activity of each recorded unit using a pulse-based linear regression (Fig. 1.1 in Extended Discussion). For each neuron, spike counts were computed in 20-ms non-overlapping bins with centers starting 1 second before the beginning of the pulse train presentation, and ending 700 ms after the end of the stimulus presentation. The activity of neuron i at time t on trial k was described as:

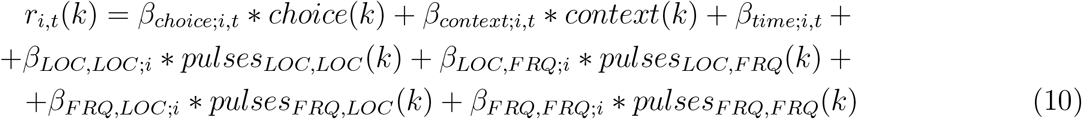

where *x*_*choice*_(*k*) indicates the rat’s choice on trial k (Right = 1, Left = 0), *x*_*c*_*ontext*(*k*) indicates the context on trial k (Location = 1, Frequency = 0), *pulses*_*LOC,LOC*_(*k*) indicates the signed location evidence (number of right pulses minus number of left pulses) presented at each time bin on trial k in the location context, *pulses*_*LOC,FRQ*_(*k*) indicates location evidence in the frequency context, *pulses*_*FRQ,LOC*_ (*k*) indicates frequency evidence (number of high pulses minus number of low pulses) in the location context, and *pulses*_*FRQ,FRQ*_(*k*) indicates frequency evidence in the frequency context. The first three regression coefficients *β*_*choice*;*i*_, *β*_*context*;*i*_ and *β*_*time*;*i*_ account for modulations of neuron i across time according to choice, context and time. The other four sets of regression coefficients *β*_*LOC,LOC*;*i*_, *β*_*LOC,FRQ*;*i*_, *β*_*FRQ,LOC*;*i*_ and *β*_*FRQ,FRQ*;*i*_ indicate the impact of a pulse on the subsequent neural activity, and the symbol ∗ indicates a convolution of each kernel with the pulse train; for example, in the case of location evidence in the location context:

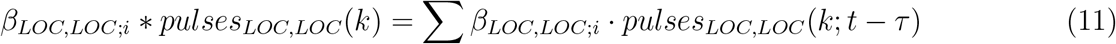

meaning that the element at position *τ* of kernel *β*_*LOC,LOC*;*i*_ represents the impact of a pulse of location evidence in the location context on the activity of unit i after a time. The three kernels for choice, context and time describe modulations from 1 second before stimulus start to 0.7s after stimulus end in 20-ms non-overlapping bins, resulting in 151-dimensional vectors. The four pulse kernels describe modulations from the time of pulse presentation to 0.65s after pulse presentation resulting in 33-dimensional vectors. To avoid overfitting, this regression was regularized using a ridge regularizer, as well as an L2 smoothing prior[8]. Pulse kernels were regarded as an approximation of the neural response to each pulse type (an assumption confirmed by analysis of recurrent neural networks, Fig. 3e,f; Ext. Data Fig. 7c,d).

We wish to emphasize that the critical difference between our previous trial-based application of TDR and the current pulse-based analysis is merely that in the previous trial-based analysis, stimuli are described as two scalar numbers, namely the expected strength of location and frequency evidence over the entirety of a trial. That is, the analysis ignores the precise timing of pulses. In contrast, the pulse-based analysis leverages knowledge of the precise timing of evidence presentation, a feature made possible by the pulse-based nature of our task. Besides that difference in how the stimulus regressors are treated, all other regressors are the same in the two methods; as a consequence, the resulting kernels are very similar across the two methods. This is true in particular for the choice kernels, thus leading to highly similar choice axes using either of the two methods, albeit the kernel-based method is regularized to reduce noise (see the high degree of alignment between the choice axes computed using either method versus the analytically computed line attractor direction in RNNs trained to perform the task, Ext. Data Fig. 7e). Details of the computation of choice axis using the kernel-based are provided in the next section, “Estimating the choice axis”.

Finally, we note that there is a difference between the granularity of the neural kernels (20 ms) and the behavioral kernels (50 ms; see section 5.1). In the case of the neural analysis, we noticed that the initial pulse-triggered response was often very fast, and that a shorter 20ms time bin was best suited to allow us to capture its shape, especially in the first time points after the pulse presentation. In contrast, we noticed that the logistic regression was often noisier, and required pooling over at least 50ms time bins to prevent behavioral kernels from being too noisy. For this reason, we decided to choose the optimal time bin size for each method, rather than using the same time bin for both analyses.

##### 6.2.1 Estimating the choice axis

To compute the population choice axis, we compiled the choice kernels across all neurons, limited to a time window during the presentation of the pulse train stimulus (0 to 1.3s after stimulus start), into a matrix *M*_*c*_ that is *N*_neurons_ x *N*_timebins_ in size. The first principal component of this matrix (i.e., the first eigenvector of 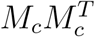, after correcting for the mean firing rate of each neuron), is the *N*_neurons_-long vector in neural space that captures the most variance across choice kernels. This vector was then taken as the choice axis. The pulse-evoked population responses, and their projection onto the choice axis, were computed by compiling pulse kernels across all N neurons recorded from the same rat (Extended Data Fig. 5). At each point in time, the pulse kernel values across all neurons are a vector *N*_neurons_ in length; this was projected onto the choice axis (which is a vector of the same length). We then studied the time-evolution of the results of this projection, which we referred to as the “projection onto the choice axis of population pulse response kernel”.

To test whether the direction of the choice axis was different across the two contexts, we computed the axis for each animal twice, using data collected only from one context at at time (Figure 1e, Ext. Data Fig. 5). To assess whether the direction of the choice axes computed for each context were significantly different from each other, for each rat we performed a random permutation test, where on each iteration we shuffled the context label of each trial. This label-shuffled data becomes the null model. We then recomputed the choice axis separately for trials labeled with each of the two contexts, and measured the angle between the two axes. Done across many shufflings, this provided us with a distribution of the angles between choice axes to be expected from the null model, i.e., if there were no difference across contexts.

##### 6.2.2 Estimating differential neural kernels

To study the differential evolution of pulse-evoked population responses across the two contexts, we computed a “differential pulse response”. For location evidence, the differential pulse response was defined as the difference between the projection onto the choice axis of the response to location pulses in the location context, and the response to location pulses in the frequency context. For frequency evidence, the differential pulse response was computed as the difference between the projection onto the choice axis of the frequency pulse response in the frequency context, minus the frequency pulse response in the location context (Fig. 1.1c in Extended Discussion).

##### 6.2.3 Summarizing the shape of the neural kernels in a “slope index”

To quantify the shape of differential pulse responses, we computed a neural “slope index”. To obtain this, we computed the straight line that provided the least-square fit of the difference between the pulse responses across the two contexts. The slope index was defined as the slope of this fitting line. As a result, a slope index = 0 indicates that the fitting line is perfectly horizontal (i.e. the difference between the two pulse responses is constant at all time points), a slope index *>* 0 indicates a rising differential response, and a slope index *<* 0 indicates a decreasing differential response. Empirically we found that differential pulse responses only displayed positive (or zero) slope indices, i.e. further amplifying the effect of relevant over irrelevant evidence onto the choice axis (Fig. 3h, Extended Data Fig. 8).

### 7 Recurrent neural networks (RNNs)

To validate our analyses of behavior and neural dynamics, and to gather a deeper understanding of the mathematical mechanisms that could underlie our rats’ context-dependent behavior, we trained Recurrent Neural Networks (RNNs) to perform a pulse-based context-dependent evidence accumulation task analogous to that performed by the rats.

The activity of the N=100 hidden units of each network (Ext. Data Fig. 6a) was defined by the dynamical equations:

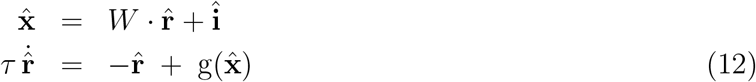

where *τ* is the network time constant, 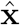 is the vector of activations of each unit, with each of its elements interpreted as roughly paralleling the net input current to a neuron, *W* is the matrix of connections between units, **î** is the external input to each unit, and g() is a pointwise nonlinearity whose output is interpreted as roughly paralleling the activity (i.e., firing rate) of a neuron given that neuron’s net input current. We used g() = tanh(), but similar results should apply with other standard nonlinearities.

The input **î** is in turn composed of several terms:

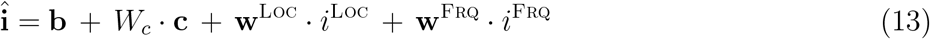

The first term, **b**, represents a bias to each unit that is constant across time and trials. In the second term, **c** is a 2-element-long column vector that encodes current context in a “one-hot” manner (in the Loc context, 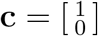, and in the Frq context,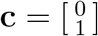). The matrix *W*_*c*_ is *N* x 2 in size, so its first column represents an additive bias to the units in the Loc context, while its second column represents an additive bias in the Frq context. In the next two terms, the time-dependent scalars *i*^Loc^ and *i*^Frq^ represent the momentary Loc and Frq evidence, respectively, with **w**^Loc^ and **w**^Frq^ representing how each of those impact the units of the network.

The output of the network was determined by a single output unit performing a linear readout of the activity of the RNN units:

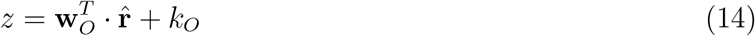

where **w**_*O*_ indicates the N x 1 vector of output weights assigned to each hidden unit, and *k*_*O*_ is a scalar representing the output bias. The choice of the network on a given trial was determined by the sign of *z* at the last time point (T = 1.3 sec). During training and analysis, evolution of the network was computed in 10 ms timesteps. During training, *τ* was set to 10ms, but in subsequent analyses *τ* was set to 100 ms, so as to replicate the autocorrelation timescale observed in neural data.

#### 7.1 Training of RNNs using backpropagation

Recurrent neural networks were trained using back-propagation-through-time with the Adam optimizer and implemented in the Python JAX framework. The weights of the network were initialized using a standard normal distribution, modified according to the number of inputs to a unit, and then rescaled. If *η* is drawn from a standard normal distribution *η N* (0, 1), input weights were chosen as 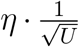; recurrent weights were chosen as 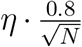 ; output weights were chosen as 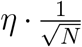; where U indicates the number of inputs (U=4), and N indicates the number of hidden units (N=100). All the biases of the network were initialized at 0. The initial conditions were also learned, and were also initialized randomly from a standard normal distribution, with each element of the initial condition initialized as 0.1. The Adam parameters for training were: b1=0.9; b2=0.999; epsilon=0.1. The learning rate followed an exponential decay with initial step size = 0.002, and decay factor = 0.99998. Training occurred over 120,000 batches with a batch size of 256 trials. Using this procedure, we trained 1000 distinct RNNs to solve the task using different random initializations on each run (Fig. 3a). All networks learned to perform the task with high accuracy (see e.g. Fig. 3c). All the code for training, analysis, and engineering of RNNs will be made available before the time of publication at: https://github.com/Brody-Lab/flexible_decision_making_rnn.

#### 7.2 Analysis of RNN mechanisms

To analyze the linear dynamics implemented by each RNN to perform context-dependent evidence accumulation, we first identified the fixed points of each trained network using a previously described optimization procedure[7, 9]. We then linearized around that fixed point, as follows.

Around any given point 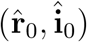, a first-order Taylor expansion tells us that the dynamics (equn. 12) will be approximated by

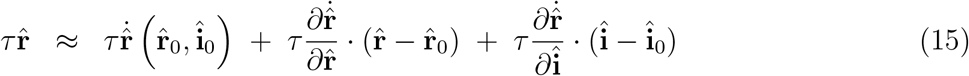

where the partial derivatives are evaluated at 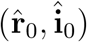. When 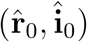 is a fixed point,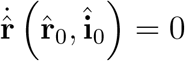. Using equation (12), we can obtain the derivatives

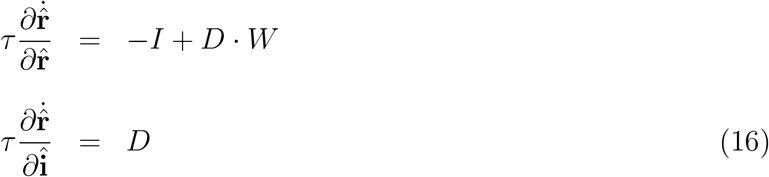

where *D* is a diagonal matrix that we will refer to as “the gain matrix”, and whose elements are given by

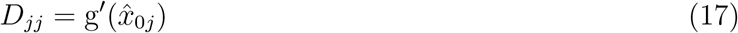

with g^*′*^ being the derivative of the pointwise nonlinearity g() and 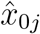 being determined by the fixed point, as they are the elements of 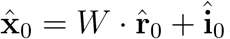.

Combining equation (16) with equation (15), and changing variables to

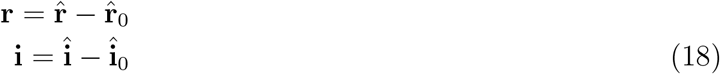

we obtain the linearized dynamics

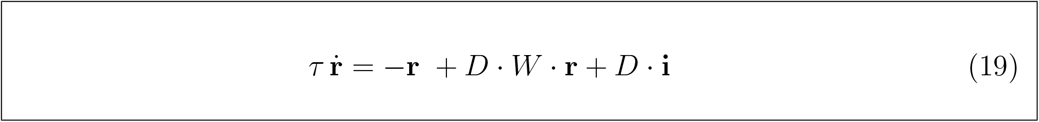

In the absence of sensory evidence, i.e., in the silences between clicks when *i*^Loc^ = 0 and *i*^Frq^ = 0, the fixed points of the system will be determined by **î**_0_ = **b** + *W*_*c*_ *·* **c**. The fixed points are therefore context-dependent, and as a consequence, the gain matrix *D* will also be context-dependent, since it is a function of the fixed point around which we are linearizing (equn. 17). The context-dependence of *D* is what leads to different linearized dynamics in the two contexts.

The linearized connectivity matrix that determines the recurrent dynamics, *D · W*, depends on *D*; and the linearized input vector, *D ·* **i**, also depends on *D*. Thus, this formulation allows both context-dependent modulation of the recurrent dynamics and of the input vector.

(The Extended Discussion describes how RNN equations linearized in the activation space 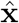, even while equivalent to the dynamics used here, do not allow observing context-dependent input modulation. This would eliminate the right and top corners of the barycentric coordinates of Fig. 2g of the main text. Analyses linearizing in activation space 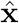 are therefore limited to describing solutions as being 100% selection vector modulation.)

For each trained RNN, we focused on the analysis of the linearized dynamics corresponding to the fixed point with the smallest absolute network output |*z*| (i.e., where the network is closest to the decision boundary), but results were similar when considering different fixed points (i.e., linearized dynamics were mostly similar across different fixed points). Similar to previous reports[7], we found that in every well-trained network, fixed points were roughly aligned to form a “line attractor” for each of the two contexts, and that eigendecomposition of the Jacobian matrix *D · W* reveals a single eigenvalue close to 0, and all other eigenvalues with a negative real value. This reflects the existence of a single stable direction of evidence accumulation (i.e. the line attractor), surrounded by stable dynamics.

The right eigenvector associated with the eigenvalue closest to 0 defined the direction of the line attractor ***ρ***, while the corresponding left eigenvector defined the direction of the selection vector **s**. For each network, we computed these vectors separately for the two contexts by setting the contextual input **c** as 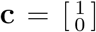 in the Loc context, and 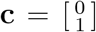) in the Frq context, before computing the fixed points and the eigendecomposition. As a result, for each network we computed the line attractor in each of the two contexts, which we denote as ***ρ***^Loc^ and ***ρ***^Frq^,and the selection vector in each of the two contexts (**s**^Loc^and **s**^Frq^), as well as the linearized input *D ·* **i** in each of the two contexts (**i**^Loc^and **i**^Frq^).Using these quantities, we directly computed the terms in equation 2 of the main text to quantify how much each of the three components contributed to differential pulse accumulation, and we plotted the results for 1000 RNNs in barycentric coordinates (Fig. 3a).

#### 7.3 Engineering of RNNs to implement arbitrary combinations of components

To engineer recurrent neural networks that would implement arbitrary combinations of components, we started from the RNN solutions obtained from standard training using backpropagation-through-time. For a given trained network, we first computed the fixed points of the network and the linearized network dynamics, and we identified the line attractor, selection vector and effective input across the two contexts (see above). Because the RNN dynamics are known (Equations 12 and 13), the linearized dynamics can be expressed in closed form as a function of the network weights:

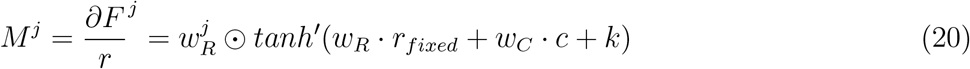

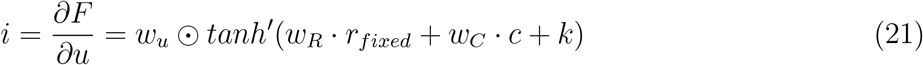

where *M*^*j*^ indicates the j-th column of the jacobian matrix, 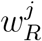 indicates the j-th column of the matrix of recurrent weights, *r*_*fixed*_ indicates the network activity at the fixed point, *tanh*^*′*^ indicates the first derivative of the hyperbolic tangent nonlinearity, and ⊙ indicates the Hadamard product or element-wise multiplication, where the elements of two vectors are multiplied element-by-element to produce a vector of the same size. We further define the “saturation factor” for each of the two contexts as:

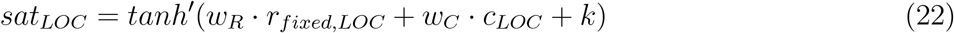

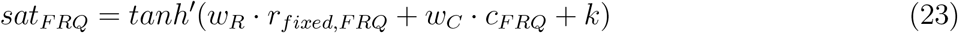

where *r*_*fixed,LOC*_ indicates the fixed point with the smallest absolute network output in the location context, *r*_*fixed,FRQ*_ indicates the fixed point with the smallest absolute network output in the frequency context, *c*_*LOC*_ indicates the context input in the location context (1,0), and *c*_*FRQ*_ indicates the context input in the frequency context (0,1). The effective input for the two contexts can therefore be computed as:

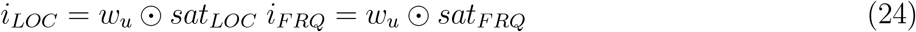

The three components of context-dependent differential integration defined in Equation 2 can therefore be rewritten as a function of the input weights *w*_*u*_. Selection vector modulation, which is equal to the dot product between the difference in the selection vector and the average effective input, can be rewritten as:

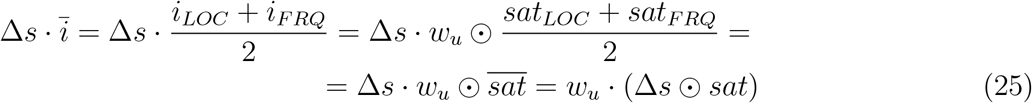

where 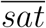 indicates the average saturation factor across contexts, and the last step took advantage of the associative property of the Hadamard and dot product. Direct input modulation, which is equal to the dot product between the difference in the effective input and the line attractor, can be rewritten as:

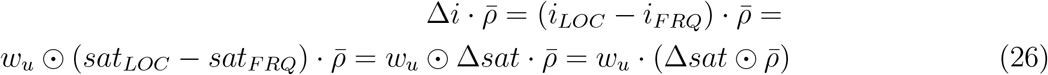

where Δ*sat* indicates the difference between the saturation factor across the two contexts. Indirect input modulation, which is equal to the dot product between the difference in the effective input and the average selection vector orthogonal to the line attractor s, can be rewritten as:

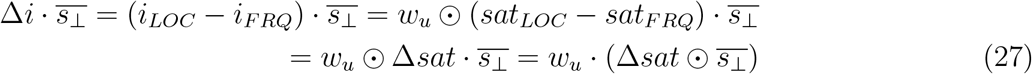

Knowledge of equations 21, 22 and 23 allow us to identify input vectors that produce network dynamics relying on any arbitrary combinations of the three components. For example, producing a network using exclusively selection vector modulation requires the first component (Eq. 21) to be large, while the second (Eq. 22) and third (Eq. 23) components must be 0. In other words, the input weights wu must satisfy:

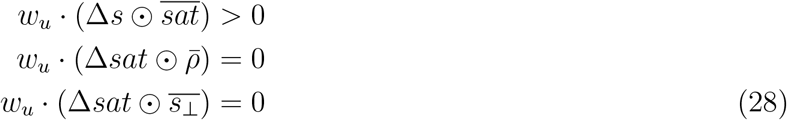

In addition, we must also require that the network does not accumulate the pulse in the irrelevant context. Because we are conducting this analysis for pulses of location evidence, this means that the dot product between the effective input and the selection vector in the frequency context should be 0 :

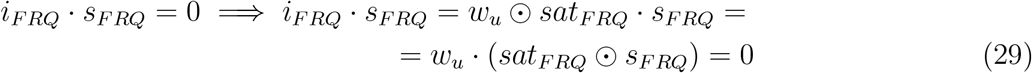

Finally, we then use the Gram-Schmidt process to find the set of weight *w*_*u*_ maximally aligned to the vector 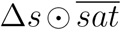, and orthogonal to vectors 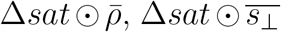 and *sat*_*FRQ*_ ⊙ *s*_*FRQ*_. Similar considerations can be applied to produce networks using different mechanisms. For example, to engineer a network that uses only direct input modulation the input weight must be maximally aligned to 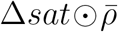 and orthogonal to 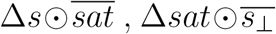 and *sat*_*FRQ*_ ⊙*s*_*FRQ*_. Engineering networks implementing combinations of mechanisms can be obtained by choosing the input vector as a linear combination between extreme network solutions. Finally, we emphasize that the mechanism chosen for one stimulus feature (e.g. location) is entirely independent from the mechanism chosen for the other stimulus feature (e.g. frequency).

### 8 Statistical methods

Comparison of the strength of the encoding of relevant vs irrelevant information (Fig. 1f,g) was performed by quantifying the variability across responses to different stimulus strengths, normalized by trial-by-trial variability, limiting the analysis to the subspace orthogonal to choice encoding. Error bars for neural and behavioral kernels were computed using bootstrapping. On each iteration of the bootstrap procedure we randomly resampled trials, with replacement, and we computed the standard error as the standard deviation of the bootstrapped values over 100 iterations.

## References

1. Okazawa, G. & Kiani, R. Neural Mechanisms that Make Perceptual Decisions Flexible. Annu. Rev. Physiol. (2022) doi:10.1146/annurev-physiol-031722-024731.

2. Livneh, Y. et al. Homeostatic circuits selectively gate food cue responses in insular cortex. Nature 546, 611–616 (2017).

3. Sarel, A. et al. Natural switches in behaviour rapidly modulate hippocampal coding. Nature 609, 119–127 (2022).

4. Mante, V., Sussillo, D., Shenoy, K. V. & Newsome, W. T. Context-dependent computation by recurrent dynamics in prefrontal cortex. Nature 503, 78–84 (2013).

5. Siegel, M., Buschman, T. J. & Miller, E. K. Cortical information flow during flexible sensorimotor decisions. Science 348, 1352–1355 (2015).

6. Sasaki, R. & Uka, T. Dynamic readout of behaviorally relevant signals from area MT during task switching. Neuron 62, 147–157 (2009).

7. Vyas, S., Golub, M. D., Sussillo, D. & Shenoy, K. V. Computation Through Neural Population Dynamics. Annu. Rev. Neurosci. 43, 249–275 (2020).

8. Gold, J. I. & Shadlen, M. N. The neural basis of decision making. Annu. Rev. Neurosci. 30, 535–574 (2007).

9. Brody, C. D. & Hanks, T. D. Neural underpinnings of the evidence accumulator. Curr. Opin. Neurobiol. (2016) doi:10.1016/j.conb.2016.01.003.

10. Brunton, B. W., Botvinick, M. M. & Brody, C. D. Rats and humans can optimally accumulate evidence for decision-making. Science 340, 95–98 (2013).

11. Erlich, J. C., Bialek, M. & Brody, C. D. A cortical substrate for memory-guided orienting in the rat. Neuron 72, 330–343 (2011).

12. Hanks, T. D. et al. Distinct relationships of parietal and prefrontal cortices to evidence accumulation. Nature 520, 220–223 (2015).

13. Leonard, C. M. The prefrontal cortex of the rat. I. Cortical projection of the mediodorsal nucleus. II. Efferent connections. Brain Res. 12, 321–343 (1969).

14. Sinnamon, H. M. & Galer, B. S. Head movements elicited by electrical stimulation of the anteromedial cortex of the rat. Physiol. Behav. 33, 185–190 (1984).

15. Aoi, M. C., Mante, V. & Pillow, J. W. Prefrontal cortex exhibits multidimensional dynamic encoding during decision-making. Nat. Neurosci. 23, 1410–1420 (2020).

16. Okazawa, G., Hatch, C. E., Mancoo, A., Machens, C. K. & Kiani, R. Representational geometry of perceptual decisions in the monkey parietal cortex. Cell 184, 3748–3761.e18 (2021).

17. Charlton, J. A. & Goris, R. L. T. Abstract deliberation by visuomotor neurons in prefrontal cortex. bioRxiv 2022.12.06.519340 (2022) doi:10.1101/2022.12.06.519340.

18. Luo, T. Z. et al. Transitions in dynamical regime and neural mode underlie perceptual decision-making. bioRxiv (2023) doi:10.1101/2023.10.15.562427.

19. Seung, H. S. How the brain keeps the eyes still. Proc. Natl. Acad. Sci. U. S. A. 93, 13339–13344 (1996).

20. Barbosa, J. et al. Early selection of task-relevant features through population gating. Nat. Commun. 14, 6837 (2023).

21. Reynolds, J. H. & Chelazzi, L. Attentional modulation of visual processing. Annu. Rev. Neurosci. 27, 611–647 (2004).

22. Noudoost, B., Chang, M. H., Steinmetz, N. A. & Moore, T. Top-down control of visual attention. Curr. Opin. Neurobiol. 20, 183–190 (2010).

23. Maunsell, J. H. R. & Treue, S. Feature-based attention in visual cortex. Trends Neurosci. 29, 317–322 (2006).

24. Wimmer, R. D. et al. Thalamic control of sensory selection in divided attention. Nature 526, 705–709 (2015).

25. Servan-Schreiber, D., Printz, H. & Cohen, J. D. A network model of catecholamine effects: gain, signal-to-noise ratio, and behavior. Science 249, 892–895 (1990).

26. Pagan, M., Valente, A., Ostojic, S. & Brody, C. D. Brief technical note on linearizing recurrent neural networks (RNNs) before vs after the pointwise nonlinearity. arXiv [cs.LG] (2023).

27. Maheswaranathan, N. & Sussillo, D. How recurrent networks implement contextual processing in sentiment analysis. arXiv [cs.CL] (2020).

28. Park, I. M., Meister, M. L. R., Huk, A. C. & Pillow, J. W. Encoding and decoding in parietal cortex during sensorimotor decision-making. Nat. Neurosci. 17, 1395–1403 (2014).

29. Peixoto, D. et al. Decoding and perturbing decision states in real time. Nature 591, 604–609 (2021).

30. Kurikawa, T., Haga, T., Handa, T., Harukuni, R. & Fukai, T. Neuronal stability in medial frontal cortex sets individual variability in decision-making. Nat. Neurosci. 21, 1764–1773 (2018).

31. Orlandi, J. G., Abdolrahmani, M., Aoki, R., Lyamzin, D. R. & Benucci, A. Distributed context-dependent choice information in mouse posterior cortex. Nat. Commun. 14, 192 (2023).

32. Hopfield, J. J. Neurons with graded response have collective computational properties like those of two-state neurons. Proc. Natl. Acad. Sci. U. S. A. 81, 3088–3092 (1984).

33. Dubreuil, A., Valente, A., Beiran, M., Mastrogiuseppe, F. & Ostojic, S. The role of population structure in computations through neural dynamics. Nat. Neurosci. 25, 783–794 (2022).

34. Langdon, C. & Engel, T. A. Latent circuit inference from heterogeneous neural responses during cognitive tasks. bioRxiv 2022.01.23.477431 (2022) doi:10.1101/2022.01.23.477431.

35. Flesch, T., Juechems, K., Dumbalska, T., Saxe, A. & Summerfield, C. Orthogonal representations for robust context-dependent task performance in brains and neural networks. Neuron 110, 1258–1270.e11 (2022).

36. Flesch, T. et al. Are task representations gated in macaque prefrontal cortex? arXiv [q-bio.NC] (2023).

37. Duan, C. A. et al. Collicular circuits for flexible sensorimotor routing. Nat. Neurosci. 24, 1110–1120 (2021).

38. Perich, M. G. & Rajan, K. Rethinking brain-wide interactions through multi-region ‘network of networks’ models. Curr. Opin. Neurobiol. 65, 146–151 (2020).

39. Orhan, A. E. & Ma, W. J. Publisher Correction: A diverse range of factors affect the nature of neural representations underlying short-term memory. Nat. Neurosci. 22, 505 (2019).

40. Wang, J., Narain, D., Hosseini, E. A. & Jazayeri, M. Flexible timing by temporal scaling of cortical responses. Nat. Neurosci. 21, 102–110 (2018).

41. Sohn, H., Narain, D., Meirhaeghe, N. & Jazayeri, M. Bayesian Computation through Cortical Latent Dynamics. Neuron 103, 934–947.e5 (2019).

42. Remington, E. D., Narain, D., Hosseini, E. A. & Jazayeri, M. Flexible Sensorimotor Computations through Rapid Reconfiguration of Cortical Dynamics. Neuron 98, 1005–1019.e5 (2018).

43. Chadwick, A. et al. Learning shapes cortical dynamics to enhance integration of relevant sensory input. bioRxiv 2021.08.02.454726 (2021) doi:10.1101/2021.08.02.454726.

44. Rodgers, C. C. & DeWeese, M. R. Neural correlates of task switching in prefrontal cortex and primary auditory cortex in a novel stimulus selection task for rodents. Neuron 82, 1157–1170 (2014).

45. Takagi, Y., Hunt, L. T., Woolrich, M. W., Behrens, T. E. & Klein-Flügge, M. C. Adapting non-invasive human recordings along multiple task-axes shows unfolding of spontaneous and over-trained choice. Elife 10, (2021).

46. Barbosa, J., Proville, R., Rodgers, C. C., Ostojic, S. & Boubenec, Y. Flexible selection of task-relevant features through across-area population gating. bioRxiv 2022.07.21.500962 (2022) doi:10.1101/2022.07.21.500962.

47. Li, N., Daie, K., Svoboda, K. & Druckmann, S. Robust neuronal dynamics in premotor cortex during motor planning. Nature 532, 459–464 (2016).

48. Ni, A. M., Ruff, D. A., Alberts, J. J., Symmonds, J. & Cohen, M. R. Learning and attention reveal a general relationship between population activity and behavior. Science 359, 463–465 (2018).

49. Ritz, H. & Shenhav, A. Humans reconfigure target and distractor processing to address distinct task demands. bioRxiv 2021.09.08.459546 (2022) doi:10.1101/2021.09.08.459546.

50. Prinz, A. A., Bucher, D. & Marder, E. Similar network activity from disparate circuit parameters. Nat. Neurosci. 7, 1345–1352 (2004).

51. Pandarinath, C. et al. Inferring single-trial neural population dynamics using sequential auto-encoders. Nature Methods vol. 15 805–815 Preprint at 10.1038/s41592-018-0109-9 (2018).

52. Kim, T. D. et al. Flow-field inference from neural data using deep recurrent networks. bioRxiv 2023.11.14.567136 (2023) doi:10.1101/2023.11.14.567136.

53. Murphy, B. K. & Miller, K. D. Balanced amplification: a new mechanism of selective amplification of neural activity patterns. Neuron 61, 635–648 (2009).

## References

[1] Mikio C Aoi, Valerio Mante, and Jonathan W Pillow. “Prefrontal cortex exhibits multidimensional dynamic encoding during decision-making”. en. In: Nat. Neurosci. 23.11 (Nov. 2020), pp. 1410–1420.

[2] Dmitriy Aronov and David W Tank. “Engagement of neural circuits underlying 2D spatial navigation in a rodent virtual reality system”. en. In: Neuron 84.2 (Oct. 2014), pp. 442–456.

[3] Jennifer Brown et al. “Expanding the Optogenetics Toolkit by Topological Inversion of Rhodopsins”. en. In: Cell 175.4 (Nov. 2018), 1131–1140.e11.

[4] Bingni W Brunton, Matthew M Botvinick, and Carlos D Brody. “Rats and humans can optimally accumulate evidence for decision-making”. In: Science 340.6128 (Apr. 2013), pp. 95–98.

[5] Jason E Chung et al. “A Fully Automated Approach to Spike Sorting”. en. In: Neuron 95.6 (Sept. 2017), 1381–1394.e6.

[6] Timothy D Hanks et al. “Distinct relationships of parietal and prefrontal cortices to evidence accumulation”. In: Nature 520.7546 (Apr. 2015), pp. 220–223.

[7] Valerio Mante et al. “Context-dependent computation by recurrent dynamics in prefrontal cortex”. en. In: Nature 503.7474 (Nov. 2013), pp. 78–84.

[8] Jonathan W Pillow et al. “Spatio-temporal correlations and visual signalling in a complete neuronal population”. en. In: Nature 454.7207 (Aug. 2008), pp. 995–999.

[9] David Sussillo and Omri Barak. “Opening the black box: low-dimensional dynamics in high-dimensional recurrent neural networks”. en. In: Neural Comput. 25.3 (Mar. 2013), pp. 626–649.

[10] Dougal Tervo and Alla Y Karpova. “Rapidly inducible, genetically targeted inactivation of neural and synaptic activity in vivo”. en. In: Curr. Opin. Neurobiol. 17.5 (Oct. 2007), pp. 581– 586.

