## Extended discussion for "A new theoretical framework jointly explains behavioral and neural variability across subjects performing flexible decision-making"

### for

#### Contents

**1 Figures illustrating methods to compute behavioral and neural kernels 5**

**2 The net effect of an input pulse i depends on the interaction between the input and recurrent dynamics, through $\mathbf{s} \cdot \mathbf{i}$ . 5**

**3 Linearizing standard RNNs in firing rate (i.e., activity) space allows observing input modulation, while linearizing in activation space does not. 15**

3.1 Two different linearizations of RNN dynamics. . . . . . . . . . . . . . . . . . 15

3.2 Although they describe the same dynamics, the two linearizations allow different analyses. . . . . . . . . . . . . . . . . . . . . . . . . . . . . . 17

**4 RNNs with rank 1 weight matrices have a 0% selection vector modulation component 18**

**5 Computation through dynamics, line attractors, and non-decision dynamics 19**

**6 Related RNN models of context-dependent evidence accumulation 20**

**7 Expanding the barycentric coordinates framework to the case of line attractors that have different orientations in the two contexts 24**

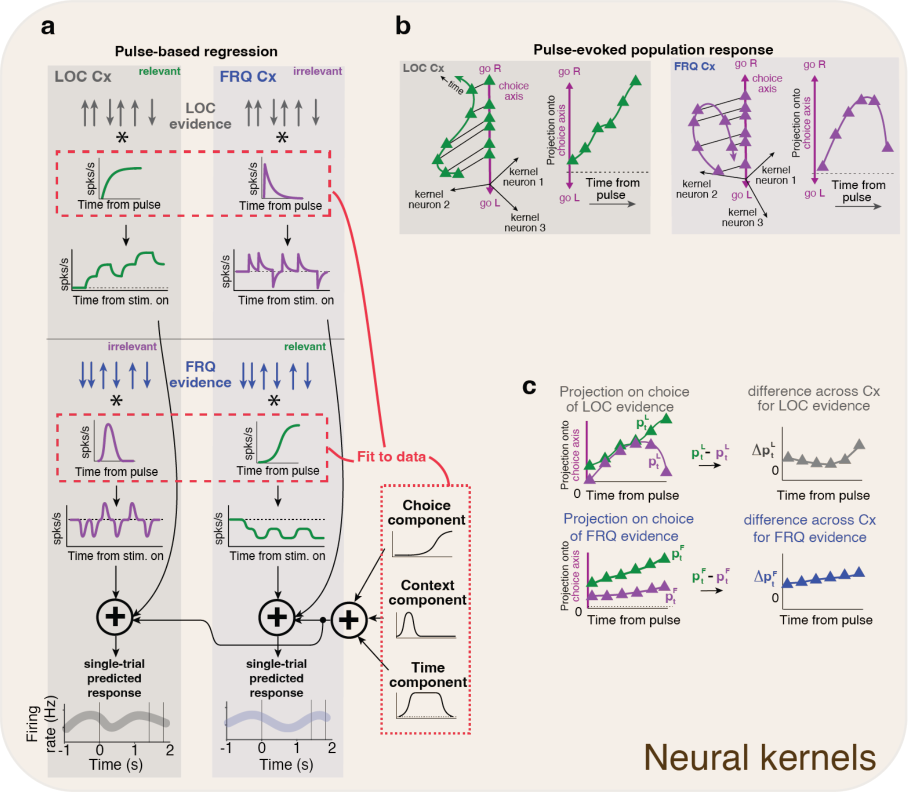

Figure 1.1: **Computation of neural pulse-evoked kernels.** **a)** We first estimate each neuron's pulse-evoked response. To do this, the neuron's firing rate in a trial is modeled as the sum of pulse-response waveforms ("kernels," red dashed rectangles) that are triggered by each of the pulses in the stimulus, plus three time-dependent components (choice, context, and time elapsed during trial, dotted rectangles). Four different pulse-response kernels are fitted to the data, corresponding to LOC and FRQ evidence x LOC and FRQ context. Kernels are different for each neuron, fixed across trials, and fit to data from all trials. The best-fitting pulse-response kernels are the estimates of that neuron's response to each type of evidence pulse in each type of context on average. **b)** For each evidence x context combination, the pulse kernels across many individual neurons recorded from the same rat form a temporal trajectory in neural space. When projected onto the choice axis, this results in an estimate of the population's pulse-evoked dynamics on the choice axis. See Methods for estimation of the choice axis. **c)** Left: pulse-evoked dynamics on choice axis for LOC (top) and for FRQ evidence (bottom), when relevant (green) and irrelevant (magenta). Right: the difference between the relevant and the irrelevant dynamics is the "differential pulse response".

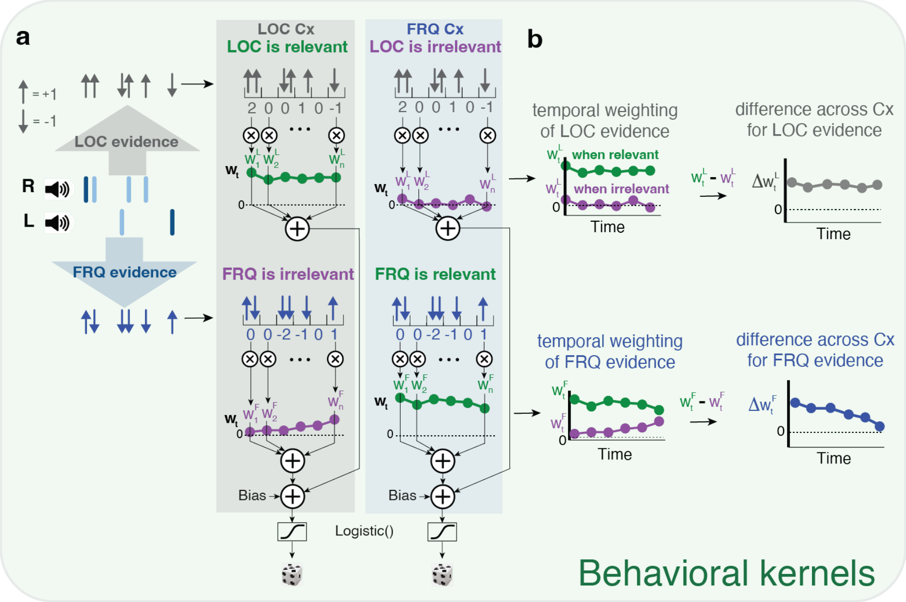

Figure 1.2: **Computation of behavioral pulse-evoked kernels. a)** A given stimulus can be thought of as providing a train of positive and negative pulses of location evidence, as well as a train of pos/neg pulses of frequency evidence. Moreover, a stimulus can be presented in the LOC context (grey background) or in the FRQ context (blue background). A behavioral model of choices in a given context computes a weighted sum of net LOC evidence from each timepoint (50 ms bins) plus a weighted sum of net FRQ evidence from each timepoint, and then passes the net sum through a logistic function, to produce an estimate of the subject's probability of choosing Right. The values of the weights are fit to best match the experimental data, and indicate the strength with which that type of information, at that point in the stimulus, influences decisions. **b)** (Top:) The difference in the LOC weights when LOC evidence is relevant minus when it is irrelevant (green minus purple $= \Delta w_L^T$, the "differential behavioral kernel") is an estimate of how the temporal weighting of LOC evidence differs across the two contexts, i.e., its context-dependence. (Bottom:) Similarly for FRQ evidence.

### 1 Figures illustrating methods to compute behavioral and neural kernels

### 2 The net effect of an input pulse i depends on the interaction between the input and recurrent dynamics, through $\mathbf{s} \cdot \mathbf{i}$ .

#### 2.1 Derivation of $\mathbf{s} \cdot \mathbf{i}$

For completeness, we include a full explanation and definition of the selection vector $\mathbf{s}$, linearization of the dynamics, and a derivation of how, in linearized dynamics with a line attractor, a transient perturbation $\mathbf{i}$ will displace the system along the line attractor by a distance $\mathbf{s} \cdot \mathbf{i}$. This derivation is not specific to RNNs, but applies to any system described by a first-order differential equation of the form in equation (1) below.

The material in this section is drawn from H.S. Seung, *Proc. Natl. Acad. Sci. USA*, 93:23 pp. 13339–13344 (1996), Mante et al., *Nature* 503:7474 pp. 78-84 (2013), and introductory linear algebra textbooks.

---

Let us consider a system with state vector $\hat{\mathbf{r}}$ and inputs $\hat{\mathbf{i}}$, whose dynamics are given by

$$\dot{\hat{\mathbf{r}}} = \mathbf{F}(\hat{\mathbf{r}}, \hat{\mathbf{i}}) \tag{1}$$

and consider one of the fixed points of the dynamics in the absence of inputs, $\hat{\mathbf{r}_0}$, such that

$$\mathbf{0} = \mathbf{F}(\hat{\mathbf{r}}_0, \mathbf{0}) \tag{2}$$

A Taylor series expansion of (1) to first order around $(\hat{\mathbf{r}}_0, \mathbf{0})$ gives

$$\dot{\hat{\mathbf{r}}} \;\approx\; \mathbf{F}(\hat{\mathbf{r}}_0, \mathbf{0}) + \underbrace{\frac{\partial \mathbf{F}}{\partial \hat{\mathbf{r}}}}_{\mathbf{M}} \cdot (\hat{\mathbf{r}} - \hat{\mathbf{r}}_0) + \underbrace{\frac{\partial \mathbf{F}}{\partial \hat{\mathbf{i}}} \cdot \hat{\mathbf{i}}}_{\mathbf{i}} \tag{3}$$

where the derivatives are evaluated at $(\hat{\mathbf{r}}_0, \mathbf{0})$. Letting

$$\begin{aligned} \mathbf{M} &= \left.\frac{\partial \mathbf{F}}{\partial \hat{\mathbf{r}}}\right|_{(\hat{\mathbf{r}}_0,\mathbf{0})} \\ \mathbf{i} &= \left.\frac{\partial \mathbf{F}}{\partial \hat{\mathbf{i}}}\right|_{(\hat{\mathbf{r}}_0,\mathbf{0})} \cdot \hat{\mathbf{i}} \end{aligned} \tag{4}$$

and changing variables to

$$\mathbf{r} = \hat{\mathbf{r}} - \hat{\mathbf{r}}_0 \tag{5}$$

(so the fixed point is at $\mathbf{r} = 0$) we obtain the linearized dynamics

$$\dot{\mathbf{r}} = \mathbf{M} \cdot \mathbf{r} + \mathbf{i} \tag{6}$$

These are the dynamics we will work with.

In the general case, the eigenvectors of $\mathbf{M}$ will not be orthogonal to each other– that is, "normal" dynamics are only a special case. Non-normal dynamics are common. For example, a simple overdamped harmonic oscillator has non-normal dynamics.

**Think of normal dynamics as a rare exception.** Many researchers build intuitions thinking about normal dynamics. We emphasize that in the context of neural networks, normal dynamics are an exceedingly rare special case, and it is dangerous to build one's intuitions based on normal dynamics alone. Normal dynamics follow from symmetric matrices $\mathbf{M}$, which are rarely observed unless there is a constraint that imposes symmetry. For example, consider the probability that a random matrix $\mathbf{M}$ will be exactly symmetric – this probability is zero. Similarly, for a trained RNN, normal dynamics are very unlikely unless a constraint imposing them has been placed on the training (such a constraint is, for example, imposed in Hopfield networks [9]). In the same vein, biological neural networks are not thought to have exactly symmetric connections, and there is no reason to expect normal dynamics.

---

In section 2.1.1 below we will follow standard linear dynamical systems textbook material to re-express general linear dynamics in eigencoordinates; in section 2.1.2 we will draw from [13] and [18] to describe the particular case that concerns us here, linear dynamics with a line attractor; and in section 2.1.3 we focus on desscribing the dynamics and transients seen if the system's state is examined exclusively through the lens of its projection onto the line attractor.

##### 2.1.1 Standard transformation to eigencoordinates

We will now follow textbook methods to express the arbitrary linear dynamics (6) in eigencoordinates. In 2.1.2 we will use this to obtain solutions for the particular case of linear dynamics with a line attractor.

**Transforming to and from eigencoordinates.** First we consider the relationship between original coordinates and eigencoordinates. Any $N$-by-$N$ matrix $\mathbf{M}$ will have $N$ right eigenvectors and eigenvalues such that

$$\mathbf{M}\mathbf{v}_i = \lambda_i \mathbf{v}_i \tag{7}$$

Collecting the eigenvectors $\mathbf{v}_i$ as the columns of a matrix $\mathbf{V}$,

$$\mathbf{V} = [\mathbf{v}_0, \mathbf{v}_1, \mathbf{v}_2, \dots] \tag{8}$$

and the eigenvalues $\lambda_i$ as the diagonal entries in a diagonal matrix $\mathbf{\Lambda}$ (all non-diagonal elements are zeroes) allows writing the above equation for all eigenvector-eigenvalue pairs together,

$$\begin{aligned} \mathbf{MV} &= \mathbf{V\Lambda} \\ \Rightarrow \mathbf{\Lambda} &= \mathbf{V}^{-1}\mathbf{MV} \end{aligned} \tag{9}$$

The first right eigenvector, $\mathbf{v}_0$, will turn out to play a special role below, so we will give it a special symbol, $\boldsymbol{\rho} \equiv \mathbf{v}_0$. We can express any point $\mathbf{r}$ as a linear combination of the eigenvectors $\mathbf{v}_i$:

$$\mathbf{r} = e_0\boldsymbol{\rho} + e_1\mathbf{v}_1 + e_2\mathbf{v}_2 + \dots \tag{10}$$

where $\mathbf{v}_i$ indicates the i$^{\text{th}}$ eigenvector and the column vector $\mathbf{e} = (e_0, e_1, e_2, \dots)^T$ represents the eigencoordinates of $\mathbf{r}$– in other words its coordinates $(r_0, r_1, r_2, \dots)$ transformed to the eigenbasis.

We can rewrite the right hand side of equation (10) as

$$e_0\boldsymbol{\rho} + e_1\mathbf{v}_1 + e_2\mathbf{v}_2 + \dots = \mathbf{V} \cdot \mathbf{e} \tag{11}$$

to obtain

$$\mathbf{r} = \mathbf{Ve}, \tag{12}$$

which tells us how to obtain the eigencoordinates of $\mathbf{r}$,

$$\mathbf{e} = \mathbf{V}^{-1}\mathbf{r}. \tag{13}$$

**Dynamics in eigencoordinates.** Given that $\mathbf{V}$ and $\mathbf{V}^{-1}$ are not functions of time, and using (6), (9), and (12), we can find the dynamics (6), re-expressed in eigencoordinates:

$$\begin{aligned}
\dot{\mathbf{e}} &= \mathbf{V}^{-1}\dot{\mathbf{r}} \\
&= \mathbf{V}^{-1}\mathbf{M}\mathbf{r} + \mathbf{V}^{-1}\mathbf{i} \\
&= \mathbf{V}^{-1}\mathbf{M}\mathbf{V}\mathbf{e} + \mathbf{V}^{-1}\mathbf{i} \\
&= \mathbf{\Lambda}\mathbf{e} + \mathbf{V}^{-1}\mathbf{i}
\end{aligned} \tag{14}$$

Because $\mathbf{\Lambda}$ is diagonal, this tells us that in the eigenbasis $\mathbf{e}$, each of the eigencoordinates evolves independently of the others.

If $\mathbf{i}$ is an instantaneous pulse at $t = 0$, it will place the system at

$$\mathbf{e}(t = 0) = \mathbf{V}^{-1}\mathbf{i}, \tag{15}$$

and thereafter, as each coordinate evolves independently, we have

$$\begin{aligned}
\dot{e}_i &= \lambda_i e_i \\
&\Rightarrow \\
e_i(t) &= e_i(t = 0)e^{\lambda_i t}
\end{aligned} \tag{16}$$

To observe the corresponding effects in neural space, we use (12) to transform back to the original coordinates, namely the neural space basis:

$$\mathbf{r}(t) = \mathbf{V}\,\mathbf{e}(t) = \sum_i \mathbf{v}_i e_i(t) \tag{17}$$

##### 2.1.2 The case of line attractor dynamics

This section is drawn from [13] and [18].

When our dynamics have a line attractor $\boldsymbol{\rho}$, this implies that every point on $\boldsymbol{\rho}$ is a fixed point, so that

$$\mathbf{0} = \mathbf{M} \cdot \boldsymbol{\rho} \tag{18}$$

which we can rewrite as

$$\mathbf{M}\boldsymbol{\rho} = 0\boldsymbol{\rho} \tag{19}$$

In other words, the column vector $\boldsymbol{\rho}$ is a right eigenvector of $\mathbf{M}$, with eigenvalue 0. We will refer to these here as the zero$^{\text{th}}$ eigenvector $\boldsymbol{\rho} = \mathbf{v}_0$ and zero$^{\text{th}}$ eigenvalue $\lambda_0$ of $\mathbf{M}$.

Using (17), and the fact that $\boldsymbol{\rho} \equiv \mathbf{v}_0$, we have

$$\mathbf{r}(t) = \mathbf{v}_0 e_0(t) + \sum_{i>0} \mathbf{v}_i e_i(t) \tag{20}$$

Using (16), and the fact that for a line attractor $\lambda_0 = 0$, we get

$$\begin{aligned} \mathbf{r}(t) &= \boldsymbol{\rho}\, e_0(t=0) e^{\lambda_0 t} + \sum_{i>0} \mathbf{v}_i e_i(t) \\ &= \boldsymbol{\rho}\, e_0(t=0) + \sum_{i>0} \mathbf{v}_i e_i(t) \end{aligned} \tag{21}$$

Following (15), $e_0(t=0)$ will be given by the dot product of the first row of $\mathbf{V}^{-1}$ and $\mathbf{i}$. That first row of $\mathbf{V}^{-1}$ (also known as the zero$^{\text{th}}$ *left* eigenvector of $\mathbf{M}$) is called the *"selection vector"* $\mathbf{s}^T$. We thus obtain

$$\mathbf{r}(t) = \boldsymbol{\rho}\, (\mathbf{s}^T \cdot \mathbf{i}) + \sum_{i>0} \mathbf{v}_i e_i(t) \tag{22}$$

$\boldsymbol{\rho}$ being a line *attractor* means that perturbations off the line attractor decay back to the line attractor. This implies that all other right eigenvectors $\mathbf{v}_i$ of $\mathbf{M}$ have eigenvalues $\lambda_i$ with negative real parts

$$\text{Re}(\lambda_i) < 0 \quad \forall i \geq 1 \tag{23}$$

As a result, as $t \to \infty$, we have $e_i(t) \to 0$ for all $i > 0$. After a sufficiently long time,

$$\mathbf{r}(t \to \infty) = \boldsymbol{\rho}\, (\mathbf{s}^T \cdot \mathbf{i}). \tag{24}$$

In sum, the effect of a input pulse $\mathbf{i}$, after transients have died away, is a displacement along the line attractor $\boldsymbol{\rho}$ of magnitude $|\boldsymbol{\rho}|(\mathbf{s}^T \cdot \mathbf{i})$.

By convention, we normalize $|\boldsymbol{\rho}| = 1$, so that the magnitude of the translation is simply $\mathbf{s}^T \cdot \mathbf{i}$

Notice also that since the zero$^{\text{th}}$ eigencoordinate has eigenvalue $\lambda_0 = 0$, following (16), it remains constant over time:

$$e_0(t) = e_0(t=0)e^{\lambda_0 t} = e_0(t=0) = \text{constant}. \tag{25}$$

Which implies that

$$e_0(t) = \mathbf{s}^T \cdot \mathbf{r}(t) = e_0(t=0) = \mathbf{s}^T \cdot \mathbf{i} \tag{26}$$

Importantly, since $\mathbf{s}^T \cdot \mathbf{r}(t)$ is constant, the flow lines of the dynamics must all be orthogonal to $\mathbf{s}^T$.

This can also be seen by multiplying (6) on the left with $\mathbf{s}^T$, which is a left eigenvector of $\mathbf{M}$ with eigenvalue $\lambda_0 = 0$. In the absence of an external input $\mathbf{i} = 0$, we

$$\begin{aligned} \mathbf{s}^T \cdot \dot{\mathbf{r}} &= \mathbf{s}^T \cdot \mathbf{M} \cdot \mathbf{r} \\ &= \lambda_0 \mathbf{s}^T \cdot \mathbf{r} \\ &= 0 \;\Rightarrow\; \text{flow lines } \dot{\mathbf{r}} \text{ are orthogonal to } \mathbf{s}^{\mathrm{T}}. \end{aligned} \tag{27}$$

It is important to keep in mind that $\mathbf{i}$ is not the input $\hat{\mathbf{i}}$, but the linearized effect of the input $\hat{\mathbf{i}}$ (see equation 4), namely, $\mathbf{i} = \frac{\partial \mathbf{F}}{\partial \hat{\mathbf{i}}} \cdot \hat{\mathbf{i}}$. It is also helpful to keep in mind that $|\mathbf{s}| \neq 1$, but that instead, as we see in the next section, $|\mathbf{s}|$ depends on the angle between $\boldsymbol{\rho}$ and $\mathbf{s}$.

##### 2.1.3 Dynamics and transients projected onto the line attractor

Following (22), and remembering that the line attractor $\boldsymbol{\rho}$ is the zero$^{\text{th}}$ right eigenvector $\mathbf{v}_0$, the projection of $\mathbf{r}$ onto the line attractor is

$$\begin{aligned} \boldsymbol{\rho}^T \cdot \mathbf{r}(t) &= \sum_i e_i(t) \boldsymbol{\rho}^T \mathbf{v}_i \\ &= e_0(t) \mathbf{v}_0^T \mathbf{v}_0 + \sum_{i>0} e_i(t) \mathbf{v}_0^T \mathbf{v}_i \end{aligned} \tag{28}$$

where $e_i(t)$ is the position of the system on the $i^{\text{th}}$ eigencoordinate. Using (26) and remembering that by convention we have normalized all right eigenvectors $\mathbf{v}_i$ to be unit length, we get

$$\boldsymbol{\rho}^T \cdot \mathbf{r}(t) = \mathbf{s}^T \cdot \mathbf{i} + \sum_{i>0} e_i(t) \mathbf{v}_0^T \cdot \mathbf{v}_i \tag{29}$$

As above, for line attractor dynamics $\mathrm{Re}(\lambda_i) < 0$ for all $i > 0$, which implies that the second term (the sum) in (29) describes terms that die away – in other words, it describes transients in the projection of $\mathbf{r}(t)$ onto the line attractor. These transients are due to the decaying eigencoordinates $e_i(t)$ of eigenvectors $\mathbf{v}_i$ that are not orthogonal to $\mathbf{v}_0$, the line attractor[1].

#### 2.2 Constraints between the selection vector s and the line attractor $\boldsymbol{\rho}$

Since $\mathbf{s}$ is the first row of $\mathbf{V}^{-1}$, and $\boldsymbol{\rho}$ is the first column on $\mathbf{V}$, and by definition $\mathbf{V}^{-1} \cdot \mathbf{V} = \mathbf{I}$, the dot product is constrained to be 1:

$$\mathbf{s} \cdot \boldsymbol{\rho} = 1 \tag{30}$$

This means that they cannot be exactly orthogonal (since their dot product is not zero), and the length of the projection of one onto the other must be equal to 1. However, that is their only mutual constraint: a given $\mathbf{s}$ is compatible with many $\boldsymbol{\rho}$, and a given $\boldsymbol{\rho}$ is compatible with many $\mathbf{s}$.

[1] For the (rare, unlikely) case of normal dynamics, $\mathbf{v}_0^T \mathbf{v}_i = 0$ for all $i > 0$ and there are no transients in the projection to the line attractor.

If we follow convention and normalize $|\boldsymbol{\rho}| = 1$, and let $\theta$ be the angle between $\boldsymbol{\rho}$ and $\mathbf{s}$, then

$$\mathbf{s} \cdot \boldsymbol{\rho} = 1 = |\mathbf{s}| \cos\theta \implies |\mathbf{s}| = 1/\cos\theta. \tag{31}$$

The length of $\mathbf{s}$ is equal to the inverse of the cosine of the angle between $\boldsymbol{\rho}$ and $\mathbf{s}$.

#### 2.3 Early gating – examples 1 and 2

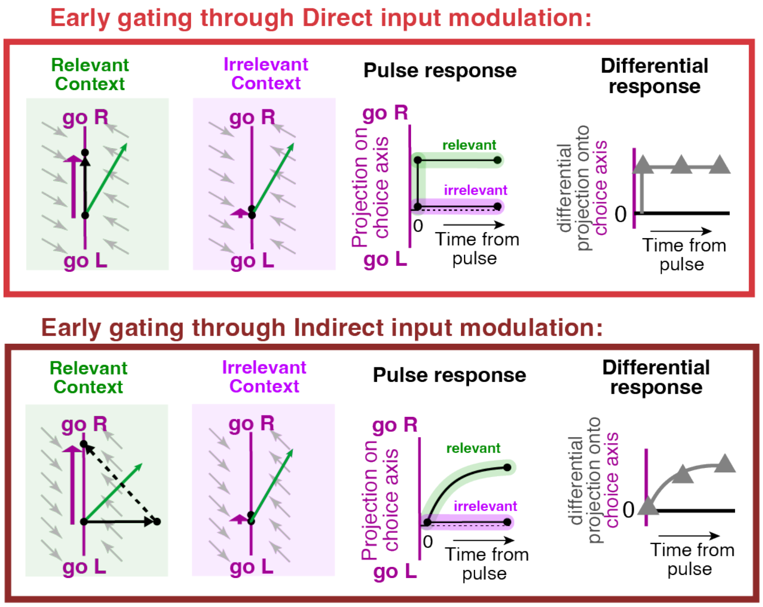

Figure 2.1: Early gating ($|\mathbf{i}| = 0$ in the irrelevant context) is a special case within the theoretical framework of the main text. Format of the panels is as in Fig. 2d–f of the main text: line attractor is in magenta, selection vector is in green, and the input vector is in solid black. Dotted black indicates the relaxation dynamics following an input pulse, and the magenta arrow indicates the net translation along the line attractor, due to an input pulse. **Example 1, Top:** Early gating in which the change in the input vector across the two contexts is entirely parallel to the line attractor (i.e., 100% direct input modulation). Direct input modulation is associated with a fast differential pulse response. **Example 2, Bottom:** early gating in which the change in the input vector is entirely orthogonal to the line attractor (i.e., 100% indirect input modulation), leading to a slow differential pulse response.

Early gating, a model in which irrelevant information is blocked before it even reaches decision-making regions, can be thought of in our terminology as a case where $|\mathbf{i}| = 0$ in the irrelevant context.

Early gating is a special case of the framework of Figure 2 of the main manuscript: it is simply the particular case in which the change in input vectors across contexts, $\Delta\mathbf{i}$, is such that $|\mathbf{i}| = 0$ in one context. As in the rest of the framework, $\Delta\mathbf{i}$ can have components parallel or orthogonal to the line attractor $\boldsymbol{\rho}$.

The examples in Fig. 2.1 show a case in which early gating is 100% direct input modulation, and another case in which it is 100% indirect input modulation.

#### 2.4 Magnitude of the input vector does not alone predict a pulse's impact on choices – example 3

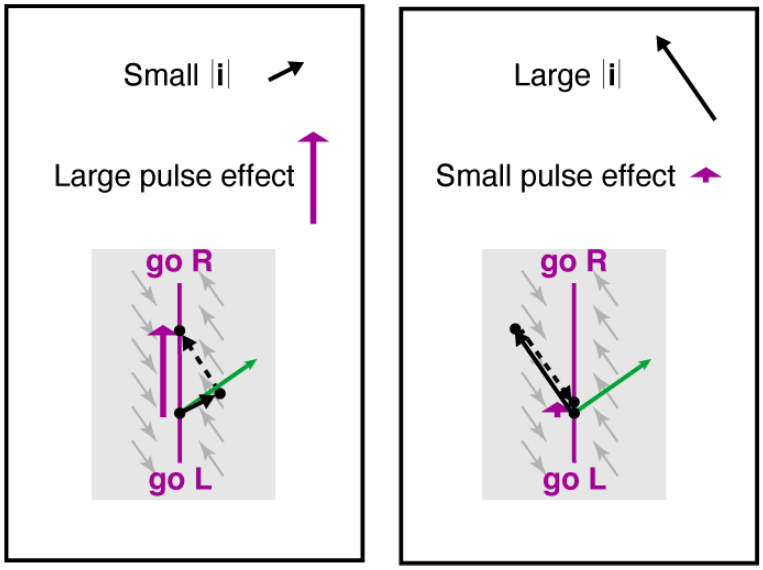

Figure 2.2: **Example 3:** Knowledge of the magnitude of the input vector $|\mathbf{i}|$ is not enough to predict the impact of the pulse on choices. Left, a small input aligned to the selection vector produces a large step forward onto the line attractor. Right, a large input orthogonal to the selection vector produces a small step forward onto the line attractor.

The impact of a pulse of evidence depends on the interaction between the input pulse and the local recurrent dynamics. Because of this, a small magnitude input pulse can be associated with a large effect on choices (Fig. 2.2, left), even while a larger magnitude impulse pulse leads to a smaller effect on choices (Fig. 2.2, right).

#### 2.5 A fixed $\mathbf{s} \cdot \mathbf{i}$ with different $\boldsymbol{\rho}$ – examples 4, 5, and 6

Following section 2.1, knowledge of the relative orientation between the line attractor $\boldsymbol{\rho}$ and the input vector $\mathbf{i}$ is not sufficient to determine the impact of a pulse on choices. Here we illustrate this, showing three examples, in which $\boldsymbol{\rho}$ changes orientation, from almost parallel to $\mathbf{i}$ to almost orthogonal to $\mathbf{i}$, yet the impact of input pulses on choice remains constant across all thee examples.

For all three examples, the input pulse will cause a movement of 0.2 along the line attractor after transients have died away.

**Example 4: $\boldsymbol{\rho}$ almost parallel to i**

$$
\boldsymbol{\rho} = \frac{1}{\sqrt{2}} \begin{bmatrix} 1 \\ 1 \end{bmatrix}
$$
$$
\mathbf{s} = \begin{bmatrix} 1 - 4\sqrt{2} \\ 5\sqrt{2} - 1 \end{bmatrix}
$$
$$
\mathbf{i} = \begin{bmatrix} 1 \\ 0.8 \end{bmatrix} \tag{32}
$$

**Example 5: $\boldsymbol{\rho}$ close to 45° to i**

$$
\boldsymbol{\rho} = \begin{bmatrix} 1 \\ 0 \end{bmatrix}
$$
$$
\mathbf{s} = \begin{bmatrix} 1 \\ -1 \end{bmatrix}
$$
$$
\mathbf{i} = \begin{bmatrix} 1 \\ 0.8 \end{bmatrix} \tag{33}
$$

**Example 6: $\boldsymbol{\rho}$ almost orthogonal to i**

$$
\boldsymbol{\rho} = \frac{1}{\sqrt{2}} \begin{bmatrix} 1 \\ -1 \end{bmatrix}
$$
$$
\mathbf{s} = \frac{1}{9} \begin{bmatrix} 1 + 4\sqrt{2} \\ 1 - 5\sqrt{2} \end{bmatrix}
$$
$$
\mathbf{i} = \begin{bmatrix} 1 \\ 0.8 \end{bmatrix} \tag{34}
$$

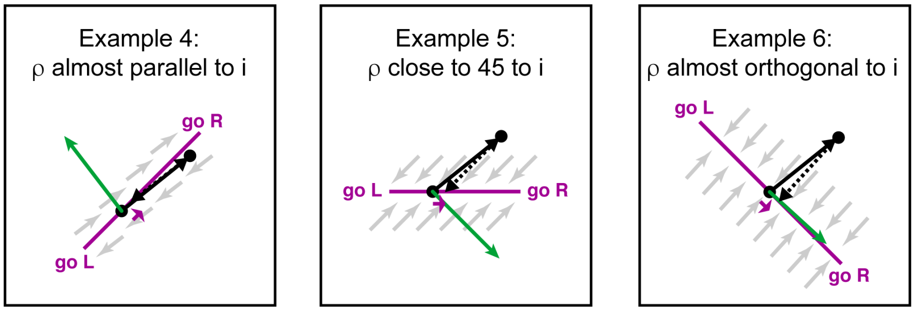

Figure 2.3: In all of these three examples, the final displacement along the line attrractor caused by the input pulse $\mathbf{i}$ is precisely the same, despite widely differing relative orientation between $\mathbf{i}$ and the line attractor $\boldsymbol{\rho}$. **Left: Example 4**, with $\boldsymbol{\rho}$ almost parallel to $\mathbf{i}$. **Center: Example 5**, with $\boldsymbol{\rho}$ close to 45° to $\mathbf{i}$. **Right: Example 6**, with $\boldsymbol{\rho}$ almost orthogonal to $\mathbf{i}$

### 3 Linearizing standard RNNs in firing rate (i.e., activity) space allows observing input modulation, while linearizing in activation space does not.

In this section we will demonstrate that linearizing RNN dynamics in their firing rate space allows observing context-dependent input vector modulation, while linearizing in their activation space does not. When using linearization to study RNN dynamics, then, the choice of linearization is critical, even though the two linearizations describe the same dynamics.

Here we follow [17]; further details are provided there. Although here we use continuous dynamics and [17] uses discrete-time dynamics, the two derivations are almost identical.

#### 3.1 Two different linearizations of RNN dynamics.

Recurrent neural networks (RNNs) are typically modeled with two very closely related sets of equations, which have been shown to be equivalent to each other [15].

The first set of equations, used in our manuscript, is:

$$\begin{aligned} \tau\, \dot{\hat{\mathbf{r}}} &= -\hat{\mathbf{r}} + \mathrm{g}(\hat{\mathbf{x}}) \\ \hat{\mathbf{x}} &= W \cdot \hat{\mathbf{r}} + \hat{\mathbf{i}} \end{aligned} \tag{35}$$

where $\tau$ is the network time constant, $\hat{\mathbf{x}}$ is the vector of activations of each unit, with each of its elements interpreted as roughly paralleling the net input current to a neuron, $W$ is the matrix of connections between units, $\hat{\mathbf{i}}$ is the external input to each unit, and g() is a pointwise nonlinearity whose output is interpreted as roughly paralleling the activity (i.e., firing rate) of a neuron given that neuron's net input current. Thus $\hat{\mathbf{x}}$ is the vector of unit activations, and $\hat{\mathbf{r}}$ is the vector of unit activities.

As described in the methods, linearizing these dynamics around $(\hat{\mathbf{r}}_0, \hat{\mathbf{i}}_0)$ leads to

$$\tau\,\dot{\mathbf{r}} = -\mathbf{r} \;+ D \cdot W\mathbf{r} + D \cdot \mathbf{i} \qquad (36)$$

Here $D$ is a diagonal matrix that we will refer to as "the gain matrix", and whose elements are given by

$$D_{jj} = \mathrm{g}'(\hat{x}_{0j}) \qquad (37)$$

with $\mathrm{g}'$ being the derivative of the pointwise nonlinearity g() and $\hat{x}_{0j}$ being determined by the fixed point, as they are the elements of $\hat{\mathbf{x}}_0 = W \cdot \hat{\mathbf{r}}_0 + \hat{\mathbf{i}}_0$.

Sometimes RNNs are modeled with a very closely related set of equations:

$$\begin{aligned} \tau\,\dot{\hat{\mathbf{x}}} &= -\hat{\mathbf{x}} \;+\; W \cdot \hat{\mathbf{r}} + \hat{\mathbf{i}} \\ \hat{\mathbf{r}} &= \mathrm{g}(\hat{\mathbf{x}}) \end{aligned} \qquad (38)$$

Following the same steps as in the methods to linearize these dynamics, we first compute the derivative of $\dot{\hat{\mathbf{x}}}$ with respect to $\hat{\mathbf{x}}$ (that is, the Jacobian) and the derivative of $\dot{\hat{\mathbf{x}}}$ with respect to the input $\hat{\mathbf{i}}$:

$$\begin{aligned} \frac{\partial \dot{\hat{\mathbf{x}}}}{\partial \hat{\mathbf{x}}} &= -I \;+\; W \cdot D \\ \frac{\partial \dot{\hat{\mathbf{x}}}}{\partial \hat{\mathbf{i}}} &= I \end{aligned} \qquad (39)$$

leads to the linearized dynamics

$$\tau\,\dot{\mathbf{x}} = -\mathbf{x} \;+ W \cdot D\,\mathbf{x} + \mathbf{i} \qquad (40)$$

In the first case (equn. 36), the diagonal gain matrix $D$ multiplies the weight matrix on the *left*, therefore scaling its *rows*, and $D$ also multiplies the input vector $\mathbf{i}$. In contrast, in the second case (equn. 40), the diagonal gain matrix $D$ multiplies the weight matrix on the *right*, therefore scaling its *columns*, and does *not* multiply the input vector $\mathbf{i}$ .

The two linearizations thus have important differences.

#### 3.2 Although they describe the same dynamics, the two linearizations allow different analyses.

In this and other papers that use RNNs to illustrate and study context-dependent decision-making [13, 6, 8, 11], the sensory input to the networks (that is, $\hat{\mathbf{i}}$) is not context-dependent; the challenge is for the network to turn the context-independent inputs into a context-dependent output. To do this, there is an additional, additive input that indicates context to the network: this extra input can the thought of as corresponding to $\hat{\mathbf{u}}_0$. It allows different computations in different contexts, because $\hat{\mathbf{u}}_0$ will set the fixed points of the system, and therefore determine the gain matrix $D$.

**In other words, $D$ is context-dependent.** This is what allows equations (36) and (40) to describe different dynamics in different contexts.

Critically, when the dynamics are linearized in $\hat{\mathbf{x}}$ activation space, as in equation (40), $D$ does not affect the input $\mathbf{i}$. Linearizing activation space dynamics therefore does not allow observing context-dependent modulation of the input in these RNNs.

In contrast, when the dynamics are linearized in $\hat{\mathbf{r}}$ activity space, as in equation (36) and in the main text, $D$ multiplies the external input $\mathbf{i}$, and therefore this linearization *does* allow observing input vector modulation, in the very same RNNs.

The two linearizations describe the same dynamics. But the linearization in firing rate space allows a richer view of the dynamics. In RNNs with constant input vectors, linearization-based analyses will allow observing the full space of solutions (as in the barycentric coordinates of **Fig. 2g** of the main text) only when linearizing in activity space, not activation space.

This important difference was not yet appreciated at the time of [13]. There, linearization in activation space was used, and therefore at that time none of the RNNs could be described as having any input vector modulation component.

It is worth noting that despite their differences, and the difference we have noted regarding the views of the dynamics that the two linearizations allow, equations (36) and (40) in fact describe the same dynamics, albeit observed in different spaces. This can be seen if we take, from equation (38), $\hat{\mathbf{r}} = \mathrm{g}(\hat{\mathbf{x}})$, and linearize it around $\hat{\mathbf{r}_0} = \mathrm{g}(\hat{\mathbf{x}_0})$:

$$\begin{aligned} \hat{\mathbf{r}} &\approx \mathrm{g}(\hat{\mathbf{x}_0}) + \mathrm{g}'(\hat{\mathbf{x}_0}) \cdot (\hat{\mathbf{x}} - \hat{\mathbf{x}}_0) \\ \Rightarrow \hat{\mathbf{r}} - \hat{\mathbf{r}_0} &= \mathrm{g}'(\hat{\mathbf{x}_0}) \cdot (\hat{\mathbf{x}} - \hat{\mathbf{x}}_0) \\ \Rightarrow \mathbf{r} &= D \cdot \mathbf{x} \end{aligned} \tag{41}$$

Using equation (41), we straightforwardly find that if we multiply equation (40) on the left by $D$, we obtain equation (36), which demonstrates the equivalence of the two types of linearized dynamics.

### 4 RNNs with rank 1 weight matrices have a 0% selection vector modulation component

RNNs that have low-rank weight matrices have received substantial interest (e.g., [18, 1, 14, 3]). This specifically includes studies of context-dependent decision-making, in some of which rank=1 matrices have been used [6, 19]. Here we show that such rank=1 RNNs, when linearized in activity space (so as to allow observing the full space of solutions), cannot display any context-dependent selection vector modulation; only rank 2 or higher weight matrices can display the full space of solutions in the barycentric coordinates of **Fig. 2g** of the main text.

Any weight matrix $W$ that is rank=1 can be expressed as the outer product of two vectors $\boldsymbol{\alpha}$ and $\boldsymbol{\beta}$:

$$W = \boldsymbol{\alpha}\boldsymbol{\beta}^T \tag{42}$$

Following the nomenclature of section 3.2, any given context will determine the diagonal gain matrix $D$. Using the activity space linearization in equation (36), the linearized weight matrix will be $D \cdot W$. Since the diagonal gain matrix $D$, when multiplied on the left, scales the rows of $W$, we find that

$$D \cdot W = \hat{\boldsymbol{\alpha}}\boldsymbol{\beta}^T$$

where

$$\hat{\alpha}_j = D_{jj}\alpha_j \tag{43}$$

Here, $\hat{\boldsymbol{\alpha}}$ will be a right eigenvector of $D \cdot W$, with eigenvalue $(\boldsymbol{\beta}^T \cdot \hat{\boldsymbol{\alpha}})$, since $D \cdot W \cdot \hat{\boldsymbol{\alpha}} = \hat{\boldsymbol{\alpha}}(\boldsymbol{\beta}^T \cdot \hat{\boldsymbol{\alpha}})$. Similarly, $\boldsymbol{\beta}^T$ will be the corresponding left eigenvector, with the same eigenvalue.

When the input vector $\mathbf{i} = 0$, and the eigenvalue $(\boldsymbol{\beta}^T \cdot \hat{\boldsymbol{\alpha}}) = 1$, all points lying along the right eigenvector $\hat{\boldsymbol{\alpha}}$ will be fixed points of the dynamics of equation (36). In other words, $\hat{\boldsymbol{\alpha}}$ is the line attractor, i.e., the choice axis.

$\boldsymbol{\beta}$, as the left eigenvector, will be the corresponding selection vector.

Notice that changing context changes $D$ and therefore changes $\hat{\boldsymbol{\alpha}}$ and also changes the linearized input vector $D \cdot \mathbf{i}$. But it does not change the selection vector $\boldsymbol{\beta}$. Consequently, rank 1 RNNs cannot display context-dependent selection vector modulation when analyzed with an activity space linearization, which is the linearization that allows displaying the full space of solutions.

(In contrast, in the activation space ($\hat{\mathbf{x}}$) linearization of equn. (40), $D$ multiplies the weight matrix on the right, and therefore modulates the selection vector $\boldsymbol{\beta}$. In other words, when seen through the lens of the activation space linearization in equn. (40), RNNs that carry out context-dependent evidence accumulation (or any other context-dependent computation), including rank=1 networks, show 100% selection vector modulation – but this is because that linearization *only* allows observing selection vector modulation. Thus a finding of 100% selection vector modulation under activation space linearization is characteristic of the type of analysis, not of the type of network.)

### 5 Computation through dynamics, line attractors, and non-decision dynamics

In this manuscript we have started from the computation-through-dynamics-inspired hypothesis that gradual accumulation of evidence occurs through line attractor dynamics. Based on this hypothesis we derived a theoretical framework that accounted for, and predicted, variability across individuals, and covariability in their behavioral and neural response characteristics. Our focus, then, was not on demonstrating line attractor dynamics *per se* but on the implications that such dynamics would have, with predictions from them found to be confirmed in the experimental data (Fig. 5 of main text).

Line attractor dynamics can coexist with other dynamics. Indeed, neural dynamics related to decision-making are often analyzed after accounting for time-dependent firing rates that are the same regardless of the subject's decision, i.e., non-decision dynamics [13, 12, 10]. Such non-decision-related, time-dependent firing rates are commonly observed in frontal regions [4], and are thought to perhaps be due to an "urgency" signal that anticipates the end of a behavioral trial; indeed, one study showed that when the time of the end of the trial was unpredictable, the non-decision dynamics were eliminated, while the decision dynamics remained intact [10].

Again following others [13, 12], and with the goal of focusing on the decision dynamics, we accounted for the non-decision dynamics through the time-only regressor in the neural kernel estimates, and by a similar time-only regressor in the TDR analyses (Fig. 1 in main text and [13, 2]). The effect of these regressors and their kernels is to essentially subtract the dynamics they account for. The subsequent analyses are thus specific to the decision dynamics. As also found previously [13, 12] the non-decision time-varying activity in our data (e.g., the dynamics of the "time" kernels in our pulse-based regression; see Fig 1.1) is almost orthogonal to the choice axis (percentage of variance of the time-varying dynamics that lies along the choice axis = 1.43%; angle between choice axis and first time-varying axis = 89.7 degrees). The non-decision dynamics can thus be thought of as lying in the null space of the decision dynamics that we are focused on. Time-varying activity in the null space is compatible with stable dynamics and representations in working memory, such as the line attractor dynamics assumed here [5].

### 6 Related RNN models of context-dependent evidence accumulation

Context-dependent accumulation of evidence for decision-making, as studied here and in [13], has received increased recent interest from the computational modeling community. In this section we will briefly discuss the relationship of our work to that of several studies that examined context-dependent decision-making in artificial neural networks trained to perform the task ([6, 19, 11, 8]). Most of these papers focus on recurrent neural networks (RNNs). The exception is ref. [8], which was mostly interested in feedforward networks, but for reasons explained in section 6.3 below, we will focus our comparison on the RNN section of their study.

#### 6.1 Relationship to Dubreuil et al. 2022 and Valente et al. 2022

Two related papers, [6, 19], investigated the role of populations of units in artificial RNNs trained to perform context-dependent evidence accumulation. These studies had a particular interest in RNNs with low rank connectivity matrices. [6] showed that rank=1 RNNs were sufficient to perform the task, as long as the units in the RNN came from at least two distinct populations (with population identity defined by connectivity). Analyzing these rank=1 networks, they further showed that the context-dependent selection mechanism could be thought of as a form of early gating, where a given context effectively lowered the gain of inputs to the RNN from units mostly tuned to evidence from the other context, thus reducing the irrelevant evidence being accumulated. [19] in addition showed that units in rank=1 RNNs could be trained to approximately mimic the activity of units in full-rank RNNs trained to perform the task.

These results thus seemed to suggest that trained full-rank RNNs, instead of performing the task using selection vector modulation (as originally suggested in [13]), performed the task using a form of early gating, albeit with the gating occurring entirely within the decision-making network.

However, subsequent to those publications, and working together with some of those authors, we gained an understanding of the important nuances in analytic views provided by different linearizations of RNNs [17]. This new understanding helped to arrive at the conclusions described in section 4 of this Extended Discussion: we now know that it is mathematically impossible for rank=1 networks to display selection vector modulation. Moreover, the results in Figure 3 of the main text (which we now understand require the richer view provided by the linearization in firing rate space) show that full-rank RNNs, trained to perform the task, predominantly use selection vector modulation. Thus, even though rank=1 networks can come close to mimicking the activity of units in full-rank RNNs [19], there remain important differences, and in particular, the mechanisms they use to perform the task may be different to those of full-rank RNNs. Rank=1 RNNs are constrained to perform the task using 100% input vector modulation. Therefore, in the barycentric coordinates of Fig. 2g of the main text, all rank=1 networks would lie along the right edge of the triangle (Fig. 6.1, green region).

The right edge of the triangle spans its full vertical extent, which real subjects also span (Figure 3 of main text). Since we have not yet determined experimentally where real subjects are distributed along the horizontal axis of the triangle, this means that although rank=1 RNNs are not consistent with the trained full-rank RNNs of Fig. 3, they are consistent with the current experimental data. Future experimental examination of placement of subjects on the horizontal axis of the barycentric coordinates will determine whether or not rank=1 RNNs are indeed sufficient to explain the mechanisms used by biological brains.

#### 6.2 Relationship to Langdon et al. 2022

[11] investigated whether a low-dimensional artificial RNN could be fit to mimic the activity of units in a high-dimensional artificial RNN trained to perform the context-dependent evidence accumulation task. In addition, their low-dimensional network was very specifically constrained, in a manner that allowed identifying each unit in the low-d network with a particular task variable. The low dimensionality, combined with the identification of individual units with task variables, made their low-dimensional circuit, which they referred to as a "latent circuit", readily interpretable. They found that their latent circuit operated by suppressing irrelevant inputs to the evidence accumulation, i.e., a type of early gating, albeit implemented within the decision-making region. They argued that their latent circuit provides an understanding of how higher-dimensional artifical RNNs perform the task.

As with [6, 19], then, these results seemed to suggest that trained full-rank high-dimensional RNNs, instead of performing the task using selection vector modulation (as originally suggested in [13]), performed the task using a form of early gating, albeit with the gating occurring entirely within the decision-making network.

However, the low-dimensional networks of [11] were very specifically and strongly constrained. They had exactly 8 units, and each unit was identified, in a one-to-one fashion with task and output variables, as follows:

- Two units represented the output of the network (one unit for a "Left" decision, and one unit for a "Right" decision), with the final output being read out as the sign of the difference in the activity of these two units. These two units thus underlay the choice axis.
- Two units were identified with one of the two features (i.e., in the nomenclature of our version of the task, one of Loc or Frq) with evidence for Right in that feature being the input to one of the units, and evidence for Left in the same feature being the input to the other unit.
- A further two units were identified with the other feature, in the same manner as the previous two units.
- The final two units were identified with the two contexts, with the inputs to these two being a one-hot vector, namely $\left[\begin{smallmatrix}1\\0\end{smallmatrix}\right]$ for one context, and $\left[\begin{smallmatrix}0\\1\end{smallmatrix}\right]$ for the other context.

Thus there were strict one-to-one constraints between which latent circuit units received which inputs and which units provided which outputs (see the diagonal matrices $w_{\text{in}}$ and $w_{\text{out}}$ in their Fig. 3a). The strengths of each of these one-to-one constrained inputs were fit through gradient descent, but always within those one-to-one constraints. Whether or not lifting those constraints would lead to different results is not yet entirely clear.

In terms of the analysis framework of equation 2 and Figure 2g of the main text, the idealized latent circuit of [11] (their Fig. 3b, summarizing the essence of their results) is a 100% indirect input modulation network (Fig. 6.1, purple dot), since context modulates the inputs, but not in the direct input to the choice axis units (i.e., the two output units); instead, the context modulated the units that themselves in turn provided input to the choice axis units. This corresponds to indirect input modulation (see ”indirect input modulation” box in our Extended Data Figure 9c).

#### 6.3 Relationship to Flesch et al. 2022

[8] investigated how feedforward artificial neural networks perform context-dependent decision-making, and compared their findings to experimental data from fMRI in humans and to non-human primate (NHP) data from ref. [13]. Their feedforward networks did not accumulate evidence over time, so were best suited to describing tasks where all the available evidence was presented simultaneously on each trial (as was the case in the task used in their fMRI experiments), or describing a particular moment in time in each trial (which they argued corresponded to late in the stimulus period for the NHP data).

The main finding was that if prior to training, feedforward networks were initialized with weights distributed with small variance, the training led the networks to learn representations in which irrelevant input was largely gated out – a form of early gating. The authors found a match between their networks and fMRI data, and argued that their networks also matched the NHP data. Partly in support of this view, they reanalyzed the data from [13], and reported that in contrast to the original analysis of [13], but consistent with their proposal, irrelevant momentary evidence was largely suppressed by the end of the stimulus period. However, later discussion between the authors of [8] and [13] led to clarification and a joint public statement [7] which reasserted that the conclusions of the original analysis of the NHP data [13], not those of the more recent analysis[8], were correct. The conclusions regarding data from their fMRI experiments, in which all available evidence was presented simultaneously in each trial, were not affected by the clarification. Thus it is unclear whether the feedforward network-based proposals of [8] are consistent with experimental data involving accumulation of evidence over time.

Nevertheless, in an effort to bridge towards accumulation of evidence over time, [8] also described an RNN version of their proposal. This can be analyzed using the dynamical systems perspective in the current manuscript. Their RNN implementation (illustrated in Fig. S5H in [8]) has a non-trivial relationship to the framework we have proposed, for it was composed of four completely separate line attractors, each one specializing for a given combination of context and relevant feature evidence sign. Each line attractor was implemented as a single unit with an autapse with self-connection weight=1, thus supporting integration of inputs over time and persistent activity. There were no connections between line attractor units. Feature evidence inputs went directly into these line attractor units. Context inputs in the color context inhibited the two line attractor units corresponding to motion, while in the motion context they inhibited the two line attractor units corresponding to color.

The critical aspect that allowed accumulation of feature evidence when that feature was relevant, but not when it was irrelevant, was that these inhibitory context inputs pushed some of the line attractor units below the threshold of their nonlinearity (which was rectified linear). Thus these context inputs were reducing the gain equally, for both how irrelevant inputs directly affected the line attractor unit, as well as of the recurrent dynamics in the line attractor unit. Therefore, in the terminology used here, their effect was 50% direct input modulation and 50% selection vector modulation, corresponding to a point at the center of the left edge of the barycentric coordinates of Fig. 2g in the main text (Fig. 6.1, blue dot).

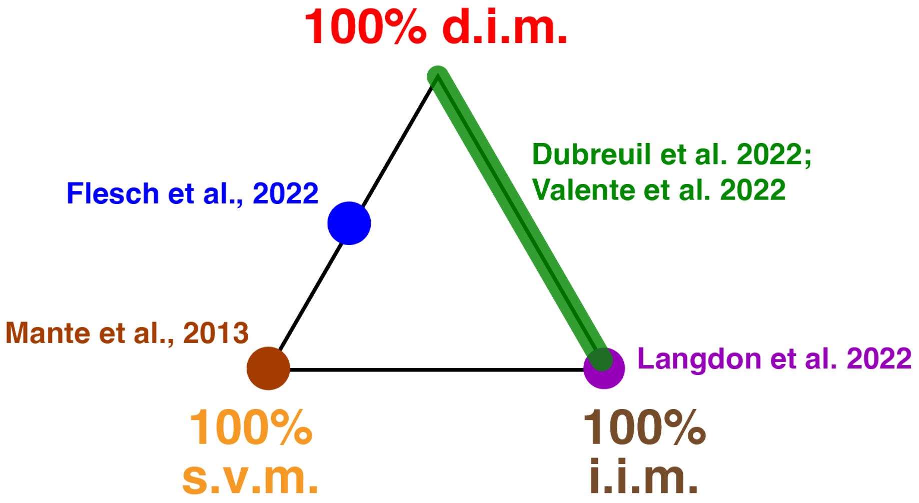

Figure 6.1: **Comparison with other papers.** Recurrent neural network models of context-dependent evidence accumulation proposed in other recent studies can be analyzed within our mathematical framework and occupy distinct positions within our solution space (same as in Fig. 1g in main text). Each colored dot/region indicates a different published network model.

### 7 Expanding the barycentric coordinates framework to the case of line attractors that have different orientations in the two contexts

In the main text, and consistent with our own experimental data and that of [13], we focused on the case where the choice axis (i.e. the line attractor) is parallel across the two contexts. However, it is possible that this result might not hold across other context-dependent decision-making tasks. For example, a recent study [16] reported a rotation of the choice axis measured in parietal cortex as monkeys switched across two different perceptual decision-making tasks. This led us to wonder whether our mathematical framework could be extended to include also the scenario described in [16]. Indeed, we found that it is possible to introduce the case of a rotated readout space as a "fourth solution" in our model. More specifically, we start from the Equation describing the differential integration of a pulse across contexts (Equation 1 in the main text):

$$\Delta(\mathbf{s}\cdot\mathbf{i}) = \underbrace{\bar{\mathbf{s}}\cdot\Delta\mathbf{i}}_{\substack{\text{input}\\ \text{modulation}}} + \underbrace{\Delta\mathbf{s}\cdot\bar{\mathbf{i}}}_{\substack{\text{selection vector}\\ \text{modulation}}} \tag{44}$$

As done in the main text, we now decompose $\Delta\mathbf{i}$ into orthogonal and parallel components, but here we will now define them as orthogonal and parallel to the average line attractor $\hat{\boldsymbol{\rho}} = (\boldsymbol{\rho}^{Loc} + \boldsymbol{\rho}^{Frq})/2$, to obtain

$$\Delta(\mathbf{s}\cdot\mathbf{i}) \quad = \quad \underbrace{\bar{\mathbf{s}}\cdot\Delta\mathbf{i}_{\perp}}_{\substack{\text{indirect input}\\ \text{modulation}}} + \underbrace{\bar{\mathbf{s}}\cdot\Delta\mathbf{i}_{\|}}_{\substack{\text{direct input}\\ \text{modulation}}} + \underbrace{\Delta\mathbf{s}\cdot\bar{\mathbf{i}}}_{\substack{\text{selection vector}\\ \text{modulation}}} \tag{45}$$

For the third term (selection vector modulation term), we will decompose the selection vectors into components parallel and orthogonal to the line attractor for their context. We note that since $\boldsymbol{\rho}\cdot\mathbf{s} = 1$, and by convention we set $|\boldsymbol{\rho}| = 1$, in any given context the component of the selection vector that is parallel to the line attractor is equal to the line attractor vector itself. Thus

$$\begin{aligned} \mathbf{s}^{\text{FRQ}} \quad &= \quad \boldsymbol{\rho}^{\text{FRQ}} + \mathbf{s}_{\perp}^{\text{FRQ}} \\ &\text{and} \\ \mathbf{s}^{\text{LOC}} \quad &= \quad \boldsymbol{\rho}^{\text{LOC}} + \mathbf{s}_{\perp}^{\text{LOC}} \end{aligned} \tag{46}$$

Taking the difference between the two selection vectors, we can decompose the selection vector modulation term in equation 45 into parallel and orthogonal components, to obtain

$$\Delta(\mathbf{s}\cdot\mathbf{i}) \quad = \quad \underbrace{\bar{\mathbf{s}}\cdot\Delta\mathbf{i}_{\perp}}_{\substack{\text{indirect input}\\ \text{modulation}}} + \underbrace{\bar{\mathbf{s}}\cdot\Delta\mathbf{i}_{\|}}_{\substack{\text{direct input}\\ \text{modulation}}} + \underbrace{\Delta\mathbf{s}_{\perp}\cdot\bar{\mathbf{i}}}_{\substack{\text{orthogonal}\\ \text{selection vector}\\ \text{modulation}}} + \underbrace{\Delta\boldsymbol{\rho}\cdot\bar{\mathbf{i}}}_{\substack{\text{line attractor}\\ \text{modulation}}} \tag{47}$$

In equation 47, the first term, $\bar{\mathbf{s}}\cdot\Delta\mathbf{i}_{\perp}$, captures changes in the input orthogonal to the average line attractor. The second term, $\bar{\mathbf{s}}\cdot\Delta\mathbf{i}_{\|}$, captures changes in the input parallel to the average line attractor. The third term, $\Delta\mathbf{s}_\perp \cdot \bar{\mathbf{i}}$, captures changes in the component of the selection vector orthogonal to the line attractor in each context. Because of the dot product with $\bar{\mathbf{i}}$, only changes along the average input direction will be relevant. Finally, the fourth term, $\Delta\boldsymbol{\rho} \cdot \bar{\mathbf{i}}$, captures changes in the line attractor across contexts along the average input direction.

The first three terms correspond to the same terms we describe in Equation 2, with the addition of a fourth term, which captures changes in the direction of the line attractor across contexts. In this more general scenario, any solution to the task can be represented as a combination of these four solutions, i.e. it can be represented as a point inside a tetrahedron (as opposed to the triangle we obtained when assuming that the line attractor is parallel across the two contexts). A graphical intuition for the pulse-evoked dynamics associated with each of these terms is provided in Extended Data Figure 10.

It is worth noting that some of the conclusions we derived in the case of parallel line attractors are no longer valid in this more general case. For example, under the assumption of parallel line attractors, the only solution that can induce an immediate differential pulse response is "direct input modulation", which relies on a context-dependent change of input direction (Fig. 2d-g in main text). However, a change of the line attractor direction across contexts can also generate an immediate differential pulse response even when the input remains constant, because the immediate projections of the same input onto the two line attractors will now in general be different (Extended Data Fig. 10, bottom left quadrant).

### References

[1] Emre Aksay et al. "Functional dissection of circuitry in a neural integrator". en. In: *Nat. Neurosci.* 10.4 (Apr. 2007), pp. 494–504.

[2] Mikio C Aoi, Valerio Mante, and Jonathan W Pillow. "Prefrontal cortex exhibits multidimensional dynamic encoding during decision-making". en. In: *Nat. Neurosci.* 23.11 (Nov. 2020), pp. 1410–1420.

[3] Manuel Beiran et al. "Parametric control of flexible timing through low-dimensional neural manifolds". en. In: *Neuron* (Jan. 2023).

[4] Carlos D Brody et al. "Timing and neural encoding of somatosensory parametric working memory in macaque prefrontal cortex". en. In: *Cereb. Cortex* 13.11 (Nov. 2003), pp. 1196–1207.

[5] Shaul Druckmann and Dmitri B Chklovskii. "Neuronal circuits underlying persistent representations despite time varying activity". en. In: *Curr. Biol.* 22.22 (Nov. 2012), pp. 2095–2103.

[6] Alexis Dubreuil et al. "The role of population structure in computations through neural dynamics". en. In: *Nat. Neurosci.* 25.6 (June 2022), pp. 783–794.

[7] Timo Flesch et al. "Are task representations gated in macaque prefrontal cortex?" In: (June 2023). arXiv: 2306.16733 [q-bio.NC].

[8] Timo Flesch et al. "Orthogonal representations for robust context-dependent task performance in brains and neural networks". en. In: *Neuron* 110.7 (Apr. 2022), 1258–1270.e11.

[9] J J Hopfield. "Neural networks and physical systems with emergent collective computational abilities". en. In: *Proc. Natl. Acad. Sci. U. S. A.* 79.8 (Apr. 1982), pp. 2554–2558.

[10] Hidehiko K Inagaki et al. "Discrete attractor dynamics underlies persistent activity in the frontal cortex". en. In: *Nature* 566.7743 (Feb. 2019), pp. 212–217.

[11] Christopher Langdon and Tatiana A Engel. "Latent circuit inference from heterogeneous neural responses during cognitive tasks". en. Jan. 2022.

[12] Nuo Li et al. "Robust neuronal dynamics in premotor cortex during motor planning". In: *Nature* 532.7600 (Apr. 2016), pp. 459–464.

[13] Valerio Mante et al. "Context-dependent computation by recurrent dynamics in prefrontal cortex". en. In: *Nature* 503.7474 (Nov. 2013), pp. 78–84.

[14] Francesca Mastrogiuseppe and Srdjan Ostojic. "Linking connectivity, dynamics, and computations in low-rank recurrent neural networks". en. In: *Neuron* 99.3 (Aug. 2018), 609–623.e29.

[15] Kenneth D Miller and Francesco Fumarola. "Mathematical equivalence of two common forms of firing rate models of neural networks". en. In: *Neural Comput.* 24.1 (Jan. 2012), pp. 25–31.

[16] Gouki Okazawa et al. "Representational geometry of perceptual decisions in the monkey parietal cortex". en. In: *Cell* 184.14 (July 2021), 3748–3761.e18.

[17] Marino Pagan et al. "Brief technical note on linearizing recurrent neural networks (RNNs) before vs after the pointwise nonlinearity". In: (Sept. 2023). arXiv: 2309.04030 [cs.LG].

[18] H S Seung. "How the brain keeps the eyes still". en. In: *Proc. Natl. Acad. Sci. U. S. A.* 93.23 (Nov. 1996), pp. 13339–13344.

[19] A Valente, Jonathan W Pillow, and S Ostojic. "Extracting computational mechanisms from neural data using low-rank RNNs". In: *Adv. Neural Inf. Process. Syst.* (2022).
